# ToxMCP: Guardrailed, Auditable Agentic Workflows for Computational Toxicology via the Model Context Protocol

**DOI:** 10.64898/2026.02.06.703989

**Authors:** Ivo Djidrovski

## Abstract

Computational toxicology increasingly relies on evidence, high-throughput screening, predictive (Q)SAR, adverse outcome pathways (AOPs), physiologically based kinetic (PBK/PBPK) models, and exposure databases to support integrated approaches to testing and assessment (IATA). Yet the practical workflow remains fragmented across heterogeneous tools, data formats, and licensing regimes. Large language models (LLMs) can lower the interface barrier, but free-text interaction alone is insufficient for regulatory-grade science: it is difficult to audit, difficult to reproduce, and prone to overconfident errors. Here we introduce ToxMCP, a collection of Model Context Protocol (MCP) servers designed as a guardrailed, federated integration layer for reproducible computational toxicology. ToxMCP wraps toxicology-relevant capabilities, including chemical identity and regulatory context (EPA CompTox), rapid ADMET profiling (ADMETlab 3.0), mechanistic pathway retrieval and structuring (AOP knowledge services), quantitative read-across workflows (OECD QSAR Toolbox), and mechanistic PBPK simulation (Open Systems Pharmacology Suite), as typed tools with explicit inputs/outputs, provenance bundles, and policy hooks (e.g., applicability domain checks, critical-action confirmation, and role-based access control). We demonstrate how natural-language risk questions can be compiled into auditable tool invocations, returning mechanistic metrics such as tissue AUC/Cmax, sensitivity curves, and conservative points of departure. We further outline an evaluation protocol for measuring computational reproducibility, task throughput, and scientific utility across multi-tool toxicology tasks. ToxMCP reframes ‘LLMs for toxicology’ from conversational summarizers into accountable orchestrators of established scientific kernels, enabling faster iteration while preserving the evidentiary structure expected in regulatory and academic settings.

**Graphical Abstract:** 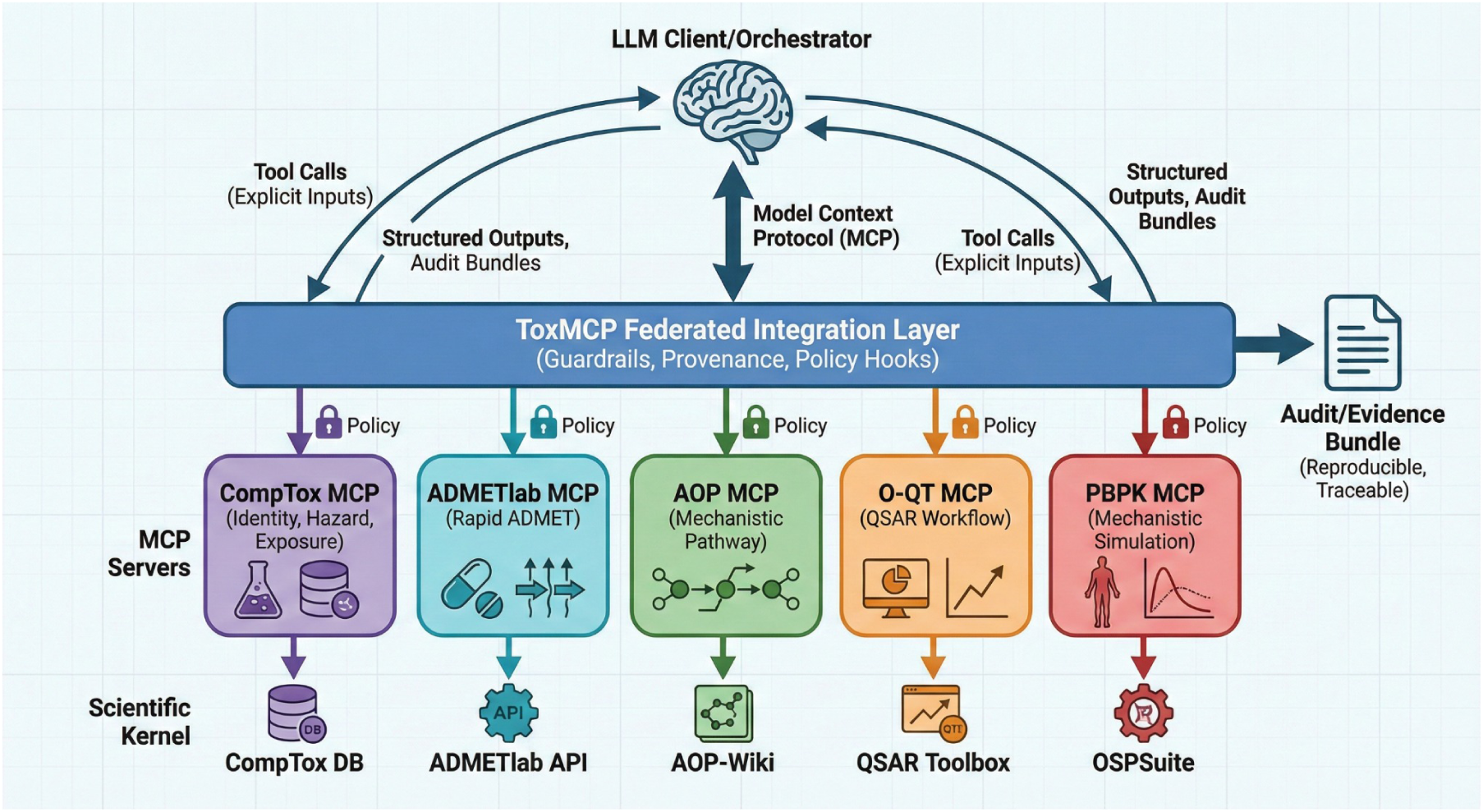

## 1. Introduction

Modern chemical safety assessment increasingly depends on assembling evidence from multiple heterogeneous sources: bioactivity screening (e.g., ToxCast/Tox21)^1,2^, exposure and use patterns, predictive models for ADMET and toxicity endpoints, mechanistic pathway representations such as adverse outcome pathways (AOPs), and physiologically based kinetic (PBK/PBPK) models for in vitro–to–in vivo extrapolation and dose–response interpretation. These components are increasingly combined into integrated approaches to testing and assessment (IATA). In practice, however, the evidence pipeline remains brittle: key datasets live behind separate APIs and dashboards, model outputs are produced by GUI-driven software, and critical metadata (applicability domain, parameter provenance, assumptions) is frequently lost between steps. This fragmentation slows NAMs-driven decision making and increases the risk of irreproducible assessments.^3,4^

Here we introduce ToxMCP, a suite of Model Context Protocol (MCP) servers that expose established computational toxicology capabilities as typed tools with explicit inputs/outputs, provenance metadata, and policy hooks. In its current form, ToxMCP federates chemical identity and regulatory context (EPA CompTox/DSSTox)^5,6^, rapid ADMET profiling (ADMETlab)^7^, mechanistic pathway retrieval and structuring (AOP knowledge services)^8,9^, quantitative read-across workflows (OECD QSAR Toolbox)^10^, and mechanistic PBK simulation (Open Systems Pharmacology Suite)^11^ behind a consistent, auditable tool interface. While the name is toxicology-specific, the underlying pattern (protocol-level tool orchestration with built-in governance) generalises to other evidence-bearing scientific workflows. Figure 1 provides a high-level topology of the federated server suite and the governance signals (policy hooks and provenance) emitted during tool execution.

**Figure 1.**
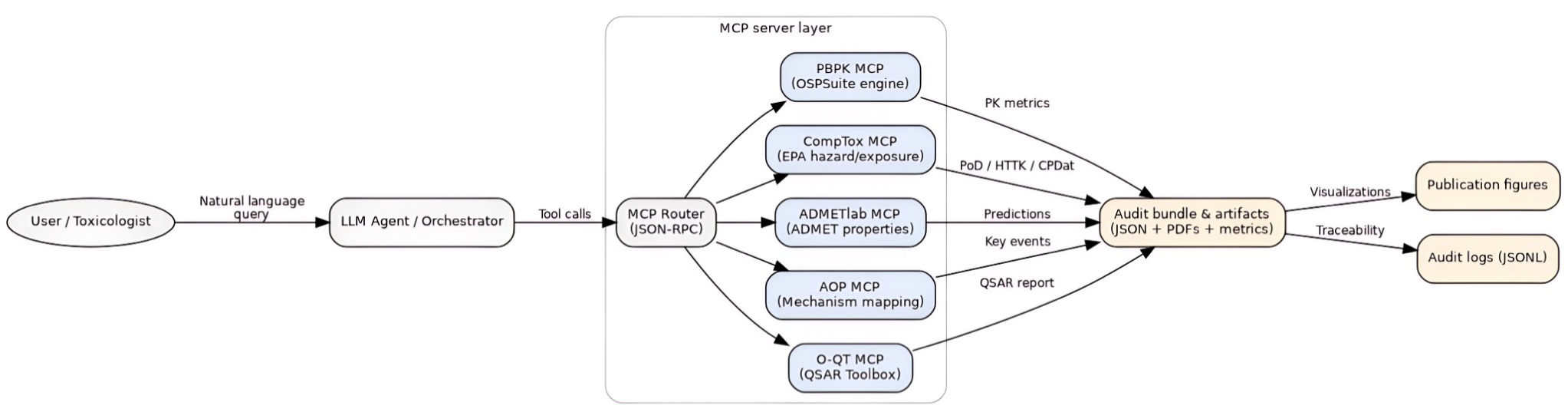
Conceptual topology of ToxMCP. A user-facing MCP client orchestrates multiple domain servers (CompTox, ADMETlab, AOP services, QSAR Toolbox, PBPK). Each server mediates access to its kernel and emits structured outputs with provenance metadata.

Large language models (LLMs) can act as a unifying interface because they translate human intent into tool calls, iterate over intermediate results, and synthesise evidence across heterogeneous resources. Recent agentic demonstrations in chemistry and drug discovery highlight the potential to plan multi-step investigations by combining LLM reasoning with external tools. However, most existing systems rely on ad hoc integrations and free-text reasoning, making it difficult to audit decisions, reproduce results, or enforce safety constraints. Empirical evaluations also show that LLMs can miscalibrate uncertainty and may present overconfident answers, which is particularly problematic for regulatory-grade toxicology.^12,13,14,15^

Integrated decision-support environments are not new in computational toxicology: earlier efforts such as Bioclipse Decision Support and OpenTox demonstrated how interoperable services, standardized representations, and reportable outputs can connect structure-based predictions to evidence and regulatory expectations.^16,17,18^ ToxMCP builds on this lineage, but shifts the integration layer to a protocol-based tool surface designed for modern LLM clients, making guardrails and provenance first-class outputs rather than application-specific features.

We argue that this approach enables a more robust, traceable, and transparent execution of chemical risk-assessment workflows. Throughout, we use “regulatory-grade” to refer to auditable, reproducible computational evidence that can be reviewed; it does not imply regulatory acceptance of any specific model or output. We further argue that the limiting factor is not model capability but infrastructure: a standardised, typed tool interface; guardrails that prevent unsafe or scientifically invalid actions; and provenance that binds each claim to machine-verifiable evidence. MCP provides a portable way to connect an LLM client to tools through structured calls, but it does not by itself define toxicology-ready evidence bundles, applicability-domain checks, or governance-aligned outputs.^19,20^ In the remainder of this manuscript, we describe the ToxMCP architecture and server suite, present exemplar multi-tool workflows, and outline an evaluation protocol for reproducibility and throughput. Our goal is to make toxicology evidence machine-actionable without sacrificing the human requirements of review, traceability, and methodological rigour.

## 2. Conceptual Framework

### 2.1 MCP as a protocol substrate for scientific tooling

The Model Context Protocol (MCP) provides a standard interface for connecting an AI client to external tools and data sources through structured requests and responses. MCP servers advertise callable tools and retrievable resources; clients invoke tools using explicit arguments rather than free-text instructions. This separation (natural language for intent, structured calls for execution) maps naturally onto scientific computing, where correctness depends on explicit parameters, versioned inputs, and deterministic kernels.^19^

For scientific use, the most important property is that the same tool surface can be deployed in different operational settings: locally (next to licensed or sensitive software), within a laboratory or institutional boundary, or as a shared service. This enables practical trust boundaries, authentication, and rate limiting at the server, while keeping the client portable. In ToxMCP, we treat MCP as a protocol substrate; the domain contribution is the toxicology-specific schemas, policy checks, and evidentiary outputs layered on top.^19^

ToxMCP adopts MCP not as a ‘chat plugin’ but as a scientific integration layer: each toxicology capability is expressed as a typed tool with explicit input contracts, structured outputs, and metadata suitable for audit. This design intentionally constrains the LLM. The model can select tools and compose workflows, but identity keys, parameters, and results originate from the underlying kernels and are returned in machine-readable form.

### 2.2 The guardrailed orchestrator pattern

In toxicology, a correct numerical prediction is not automatically a valid prediction. The OECD (Q)SAR Assessment Framework emphasizes that regulatory acceptance depends on transparency, applicability domain, and appropriate interpretation of uncertainty. PBK model guidance similarly emphasizes explicit documentation of assumptions, parameter sources, and validation context. A ‘tool-enabled LLM’ therefore requires more than API connectivity: it requires a policy-aware orchestrator that can enforce minimum scientific constraints.^21,22^

ToxMCP implements each MCP server as a guardrailed orchestrator sitting between the LLM and the underlying scientific system. In practice, tool invocation is treated as a lifecycle with four stages (Figure 2):

1. Identity normalization: ambiguous chemical mentions are resolved to canonical identifiers (e.g., DTXSID, InChIKey, CASRN) to avoid downstream mismatches.
2. Policy verification: before execution, server-side rules evaluate whether the request is permissible and interpretable (e.g., applicability domain checks for QSAR models; confirmation requirements for state-changing actions; rate limits).
3. Execution: the scientific kernel is invoked with explicitly logged inputs.
4. Audit bundling: inputs, policy decisions, outputs, and system metadata are packaged into a provenance object that can be stored or exported as part of a report.^21,23,24^

**Figure 2.**
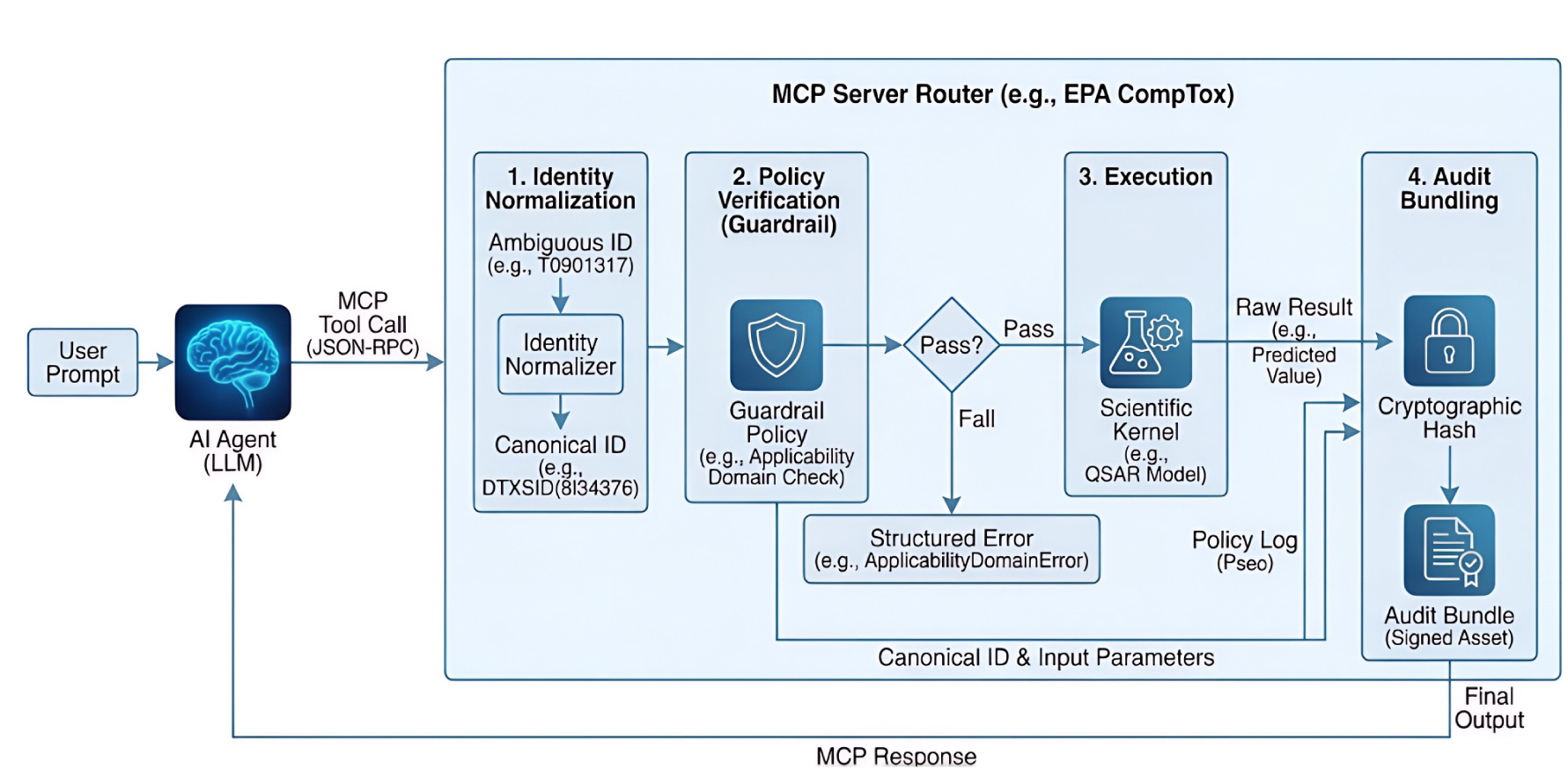
The guardrailed orchestrator workflow. Tool calls are compiled into a four-stage lifecycle: (1) identity normalization to a canonical key, (2) policy verification (applicability and safety checks), (3) deterministic scientific execution, and (4) audit bundling (hashed, signed assets) to support replay and review.

This pattern converts agentic workflows from ‘opaque conversations’ into traceable computational graphs. It also provides a mechanism for aligning automated analysis with community standards. For example, a QSAR tool call can be configured to fail closed outside the model’s applicability domain rather than returning a superficially plausible prediction. Similarly, PBPK simulations can be treated as critical actions requiring explicit user confirmation, preventing unattended compute-heavy or potentially misleading runs (Figure 3).^21,23,24^

**Figure 3.**
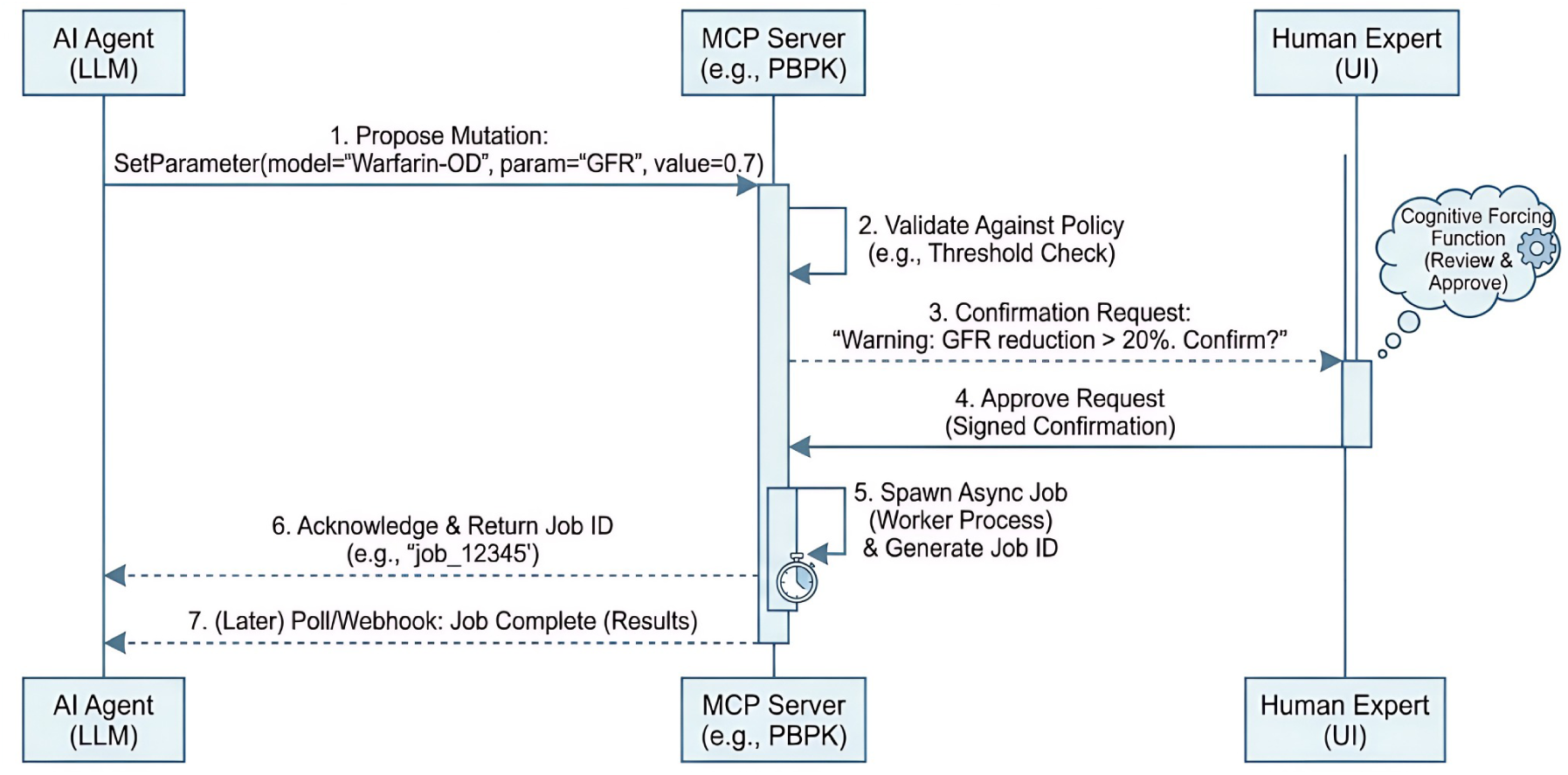
Human-on-the-loop confirmation sequence for critical actions. The MCP server enforces explicit confirmation before executing high-cost or safety-critical operations (e.g., PBPK runs or report exports), then returns a job handle for asynchronous completion and traceable results.

#### Box 1. Governance terms used throughout the manuscript

To help readers from toxicology, software engineering, and regulatory science align terminology, we use the following governance concepts in a consistent way:

| Dimension | Provenance bundles | Evidence bundles | Policy hooks |
| --- | --- | --- | --- |
| Core purpose | Explain how data, models, or results were produced (what ran, with which inputs, and when). | Justify why a claim or decision is supported (what evidence is relevant, sufficient, and how uncertainty is handled). | Enforce what is allowed or required at a control point (what may run, under which constraints). |
| Primary question | “Where did this come from, and how can I replay it?” | “Is this evidence appropriate for the intended interpretation?” | “May this proceed under the rules?” |
| Nature | Descriptive record | Curated argument / safety case | Normative, enforceable constraints |
| Typical contents | Inputs, parameters, tool and dataset versions, timestamps, actors, hashes. | Selected results, rationale, uncertainty notes, links to provenance, citations. | Rules, triggers, required confirmations, access control decisions, refusal reasons. |
| Risk if absent | No traceability or reproducibility. | Decisions lack justification or credibility. | Rules cannot be enforced consistently; higher risk of misuse. |

In ToxMCP, an “audit bundle” is an exportable package that links narrative claims to the underlying provenance and referenced artefacts; “critical actions” are tool calls that require explicit user confirmation (e.g., high-cost simulations or report export).

Additional terminology used here: “transport-agnostic” means the protocol does not depend on a single communication transport (e.g., HTTP vs. stdio is an implementation detail); “scientific kernel” denotes the authoritative execution engine or dataset a server mediates (e.g., PK-Sim/MoBi, QSAR Toolbox, AOP-DB); “cryptographic hash” is a content-derived fingerprint used to identify inputs/outputs; “JSON-RPC” is the structured request/response message format used by MCP; and a “guardrail policy” is the server-side rule set that validates requests (schema, permissions, applicability domain) and records violations.

### 2.3 Asynchronous state for long-running simulations and large outputs

A practical barrier to agentic toxicology is that key computations are not instantaneous. PBPK simulations, population runs, and read-across workflows can exceed typical request timeouts. ToxMCP therefore embraces asynchrony as a first-class design choice. Long-running tools return a job identifier; clients poll job status, retrieve result references, and then request post-processing (e.g., PK metric extraction) as separate tool calls.^11,22^

A second barrier is payload size. Time-series simulation outputs and read-across evidence matrices can be large enough to overwhelm context windows and network transport. ToxMCP uses two complementary strategies: (i) response shaping (e.g., downsampling time series for interactive inspection), and (ii) a ‘claim-check’ pattern in which large objects are stored server-side and returned as references. This allows the LLM to reason over summaries and metrics while preserving access to full-resolution artifacts for later export or review.

### 2.4 Provenance as a first-class output

A core design choice of ToxMCP is that every scientifically meaningful result should be accompanied by sufficient metadata to reconstruct the computation. This includes the identity resolution path, model or database versions, parameter changes, policy checks, and a hash or signature linking these elements. In practice, MCP messages provide a natural envelope for attaching such metadata (e.g., via structured output fields).^25,26^

We refer to these packaged outputs as evidentiary objects: portable bundles that connect a human-readable narrative to machine-verifiable computational provenance. Evidentiary objects enable a workflow where an arXiv preprint, a regulatory dossier, and an internal lab notebook can all refer to the same underlying computation by hash and artifact reference, and, when deposited externally, by a resolvable DOI that points to a versioned bundle. The hash supports byte-level integrity verification, while the DOI supports persistent citation and discovery. Figure 4 shows the structure of a signed audit bundle and its role as an immutable, replayable asset.

**Figure 4.**
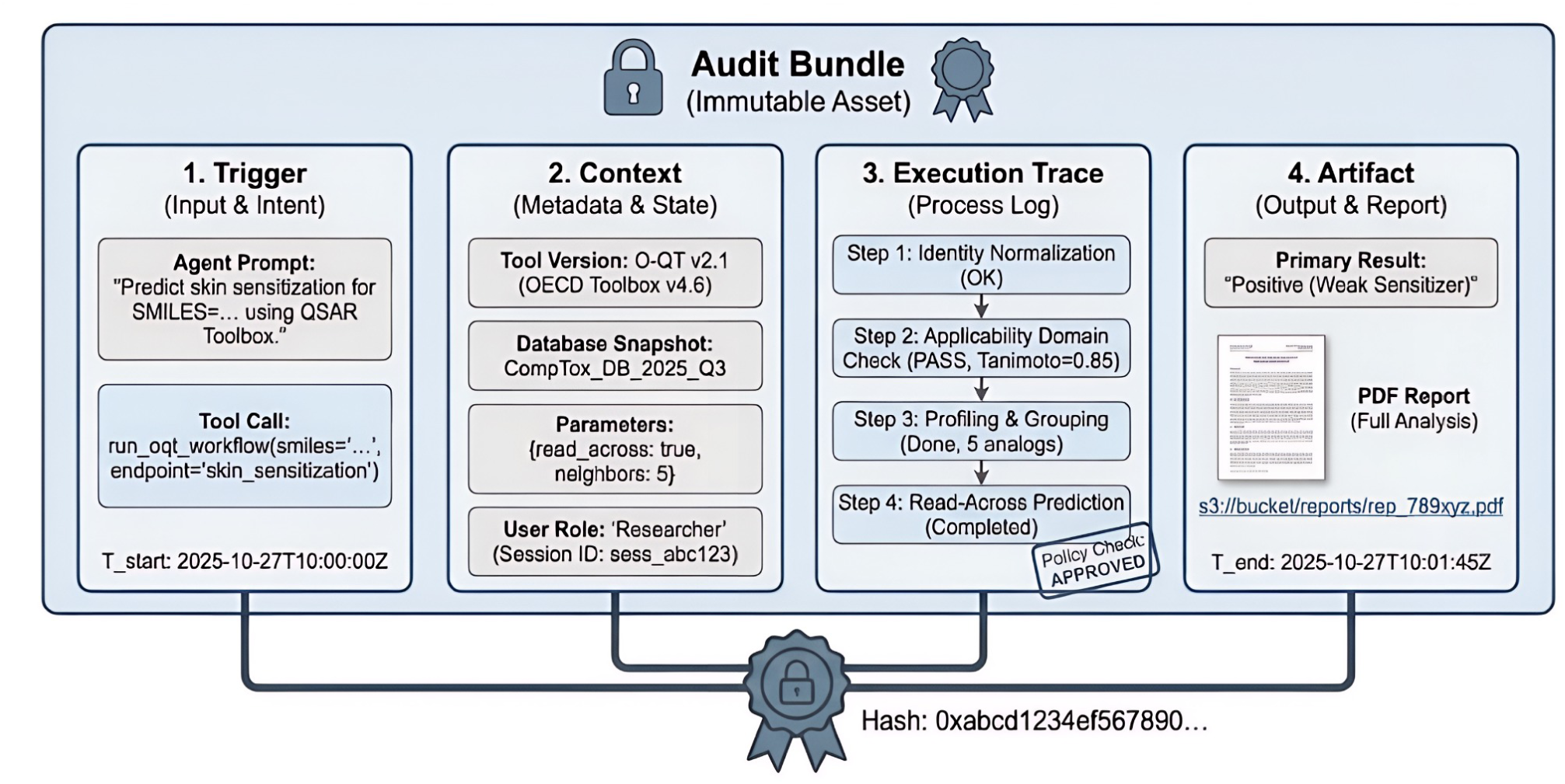
Structure of a cryptographically signed audit bundle. Each workflow emits a self-contained evidence object that captures the trigger (prompt and intent), context (tool versions and dataset snapshots), execution trace, and artifacts (reports and metrics), bound together by a content hash to support replay, peer review, and regulatory audit.

**Figure 5.**
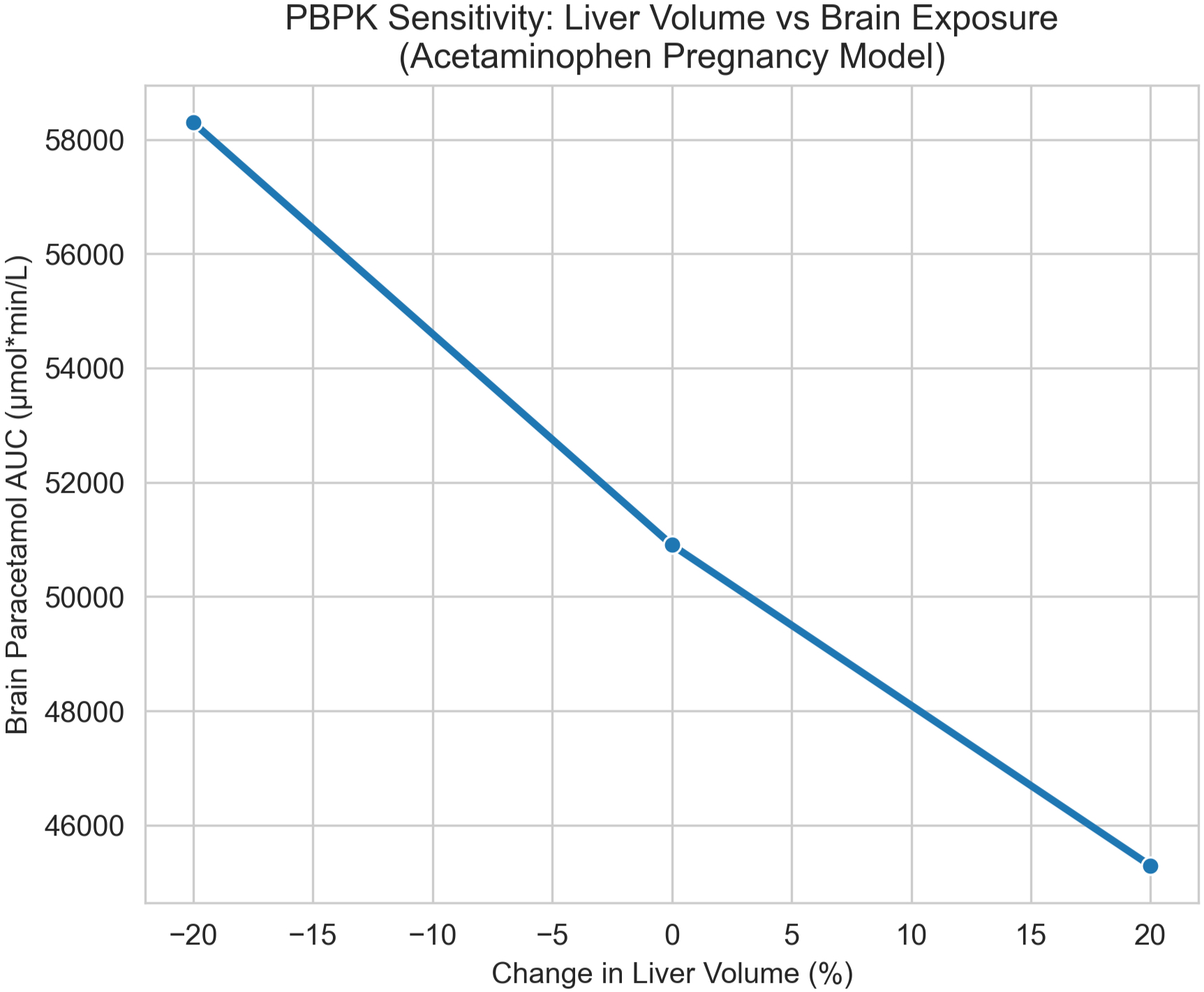
PBPK sensitivity: liver volume versus brain exposure in the acetaminophen pregnancy model. Directional changes in liver volume induce measurable shifts in predicted brain intracellular AUC, illustrating how ToxMCP supports rapid, hypothesis-driven parameter sweeps.

### 2.5 Safety, integrity, and governance threat model

Agentic orchestration introduces failure modes beyond those of individual models or datasets: identity mix-ups (wrong chemical/endpoint), hallucinated values or fabricated citations, tool misuse outside applicability domains, prompt injection via retrieved resources, and unsafe actions executed without appropriate authorization.

ToxMCP addresses these risks by making controls part of the tool interface rather than relying on free-text “best effort” reasoning: (i) typed schemas with strict validation (fail-closed on missing fields), (ii) canonical identifiers (e.g., DTXSID/InChIKey) and cross-tool identity checks, (iii) policy hooks for applicability-domain gates and critical-action confirmation, and (iv) provenance bundles that bind every numeric claim to a tool output and record the execution environment (software versions, model hashes, timestamps).^19,20,25,26^

Operationally, we treat several invariants as non-negotiable for reviewable outputs: (1) every numeric value in a report is traceable to a tool output with a stable artifact identifier; (2) every kernel execution is replayable from captured inputs plus a versioned model/data snapshot; and (3) guardrail-triggered refusals are explicitly logged and surfaced to the user, rather than bypassed. These invariants make it possible to audit both successes and failures, and to distinguish scientific uncertainty from tooling or governance errors.

## 3. Implementation

### 3.1 System overview

ToxMCP is implemented as a federated set of MCP servers, each encapsulating a coherent toxicology capability and its domain-specific constraints. The servers can be deployed independently and combined by any MCP-capable client (e.g., an IDE assistant, a laboratory notebook agent, or a command-line orchestrator). This design avoids a monolithic ‘all-in-one’ agent and instead encourages separation of concerns: each server owns its data sources, rate limits, licensing constraints, and policy checks.

At a high level, a ToxMCP workflow proceeds as follows: the user provides a goal in natural language; the client proposes a plan and selects tools; each tool call executes on a server that validates inputs, enforces safety policies, and returns structured results plus metadata; the client then composes a narrative and (optionally) exports formal artifacts (tables, PDFs, audit bundles). Figure 1 sketches this topology.

Design principle: keep computation close to the authoritative kernel. For example, PBPK simulation runs inside the PBPK server’s environment (where Open Systems Pharmacology binaries and model files are available), rather than being reproduced via approximate surrogate code in the client. Similarly, QSAR Toolbox workflows run inside an O-QT server that can enforce authentication and capture official reports.^10,11,27^

### 3.2 ToxMCP server suite

The current ToxMCP server suite keeps computation close to authoritative kernels while exposing a typed, auditable tool interface. Table 1 summarizes each server, its primary kernel/data source, representative tools, and the governance hooks enforced at the server boundary.

**Table 1.** ToxMCP server suite (abridged): kernels, representative tools, and governance hooks.

| Server | Primary kernel / data | Representative tools | Guardrails & provenance hooks |
| --- | --- | --- | --- |
| PBPK MCP | Open Systems Pharmacology (PK-Sim/MoBi) models; ospsuite execution | load_simulation;<br>list_parameters;<br>set_parameter_value†;<br>run_simulation†;<br>calculate_pk_parameters;<br>run_sensitivity_analysis;<br>run_population_simulation†;<br>export_results | Critical-action confirmation†; RBAC; async jobs + claim-check for large results; audit trail; model hash parity tests |
| EPA CompTox MCP | EPA CompTox (DSSTox identity; CTX APIs; hazard/exposure/bioactivity aggregates) | search_chemical;<br>get_chemical_details;<br>get_exposure_httk;<br>search_hazard;<br>get_hazard_details | Identity normalization; optional applicability-domain policy checks; response schema validation; audit bundle generation; transport hardening |
| ADMETlab MCP | ADMETlab 3.0 web service (in silico ADMET and property prediction) | wash_smiles;<br>wash_batch_smiles;<br>render_svg; predict_admet;<br>predict_admet_batch | Rate limiting; batching; retries/backoff; response normalization; caching for interactive use |
| AOP MCP | AOP knowledge services (AOP-Wiki/AOP-DB; key events; key event relationships) | search_aops; get_aop;<br>list_key_events; list_kers;<br>map_chemical_to_aops;<br>get_evidence_matrix;<br>create_draft_aop | Semantic validation of AOP objects; evidence matrix structuring; provenance for knowledge sources |
| O-QT MCP | OECD QSAR Toolbox workflows (profiling, | profile_endpoint;<br>find_analogues; | OAuth2/OIDC authentication; |
|  | analog selection,<br>read-across, reporting) | run_read_across;<br>generate_pdf_report†;<br>export_matrix | RBAC;<br>auditable report<br>artifacts (PDF +<br>data); strict<br>timeouts and<br>cancellation |

†Denotes tools configured as ‘critical actions’ that require explicit user confirmation in compliant clients.

### 3.3 PBPK MCP: mechanistic PBK simulation as a tool

The PBPK MCP server exposes mechanistic simulation workflows based on the Open Systems Pharmacology Suite (PK-Sim/MoBi) through a minimal set of typed tools. This design deliberately mirrors how toxicologists interact with PBPK models: loading a parameterized model file, inspecting and modifying parameters, running a simulation, and extracting interpretable pharmacokinetic metrics.^11^

Tool surface. The server implements tool calls for (i) model loading, (ii) parameter enumeration and editing, (iii) simulation execution with asynchronous job control, and (iv) post-processing of outputs into PK metrics. Parameter paths use the OSPSuite canonical namespace convention (e.g., “Organism|Age”, “Organism|Brain|Intracellular|Compound|Concentration”), enabling precise specification of the biological quantity under study. AUC/Cmax/Tmax are computed directly from the returned time series using standard numerical integration and peak finding.

PBPK simulations can be computationally expensive and can be misused if parameter edits are applied inadvertently. The PBPK server therefore classifies ‘run’ and ‘edit’ operations as critical actions: in MCP clients that honor the critical-action contract, these tools require explicit human confirmation prior to execution. Additionally, the server supports role-based access control (RBAC) to restrict write actions (e.g., parameter mutation) to authorized users in shared environments.

To keep PBPK results tractable in agentic settings, we implement two patterns. First, long simulations run asynchronously; clients poll job status until completion and then retrieve results by identifier. Second, time-series outputs can be downsampled for interactive inspection, while large multi-subject population results are stored server-side and returned as claim-check references. This supports workflows where an LLM reasons over metrics and summaries but retains the ability to export full-resolution results when needed (e.g., for supplementary data).

To support computational reproducibility, PBPK model execution is validated with an internal parity suite that checks file identity by hash and regression-tests key outputs against expected values within tolerance. This provides a practical bridge between software engineering expectations (continuous integration) and toxicology expectations (traceable model versions).

### 3.4 EPA CompTox MCP: identity, hazard, and exposure as machine-actionable evidence

The EPA CompTox ecosystem provides curated chemical identity (DSSTox), bioactivity screening aggregates, exposure predictors, and toxicity values used in regulatory contexts. Access to these resources is often mediated through dashboards and heterogeneous APIs. The EPA CompTox MCP server wraps these capabilities into a coherent tool catalog centered on three tasks: identity resolution, exposure/toxicokinetic parameter retrieval, and hazard/point-of-departure retrieval.^5,6^

Identity resolution is treated as a mandatory first step. Tools accept flexible queries (common names, CASRN, SMILES, InChIKey) and return canonical identifiers (DTXSID and related keys). Downstream tools then consume these canonical identifiers to retrieve chemical properties, HTTK-derived toxicokinetic predictions (e.g., Css per unit dose, half-life, unbound fraction, volume of distribution), and hazard values (e.g., minimal risk levels, reference doses, or curated toxicological points of departure where available).^6,28^

Guardrails and provenance. The CompTox server is explicitly designed for ‘regulatory-adjacent’ use: it can attach structured metadata describing the source datasets, retrieval time, and any policy filters applied (e.g., selecting high-confidence values). An optional applicability-domain policy layer can flag or block predictions that fall outside the documented scope of a given model. Outputs can be returned with an audit bundle linking query, resolution path, and response, enabling downstream reporting without re-querying the upstream API.

### 3.5 ADMETlab MCP: high-throughput ADMET screening

To support early triage and multi-chemical workflows, ToxMCP includes an ADMETlab MCP server that exposes ADMETlab 3.0 predictions through consistent request/response schemas. While PBPK and QSAR Toolbox workflows offer mechanistic depth, rapid ADMET screening provides breadth: it can prioritize chemicals for which mechanistic modeling is most informative or identify obvious inconsistencies (e.g., predicted permeability incompatible with an assumed target tissue). Supplementary Figure S4 summarizes the ADMETlab MCP server architecture, including client-side guardrails and caching.^7^

The server offers tools for SMILES ‘washing’ and canonicalization, structure rendering (SVG), and ADMET prediction for single molecules or batches. Practical deployment concerns are handled explicitly: requests are rate-limited to avoid upstream throttling, large batches are chunked, and network failures trigger retry/backoff. This turns an interactive web service into a stable component for automated pipelines.

ADMET predictions are returned with feature-level metadata (e.g., prediction type, units, and confidence fields when provided). Downstream, these can be used as priors for PBK parameterization (e.g., permeability-limited distribution), or as endpoints in integrated hazard screening.

### 3.6 AOP MCP: mechanistic structure and evidence matrices

AOP frameworks provide a mechanistic scaffold linking molecular initiating events (MIEs) through key events (KEs) and key event relationships (KERs) to adverse outcomes. In practice, AOP resources are often consulted manually, and the mapping between chemicals, assays, and AOP elements is assembled ad hoc. The AOP MCP server makes this scaffolding programmatically accessible. Supplementary Figure S3 summarizes the AOP MCP server architecture and data-flow to RDF/SPARQL sources.^8,9^

Core tools support searching AOPs, retrieving full AOP objects, listing KEs and KERs, and mapping chemicals to AOP-relevant terms where cross-references exist. Crucially, the server also supports evidence matrices: structured representations of empirical, in vitro, in vivo, and computational evidence supporting each relationship. Evidence matrices serve two functions in agentic workflows: they provide a machine-readable ‘mechanistic justification’ for narrative claims, and they can be exported as supplementary material or regulatory annexes.

Beyond retrieval, the server includes draft-authoring helpers that can assemble an AOP narrative skeleton from structured components. This is not intended to replace expert review; rather, it reduces the friction of producing a structured AOP dossier that can be iteratively refined by domain experts.

### 3.7 O-QT MCP: QSAR Toolbox workflows and report artifacts

The OECD QSAR Toolbox is widely used for chemical profiling, analogue identification, and read-across workflows central to non-animal hazard assessment. However, its use is often constrained by GUI-driven interaction and environment-specific configuration. In our previous work, we developed the O-QT Assistant: an open-source (Apache 2.0) multi-agent LLM pipeline that interprets QSAR Toolbox API outputs and compiles them into a narrative assessment report with a structured JSON log, enabling auditability against the underlying evidence. The O-QT MCP server wraps QSAR Toolbox operations as tools that can be orchestrated by an MCP client while preserving security and audit requirements. Supplementary Figure S5 summarizes the O-QT MCP server architecture and its integration with an external QSAR Toolbox host.^10,21,23,24,27^

The server provides profiling and analogue-search tools, read-across execution, and artifact export. A key design choice is that report generation is treated as a first-class output: the server can emit a PDF report (and the underlying tabular evidence) as an artifact that can be attached to a manuscript or dossier. This makes the agentic workflow compatible with existing review practices, where official QSAR Toolbox outputs are expected.

Because QSAR Toolbox installations may be licensed or may interface with protected endpoints, O-QT includes an authentication layer (OAuth2/OIDC) and role-based access control. This allows institutions to deploy a shared server without exposing credentials to the LLM client, while still enabling automated workflows under controlled policies.

### 3.8 Cross-server orchestration: from ‘chemical question’ to computational evidence

The value of ToxMCP emerges when multiple servers are composed into a single workflow. We highlight a canonical toxicology question: “Does chemical X reach tissue Y at relevant exposure levels, and what is a conservative point of departure for risk characterization?” Addressing this requires, at minimum: identity resolution (CompTox), a hazard value or point of departure (CompTox / curated sources), toxicokinetic parameters or PBK models (CompTox HTTK or PBPK MCP), and post-processing into tissue metrics (PBPK MCP). Optionally, mechanistic explanation can be generated by mapping to AOP elements (AOP MCP) and read-across evidence (O-QT).

Algorithm 1 sketches this orchestration pattern in a client-agnostic way. Importantly, each step yields structured objects that can be cached, audited, and cited, reducing the temptation for the LLM to ‘fill gaps’ with free-text speculation.

#### Algorithm 1.

Hazard-to-kinetics orchestration pattern (high-level).

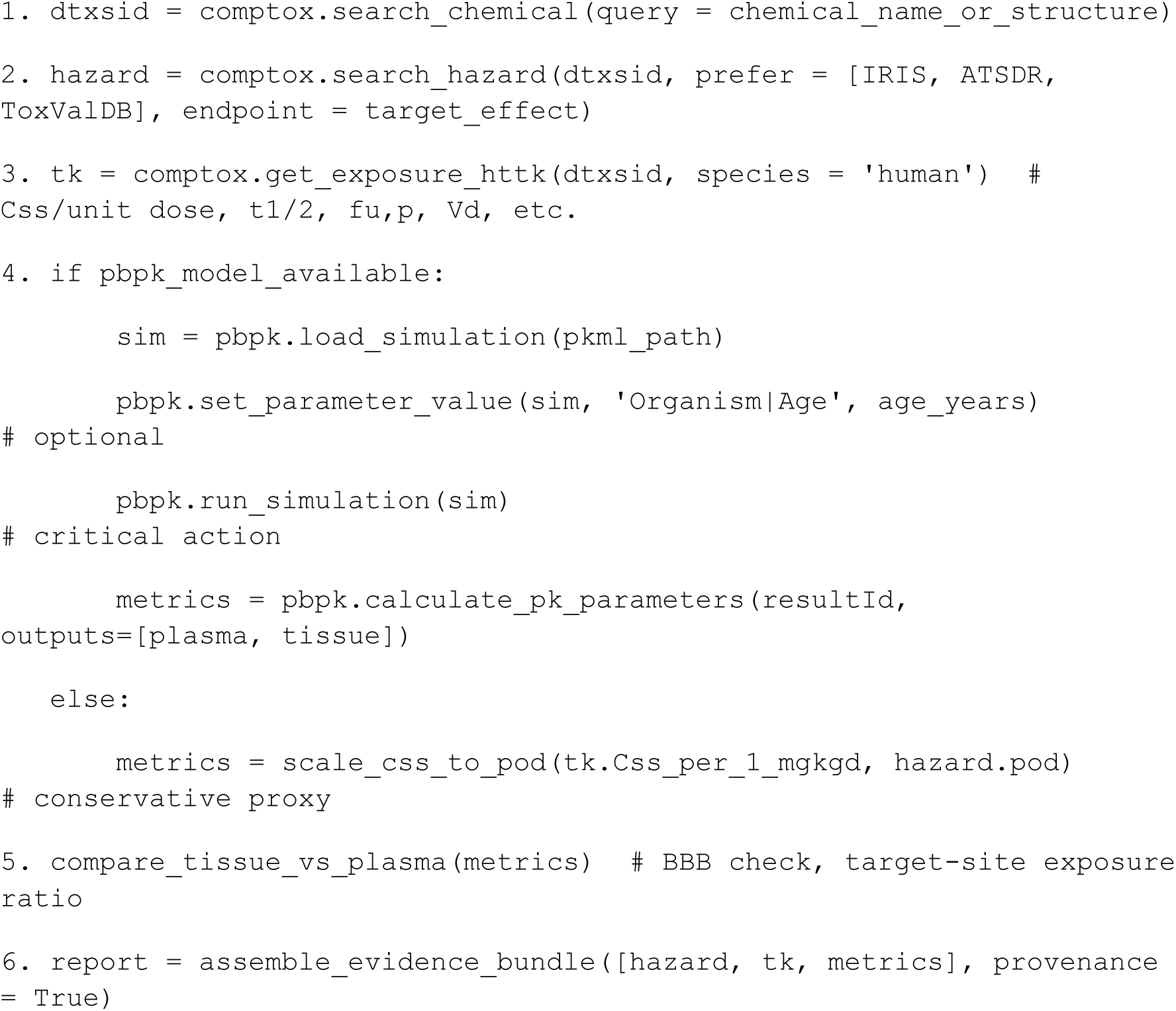

## 4. Results

The following case studies illustrate how toxicology-relevant questions can be executed as reproducible tool workflows. Where numerical values are reported, they originate from executed tool runs and are accompanied by the minimal metadata needed to reproduce the computation (model file, parameter edits, and output variables). Additional case studies can be added by extending the same templates (see Section 5 and Appendix B).

### 4.1 PBPK-derived brain exposure metrics for acetaminophen (model proxy)

We first demonstrate a core capability needed for agentic toxicology: extracting organ-level exposure metrics from a mechanistic PBPK simulation and expressing them as auditable, replayable quantities. Acetaminophen PBPK models for pregnancy are commonly used to characterize maternal kinetics, metabolite formation, and maternal–fetal transfer; in this benchmark we use an available pregnancy-configured model instance as a reproducible template to demonstrate how ToxMCP (i) loads an Open Systems Pharmacology model, (ii) executes a simulation under critical-action confirmation, and (iii) computes PK metrics from explicit output paths.^11,29^

We queried two outputs from the simulation results: brain intracellular paracetamol concentration and peripheral venous plasma paracetamol concentration (parameter paths shown in Table 2). We computed AUC for each variable and report a brain-to-plasma AUC ratio as a compact descriptor of relative tissue exposure. In this context, the ratio is used as a demonstration metric for tissue-versus-plasma comparison and should not be interpreted as a validated BBB permeability estimate without additional model qualification and context. Future benchmarks will add external qualification checks by comparing selected PBPK outputs against published validation datasets where available, and by explicitly documenting model domain and assumptions.

**Table 2.** Acetaminophen proxy BBB analysis (simulation run: acetaminophen_pregnancy_1764095178).

| Output | AUC (μmol·h/L) | Notes |
| --- | --- | --- |
| Brain intracellular paracetamol concentration | 50,903.84 | Organism\|Brain\|Intracellular\|Paracetamol\|Concentration |
| Arterial blood plasma paracetamol concentration | 52,036.25 | Organism\|ArterialBlood\|Plasma\|Paracetamol\|Concentration |
| Brain/Blood AUC ratio | 0.978 | BBB heuristic: ratio > 0.50 ⇒ ‘crosses’ |

Result interpretation: brain intracellular AUC was 50,903.84 μmol·h/L and arterial blood plasma AUC was 52,036.25 μmol·h/L, yielding a brain/plasma AUC ratio of 0.978. Within this model instance, brain intracellular exposure closely tracks plasma exposure. Crucially, the associated evidence bundle records the model identity (hash), output paths, and metric extraction trace, enabling deterministic replay and audit.

**Table 3.** Effect of liver volume on brain exposure (brain intracellular paracetamol AUC).

| Scenario | Organism\|Liver\|Volume | Brain AUC<br>($\mu\text{mol}\cdot\text{h/L}$) | % change vs<br>baseline |
| --- | --- | --- | --- |
| Baseline | 1.5 | 59,058.37 | 0% |
| Severe impairment | 0.75 | 84,501.28 | +43.1% |
| Hypertrophy | 3.0 | 37,863.93 | -35.9% |

**Table 4.** Brain intracellular Cmax across three ages (acetaminophen proxy model).

| Subject | Age (years) | Brain Cmax ($\mu\text{mol/L}$) |
| --- | --- | --- |
| Subject A | 20 | 160.61 |
| Subject B | 40 | 160.61 |
| Subject C | 60 | 160.61 |

Population mean brain Cmax across these three subjects was 160.61 $\mu\text{mol/L}$ .

**Table 5.** Chlorpyrifos: HTTK parameters and PoD-scaled plasma exposure proxy.

| Quantity | Value | Notes |
| --- | --- | --- |
| Identity | DTXSID4020458; CAS 2921-88-2 | Resolved via CompTox identity search |
| HTTK Css per 1 mg/kg/day (median) | 16.08 mg/L | Human prediction |
| HTTK Css per 1 mg/kg/day (95th) | 99.1 mg/L | Human prediction (variability) |
| HTTK half-life | 488 h | Human prediction |
| HTTK Vd | 2.8 L/kg | Human prediction |
| HTTK fu,p | 0.0079 | Fraction unbound in plasma |
| PoD proxy (ATSDR chronic oral MRL) | 0.001 mg/kg/day | Health-based guidance value |
| PoD-scaled plasma C<sub>ss</sub> (median) | 0.0161 mg/L | Linear scaling from 1 mg/kg/day |
| PoD-scaled unbound plasma C<sub>ss</sub> (median) | 0.000127 mg/L (0.127 µg/L) | C<sub>ss</sub> × f<sub>u,p</sub> |
| PoD-scaled plasma C<sub>ss</sub> (95th) | 0.0991 mg/L | Inter-individual variability proxy |
| PoD-scaled unbound plasma C<sub>ss</sub> (95th) | 0.000783 mg/L (0.783 µg/L) | C<sub>ss</sub> × f<sub>u,p</sub> |

Interpretation: chlorpyrifos is predicted to have a large apparent volume of distribution and a low unbound fraction in plasma, suggesting extensive tissue distribution in the underlying screening parameterization. The PoD-scaled unbound plasma C<sub>ss</sub> provides a conservative proxy for concentrations available to cross the BBB. A full mechanistic brain exposure estimate would require a brain partition coefficient or a dedicated chlorpyrifos PBPK model; this is a priority extension enabled by the same ToxMCP workflow.

### 4.2 Sensitivity analysis: hepatic volume as a determinant of brain exposure

Next, we used the PBPK server’s sensitivity workflow to explore how an anatomical parameter influences brain exposure. Liver volume is a proxy for hepatic capacity in PBPK models, capturing both anatomical scaling and, indirectly, the space for metabolism-limited clearance processes. We executed three simulations that varied Organism|Liver|Volume while keeping all other parameters fixed: baseline (1.5), severe impairment (0.75), and hypertrophy (3.0). For each run we extracted the brain intracellular AUC.

This experiment demonstrates a key advantage of tool-based PBPK: parameter perturbations are explicit, logged, and reproducible. The same protocol can be extended to permeability parameters, enzyme activities, or transporter expression when the model supports them.

### 4.3 Virtual population micro-study: age variation and brain Cmax

To illustrate population-style workflows, we simulated three virtual individuals by varying age (20, 40, 60 years) while using the same acetaminophen model file. For each subject, the PBPK server executed the simulation and extracted the brain intracellular Cmax.

In these runs, brain Cmax was invariant across age (160.61 µmol/L for all three subjects). This result is informative in two ways. First, it demonstrates that the pipeline can be used to test expectations about variability. Second, it highlights a common modeling pitfall: demographic parameters may not propagate to physiological scaling unless the PBPK model explicitly couples them to organ volumes, flows, and clearance. In an applied setting, this invariance would trigger a model audit: which parameters are age-dependent, and are they active in the current configuration?

### 4.4 Hazard-to-kinetics: chlorpyrifos point of departure and brain-relevant exposure proxies

A central promise of federated toxicology tooling is rapid hazard-to-kinetics integration: retrieving a health-based point of departure (PoD) and linking it to internal dose metrics. We demonstrate this with chlorpyrifos, an organophosphate insecticide with well-established cholinesterase inhibition as a sensitive endpoint.

Using the EPA CompTox MCP server, chlorpyrifos was resolved to DTXSID4020458 (CASRN 2921-88-2). The server returned high-throughput toxicokinetic (HTTK) predictions for human kinetics, including a steady-state plasma concentration (Css) per unit oral dose, half-life, volume of distribution, and fraction unbound in plasma. As a conservative PoD proxy, we used the ATSDR chronic-duration oral minimal risk level (MRL) of 0.001 mg/kg/day, which is derived from a rat NOAEL for acetylcholinesterase inhibition with standard uncertainty factors.^30,31,32^

Because a chlorpyrifos-specific brain PBPK model file was not available in this demonstration environment, we used a PBK-style steady-state scaling as an initial proxy: Css scales approximately linearly with dose under first-order kinetics. We therefore scaled the CompTox HTTK Css estimate (per 1 mg/kg/day) to the PoD dose, and computed the corresponding unbound plasma Css as a lower bound on brain-available concentration.

### 4.5 Extensibility templates: consumer product chemicals and mechanistic anchoring

Beyond mechanistic PBPK outputs, toxicology workflows often begin with pragmatic exposure questions: Where is a chemical used, what routes dominate, and which populations are plausibly exposed? The CompTox ecosystem aggregates chemical use and product-presence data (e.g., CPDat) that can seed such analyses. ToxMCP supports a template workflow for consumer product chemicals:

1. resolve identity (DTXSID, CASRN),
2. retrieve product-use categories and any available concentration ranges,
3. retrieve HTTK parameters (Css/unit dose) for screening-level internal dose estimates,
4. optionally run PBPK simulations for high-priority compounds,
5. map hazard endpoints and mechanistic pathways (AOP), and
6. generate an auditable report bundle.

This template is particularly useful for rapid ‘what-if’ experiments during manuscript development. A small number of chemicals can be explored deeply (PBPK + AOP + QSAR), while a larger chemical set can be triaged with CompTox/ADMETlab to justify prioritization. The resulting narrative can explicitly separate high-confidence mechanistic analyses from screening-level predictions, improving scientific clarity.

### 4.6 Multi-chemical benchmark suite and evidence bundles

To demonstrate scale beyond a small number of hand-curated case studies, we assembled a benchmark set of 12 chemicals spanning pharmaceuticals, pesticides, and consumer product ingredients. For each chemical, the multi-MCP pipeline resolves canonical identifiers, retrieves hazard and toxicokinetic screening data (CompTox/HTTK), maps mechanistic context (AOP), assigns QSAR hazard classes (O-QT), and retrieves product-use signals (CPDat where available). Figure 6 summarizes dossier completeness across domains, and Figure 7 illustrates a simple hazard-to-kinetics prioritization view derived from the cached benchmark artifacts.^33^

**Figure 6.**
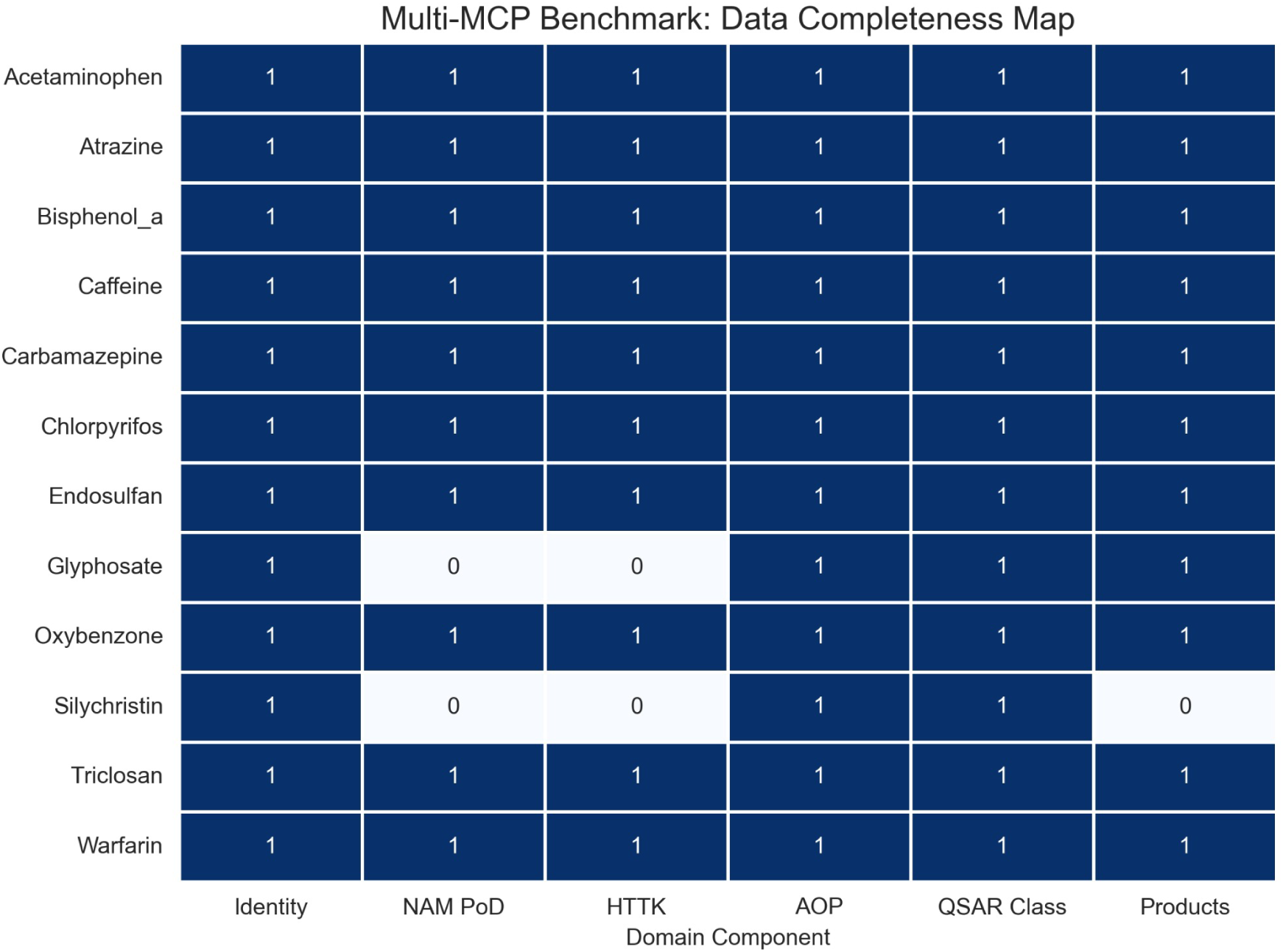
Multi-MCP benchmark coverage across a 12-chemical v1 set. The completeness map records whether each dossier includes the core evidence components: canonical identity, NAM PoD, HTTK kinetics, AOP mapping, QSAR class, and consumer product presence. This provides a transparent benchmark for coverage and failure-mode analysis.

**Figure 7.**
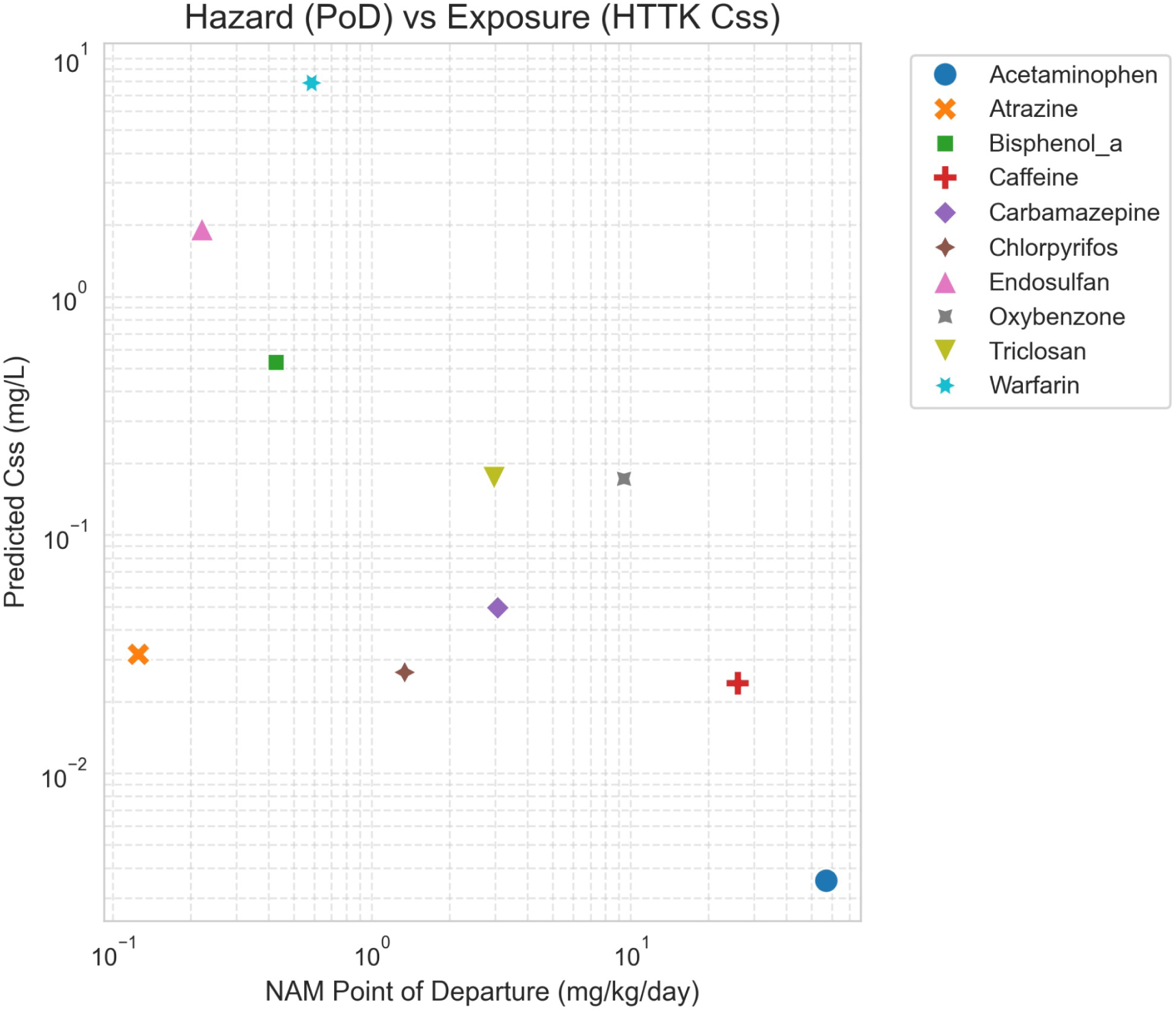
Hazard-to-kinetics prioritization from cached benchmark artifacts. For each chemical, we pair a screening-level NAM point of departure (PoD) with HTTK-predicted steady-state concentration (Css) per unit dose, enabling rapid ranking and identification of high-exposure / low-PoD regimes for deeper modeling.

We package these outputs as an evidence bundle (Supplementary Archive S1) containing per-chemical dossiers (raw MCP JSON responses + derived artifacts), a benchmark summary table, and scripts to reproduce figures from cached results. This design allows readers to inspect the full provenance of each value reported in the manuscript and to replay workflows under controlled conditions.

## 5. Evaluation Plan

### 5.1 What should be evaluated in agentic toxicology?

Evaluating an agentic toxicology framework requires more than measuring whether an LLM produces fluent text. The unit of evaluation is a workflow: a sequence of tool calls that transforms a scientific question into computational evidence and an interpretable report. We therefore propose evaluation axes that reflect toxicology practice: correctness of retrieved identifiers and values, reproducibility of computations, appropriateness of model usage (e.g., respecting applicability domain), and efficiency gains relative to manual workflows. Here, ‘reproducibility’ refers to computational reproducibility (re-running the same workflow yields the same outputs), distinct from experimental replicability or biological reproducibility.

ToxMCP provides natural instrumentation points for such evaluation because tool calls are explicit. Every workflow yields a machine-readable trace that can be compared against reference traces, replayed under controlled conditions, and scored by automated tests.

### 5.4 Proposed benchmark task suite

We outline a benchmark suite spanning common computational toxicology tasks, each designed to require multi-step reasoning and cross-tool integration. The suite covers: resolving ambiguous chemical mentions and regulatory context; retrieving appropriate points of departure (PoDs) for specified endpoints and exposure routes; screening exposure and toxicokinetics by combining HTTK predictions with user-defined scenarios; executing PBPK models (including controlled parameter edits) and extracting tissue AUC/Cmax/Tmax; running sensitivity and uncertainty analyses (parameter sweeps or small virtual populations) and summarizing directional effects; anchoring hazard endpoints mechanistically by mapping to AOP elements and assembling an evidence matrix; and executing QSAR Toolbox read-across workflows that export a dossier/report artifact.

To support both reproducibility and breadth, we recommend a two-tier benchmarking strategy. In this preprint, we provide a v1 evidence bundle covering a 12-chemical multi-MCP benchmark (Supplementary Archive S1), with cached artifacts enabling deterministic replay. Building on this, future releases can include: (i) a small ‘gold’ set of fully reproducible workflows (5–10) that exercise the full server stack end-to-end, and (ii) a larger screening set (50–200 chemicals) emphasizing breadth, where only selected steps (identity resolution, HTTK, ADMET) are executed automatically and cached for replay.

### 5.3 Metrics and scoring

We propose four classes of quantitative metrics:

1. Workflow validity: percentage of tasks completed without schema errors, missing identifiers, or policy violations.
2. Reproducibility: replay success rate (same trace → same outputs), hash stability of artifacts, and variance of numerical results.
3. Scientific compliance: rate at which outputs include required metadata (applicability domain flags, data source citations, model versions).
4. Efficiency: wall-clock time, number of tool calls, and human intervention points compared to a manual baseline.

For PBPK runs, reproducibility should be assessed at multiple levels: model file hash, simulation configuration hash, and key PK metric regression within tolerance. For knowledge retrieval (CompTox/AOP), reproducibility should consider API versioning and caching, and should capture retrieval timestamps and dataset identifiers.

Using the benchmark harness shipped in the evidence bundle (Supplementary Archive S1), we replay each PBPK model twice from a fresh session and assert metric regression within a 1% tolerance. In the current bundle, all PBPK metrics were identical within numerical precision (Figure 8; Supplementary Figures S1–S2).

**Figure 8.**
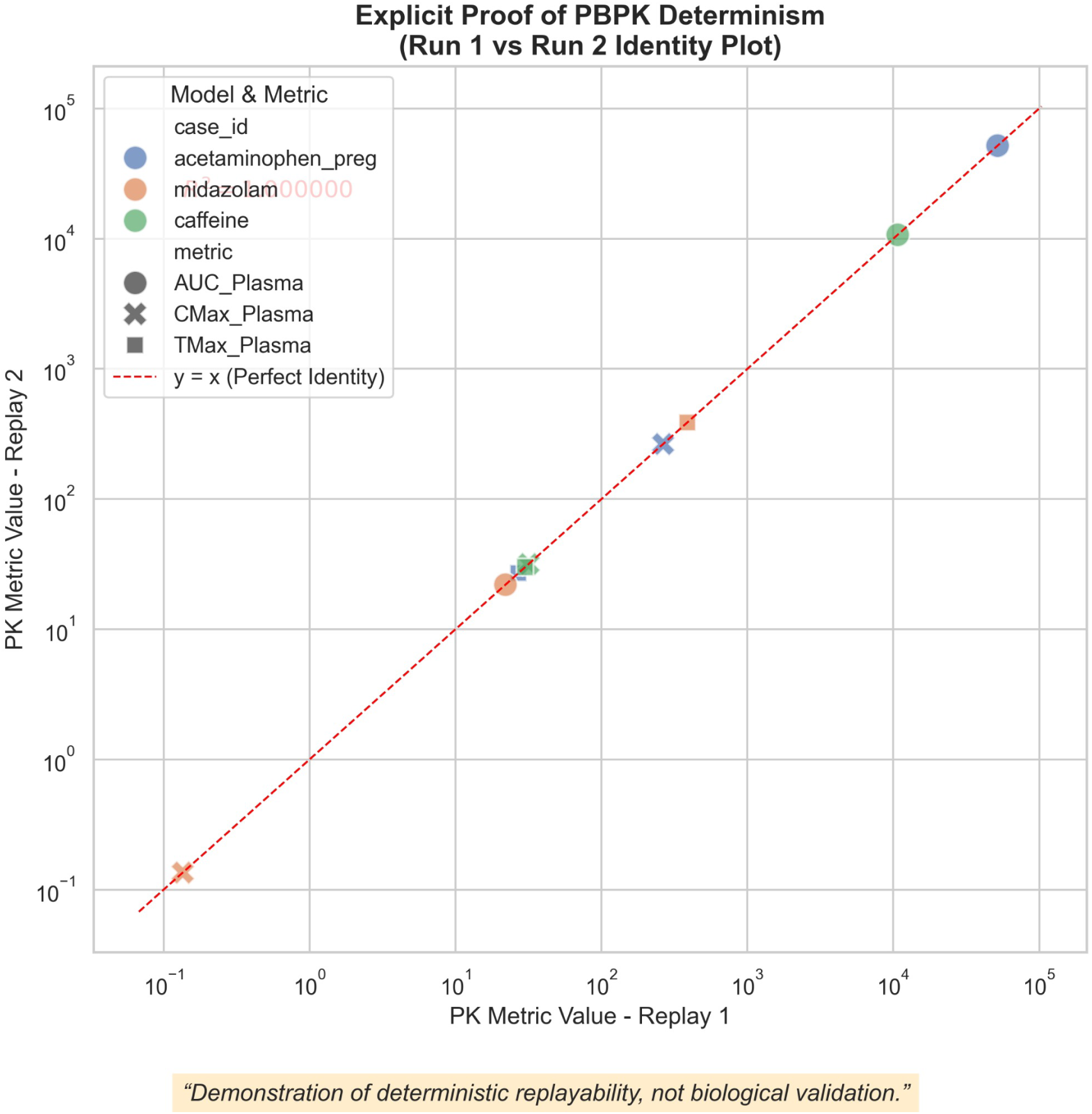
Deterministic replayability (‘engine truth’) for PBPK outputs. Identity plot comparing key PK metrics (AUC, Cmax, Tmax) across two independent replays of three PBPK models. Points lying on the y=x line indicate deterministic parity given identical model files and parameterization.

Qualitative evaluation remains important for high-impact publication. We recommend a blinded expert review in which toxicologists score report clarity, appropriateness of model choice, and whether key limitations are communicated. Because ToxMCP exposes traces, such review can focus on scientific judgment rather than reconstructing what was done.

### 5.4 Baselines

Baselines should reflect realistic alternatives. We propose three:

Baseline 1: manual workflow using web dashboards/GUI tools (CompTox Dashboard, QSAR Toolbox GUI, PBPK GUI execution).

Baseline 2: scripted workflow without LLM orchestration (hand-written Python/R scripts calling the same APIs).

Baseline 3: ‘chat-only’ LLM workflow (no tool calls), which should be expected to fail on reproducibility and audit criteria.

These baselines isolate the contribution of the MCP integration layer. In particular, Baseline 2 helps quantify whether ToxMCP’s main advantage is not just automation but the packaging of policy, provenance, and interoperability.

### 5.5 Anticipated failure modes and mitigations

Agentic toxicology introduces distinct failure modes: incorrect identity mapping, silent misuse of a model outside its domain, incomplete provenance, and overconfident narrative synthesis. ToxMCP mitigates these through: (i) canonical identifiers as mandatory outputs, (ii) policy hooks that can fail closed, (iii) structured outputs with required metadata, and (iv) report artifacts that can be independently reviewed.

We explicitly treat ‘guardrail-triggered refusal’ as a successful outcome in evaluation when it prevents invalid analysis. For example, a QSAR tool call that is blocked due to applicability domain mismatch is preferable to a plausible but non-defensible prediction.

### 5.6 Benchmark packaging and evidence bundle contract

For the benchmark to be machine-checkable, each task instance must specify (i) canonical inputs, (ii) required tool calls and constraints, (iii) expected outputs with tolerances, and (iv) minimum evidence artifacts. This turns ‘agentic performance’ from a qualitative impression into a testable contract that can be replayed and scored automatically.

We propose the following minimal evidence contract per task: (1) a machine-readable task specification (chemical identifier, endpoint, scenario parameters); (2) a complete tool-call trace (JSONL) including all inputs, outputs, and policy decisions; (3) cached raw tool outputs and derived artifacts (reports, tables, figures); (4) model/data version pins and cryptographic hashes for any executed kernels (e.g., PBPK model file, simulation configuration); and (5) an expected-metrics manifest (expected_metrics.json) plus a validator script that asserts schema compliance and numerical regression within tolerance.

Scoring should explicitly separate correctness, provenance, and safety/compliance. Correctness compares reported values against expected metrics (or reference traces) within defined tolerances; provenance requires that every reported numeric value links to a traceable artifact identifier in the bundle; and safety/compliance requires that disallowed actions are prevented and logged. Importantly, guardrail-triggered refusal is scored as correct when it prevents an invalid analysis (e.g., applicability-domain mismatch), provided that the refusal is explicit and auditable.

## 6. Discussion

### 6.1 From ‘AI-generated text’ to ‘AI-orchestrated evidence’

The main conceptual shift of ToxMCP is to treat the LLM as an orchestrator rather than as an authority. In this framing, the scientific kernel remains the source of truth, and the LLM’s role is to translate intent into explicit computations and to assemble interpretable narratives around verified outputs. This is aligned with how toxicology decisions are made: not by accepting a single model output, but by integrating multiple evidence streams and documenting assumptions.

This also clarifies why MCP matters. Many agent frameworks can call tools, but MCP provides a protocol-level standard that is independent of any one LLM vendor or orchestration library. Because the interface boundary is stable, labs can swap the ‘reasoning model’ layer (e.g., proprietary cloud models vs. on-premise open-weight models) without rewriting integrations to PBPK/QSAR/AOP kernels. This reduces vendor lock-in and supports data-sovereignty constraints by keeping sensitive inputs and licensed kernels within institutional boundaries.^19,20^

### 6.2 Alignment with regulatory expectations

ToxMCP was designed with regulatory-adjacent expectations in mind. OECD guidance for PBK models stresses the need for transparent documentation of model structure, parameters, and validation context. OECD’s (Q)SAR Assessment Framework similarly emphasizes applicability domain, transparency, and uncertainty communication. By embedding policy checks and provenance at the tool boundary, ToxMCP makes it easier to operationalize these expectations in daily computational practice.^21,22,26^

However, ToxMCP is not a replacement for regulatory review. It is a workflow accelerator and an evidence organizer. Its outputs must still be interpreted by experts, and any downstream decision-making should follow established risk assessment frameworks. The goal is to reduce clerical friction and to increase traceability, not to automate scientific judgement.

### 6.3 Limitations

Several limitations should be noted. First, data availability remains a bottleneck: a hazard-to-kinetics analysis is only as good as the accessible PoDs, exposure scenarios, and kinetic models. For many chemicals, HTTK parameters may be available but validated PBPK models are not. ToxMCP makes this gap explicit by separating screening-level proxies from mechanistic simulations.

Second, external services evolve. CompTox APIs, ADMETlab endpoints, and knowledge bases may change over time. ToxMCP mitigates this with explicit version capture and conformance testing, but long-term stability will require maintenance and community governance.

Third, agentic workflows can amplify errors if not carefully bounded. We intentionally include critical-action confirmation and RBAC hooks, but safe deployment also depends on client behavior and institutional policies.

Finally, the current demonstrations focus on a small number of workflows. A larger empirical evaluation across many chemicals and endpoints is necessary to quantify generality and to identify where agentic approaches offer the largest benefit.

### 6.4 Roadmap

We view adoption and validation as part of the technical contribution, not an afterthought. The near-term roadmap therefore combines scientific extensions with community-facing actions: (1) **Documentation and onboarding**: a versioned Quickstart, tutorial notebooks, and ‘gold trace’ workflows that reproduce each case study end-to-end (including audit-bundle export); (2) **Community validation**: a benchmark suite and conformance tests for each server (schema validation, regression tests for deterministic kernels, and failure-mode tests for guardrails); (3) **Governance artefacts**: a maintained glossary that expands Box 1, plus lightweight contribution and deprecation policies for tool schemas and audit bundles; (4) **Reference deployment**: a hosted pilot instance (e.g., within VHP4Safety infrastructure) for community evaluation, training, and inter-lab reproducibility exercises, subject to institutional approvals and licensing constraints for upstream tooling; and (5) **Scientific extensions**: parameterisable PBPK templates for chemicals without dedicated models, deeper AOP integration (including assay-to-key-event mapping), FAIR-aligned evidence-bundle formats, and automated generation of manuscript-ready figures and supplementary tables from stored artefacts.

In the longer term, we envision ToxMCP as a community-maintained suite where new domain servers (omics, exposure modelling, mixture toxicology) can be added without breaking interoperability. Because MCP is a protocol rather than a platform, participation does not require vendor lock-in: contributors implement typed tools, adopt shared schemas for evidence and audit bundles, and ship regression/conformance tests that make behaviour reviewable. This community focus is also the mechanism by which governance terminology, policy hooks, and evaluation criteria can converge over time.

For community adoption and external validation, we will publish versioned documentation and tutorial workflows, release conformance ‘gold’ traces and regression tests, maintain transparent contribution and governance processes, and (where feasible) operate an optional reference deployment (e.g., via VHP4Safety) for interoperability testing and evaluation.

## 7. Conclusion

ToxMCP provides an infrastructure blueprint for accountable agentic workflows in computational toxicology. By exposing established toxicology kernels and databases as typed MCP tools with guardrails and provenance, ToxMCP enables LLMs to accelerate multi-step analyses without collapsing scientific rigor into free-text speculation. The resulting workflows are reproducible, auditable, and extensible properties that are essential for both high-impact academic work and regulatory-adjacent decision support. We anticipate that protocol-level tool standardization will become a foundational ingredient for trustworthy AI in toxicology, and we release ToxMCP as a step toward that future.

## 8. Data and Code Availability

All ToxMCP servers are intended to be released as open-source code with containerized deployment recipes, alongside example MCP client workflows (notebooks/scripts) and conformance tests. Where upstream tools are licensed or non-redistributable (e.g., QSAR Toolbox), the repository provides integration stubs, schema definitions, and documentation rather than redistributing binaries.

Licensing note: Supplementary Archive S1 redistributes only derived result artifacts (e.g., exported JSON summaries / report artifacts) generated using third-party tools and does not include the OECD QSAR Toolbox installer/executables or donated databases/modules covered by the QSAR Toolbox End User Licence Agreement (EULA); reproducing QSAR workflows therefore requires users to separately download and use QSAR Toolbox under its EULA. PBPK simulations are executed with the Open Systems Pharmacology Suite (PK-Sim®/MoBi®; GPLv2), and the archived PKML model files are provided as data fixtures enabling deterministic replay; this bundle does not ship OSP binaries and assumes users install OSP separately.

For this submission, we provide a submission-ready evidence bundle as Supplementary Archive S1 (ToxMCP_Benchmark_Submission.zip) containing: (i) per-chemical dossiers with raw MCP responses (JSON) and derived artifacts (reports, traces), (ii) benchmark summary tables (CSV/Markdown) used for figures and tables, (iii) canonical PBPK model files (PKML) or recorded SHA-256 hashes where redistribution is restricted, and (iv) a ‘gold’ expected-metrics manifest (expected_metrics.json). A top-level MANIFEST.md inventories all files with SHA-256 checksums to support integrity checks.

To verify integrity and reproduce results, (1) check file hashes against MANIFEST.md, (2) run scripts/validate_bundle.py to confirm schema compliance and metric parity, and (3) optionally re-run PBPK simulations via the replay harness (run_benchmark.py) to test deterministic parity of AUC/Cmax/Tmax given identical model inputs. This explicitly separates ‘engine truth’ from biological validation.

For each case study, we recommend publishing the corresponding tool-call trace and audit-bundle identifiers as supplementary material; this allows third parties to replay workflows on their own infrastructure while verifying that the same evidence objects are produced.

## Appendix A. Tool catalogue (abridged)

### A.1 PBPK MCP

- list_models(): list available PBPK model files/templates.
- load_simulation(pkml_path): load a PK-Sim/MoBi model into a session and return a simulationId.
- list_parameters(simulationId, path_prefix=None, limit=…): enumerate parameter paths for discovery and validation.
- get_parameter_value(simulationId, parameterPath): read a parameter value.
- set_parameter_value(simulationId, parameterPath, value) [critical]: mutate a parameter (logged and permissioned).
- run_simulation(simulationId, …) [critical]: execute a simulation asynchronously and return a jobId/resultId.
- get_job_status(jobId): poll status for long-running jobs.
- calculate_pk_parameters(resultId, outputPath): compute AUC/Cmax/Tmax (and other metrics as configured).
- run_sensitivity_analysis(simulationId, parameterPath, values): run multiple perturbation scenarios and return a comparative summary.
- run_population_simulation(simulationId, subjects) [critical]: simulate multiple subjects and return a claim-check reference.
- export_results(resultRef, format): export results (CSV/JSON) for supplementary material.

### A.2 EPA CompTox MCP

- search_chemical(query): resolve names/identifiers to canonical DTXSID and related keys.
- get_chemical_details(dtxsid): retrieve curated identity and link-outs.
- get_chemical_properties(dtxsid): retrieve physicochemical properties and descriptors where available.
- get_exposure_httk(dtxsid, species=’human’): retrieve HTTK-based toxicokinetic predictions (Css/unit dose, t1/2, fu,p, Vd).
- search_hazard(dtxsid, endpoint/route/duration filters): retrieve hazard values and candidate PoDs.
- get_hazard_details(hazard_id): retrieve detailed provenance and supporting fields for a selected hazard value.

### A.3 ADMETlab MCP

- wash_smiles(smiles): canonicalize and validate SMILES.
- wash_batch_smiles(smiles_list): batch canonicalization for high-throughput workflows.
- render_svg(smiles): generate structure rendering for reports.
- predict_admet(smiles): retrieve ADMET prediction panel for a single molecule.
- predict_admet_batch(smiles_list): batch prediction for screening sets.

### A.4 AOP MCP

- search_aops(query): find relevant AOPs by keyword or identifier.
- get_aop(aop_id): retrieve full AOP object (title, MIE, AO, KEs, KERs).
- list_key_events(aop_id) / list_kers(aop_id): retrieve structured key events and relationships.
- map_chemical_to_aops(identifier): map chemicals to AOP-relevant entities where cross-links exist.
- get_evidence_matrix(aop_id): retrieve structured evidence supporting KERs and AOP plausibility.
- create_draft_aop(…): assemble a draft narrative skeleton from structured content.

### A.5 O-QT MCP

- profile_endpoint(smiles, endpoint): generate QSAR Toolbox profiling for a target endpoint.
- find_analogues(smiles, filters): identify candidate analogues for read-across.
- run_readacross(target, analogues, endpoint): execute read-across and capture evidence tables.
- generate_pdf_report(workflow_id) [critical]: export a dossier-style PDF report artifact.
- export_matrix(workflow_id): export evidence matrices for supplementary material.

## Appendix B. Example interaction traces (excerpt)

The following excerpts illustrate how natural-language tasks can be compiled into tool calls. For full traces, we recommend exporting MCP message logs as supplementary files.

Example B1: BBB analysis (acetaminophen proxy). A user requests a BBB comparison; the client loads the model, sets Organism|Age, runs the simulation (critical), computes PK metrics, and returns a table plus interpretation. Example B2: liver volume sensitivity. A user requests three parameter scenarios; the client runs three simulations and returns a percent-change table.

## Appendix C. Author contributions, acknowledgements, and disclosures (placeholders)

### C.1 Author contributions

I.D. conceived the ToxMCP framework, implemented the MCP server suite, designed the case studies and evaluation plan, and wrote the manuscript.

### C.2 Acknowledgements

We thank the participants of VHP4 Safety project for feedback on toxicology workflows, PBK modelling, and protocol design. We also acknowledge the communities and maintainers of the software and data resources that underpin the ToxMCP server suite, including the Open Systems Pharmacology Suite (PK-Sim and MoBi), the U.S. EPA CompTox/DSSTox ecosystem and HTTK resources, ADMETlab 3.0, the OECD QSAR Toolbox, and the AOP-Wiki/AOP-DB communities. This work was supported by the Dutch Research Council (NWO) through the VHP4Safety project.

### C.3 Competing interests

The authors declare no competing interests.

### C.4 Ethics statement

This work uses publicly available datasets and in silico simulations; no new experiments involving humans or animals were performed.

## Appendix D. Supplementary Figures

**Supplementary Figure S1.**
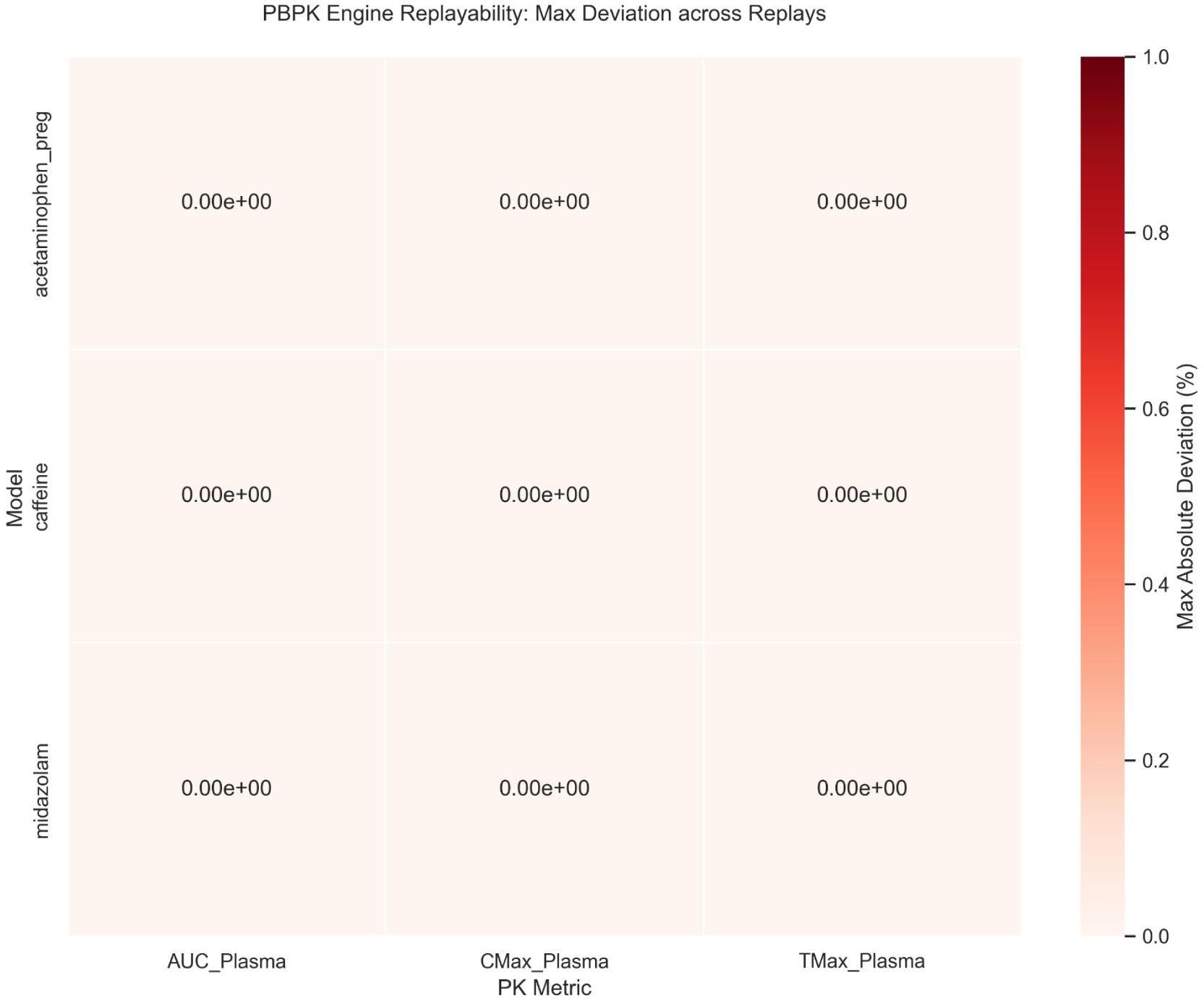
Deterministic replay heatmap for PBPK metrics across two runs and three models (1% tolerance).

**Supplementary Figure S2.**
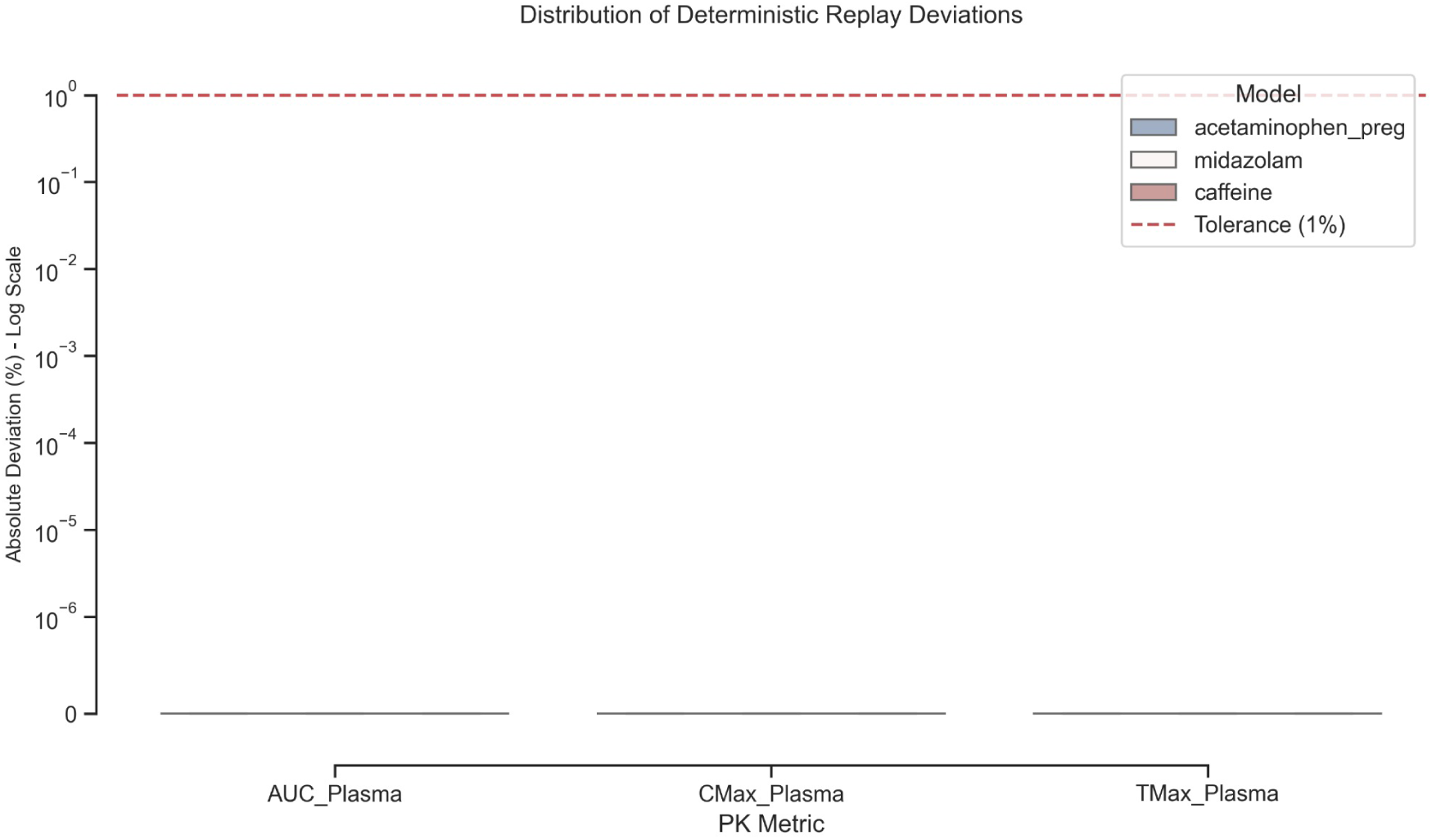
Distribution of deterministic replay deviations (absolute %), shown on a log scale. All deviations are at or near machine precision in the current bundle.

**Supplementary Figure S3.**
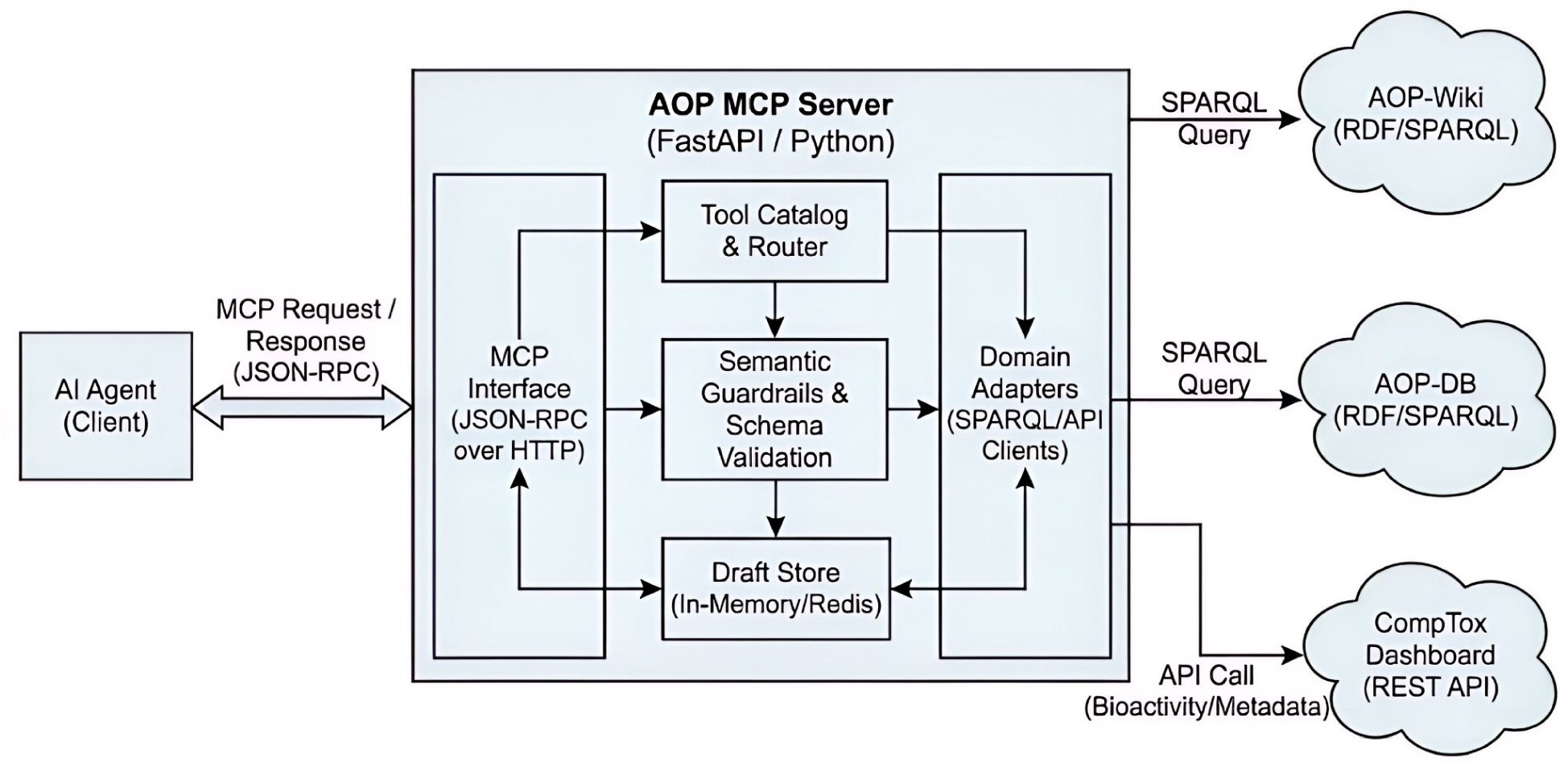
AOP MCP server architecture and data flow. The server routes MCP requests to RDF/SPARQL sources (e.g., AOP-Wiki) through schema validation and semantic guardrails.

**Supplementary Figure S4.**
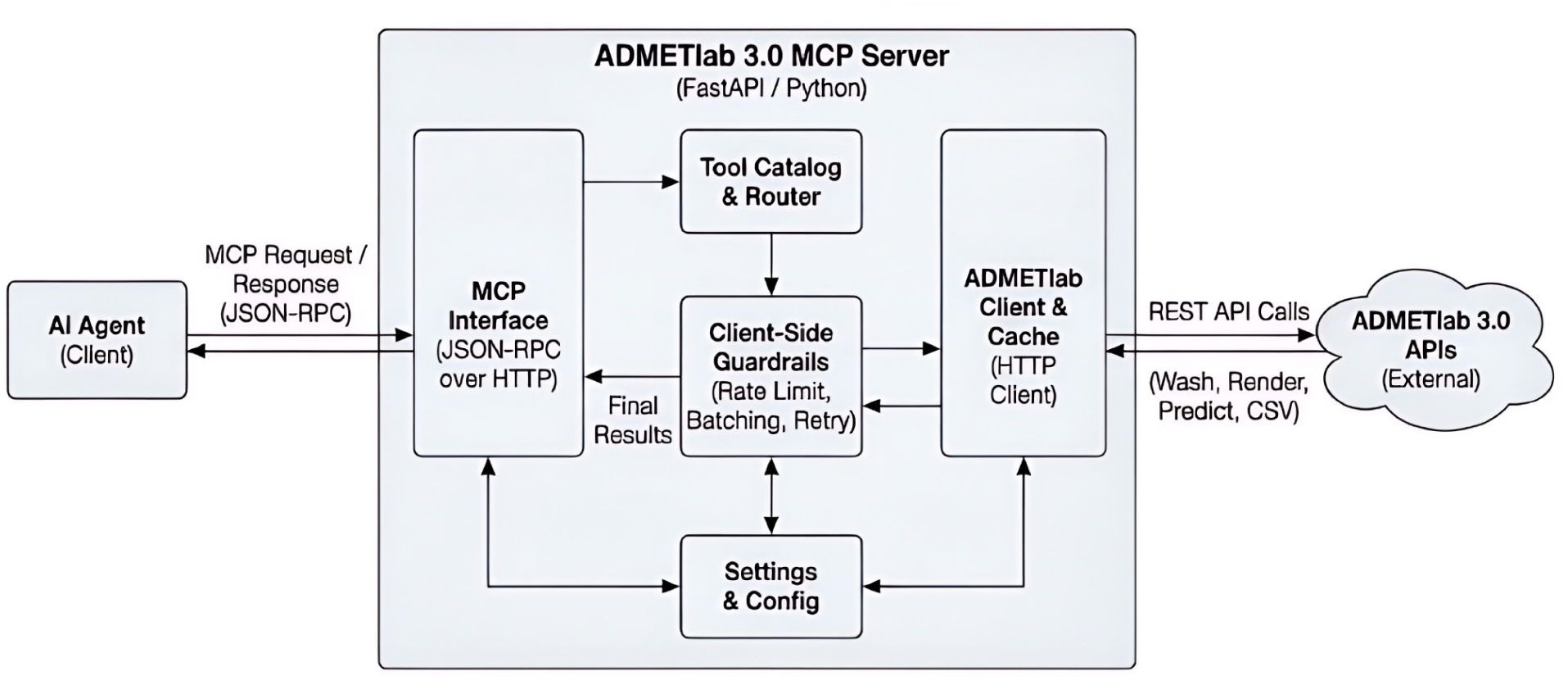
ADMETlab 3.0 MCP server architecture and data flow. A local MCP façade adds batching, retry, caching, and rate limits around external ADMETlab APIs.

**Supplementary Figure S5.**
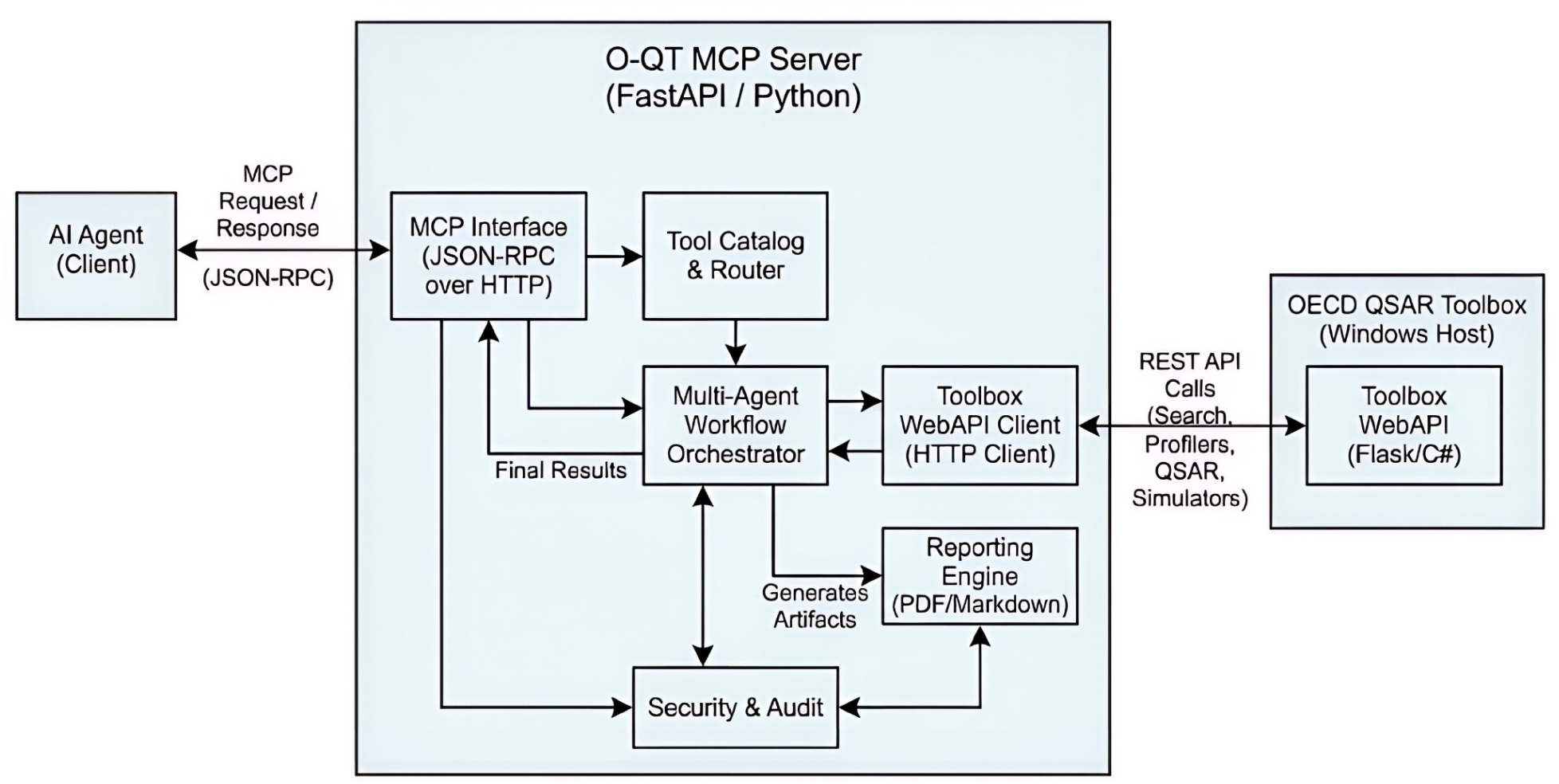
O-QT MCP server architecture and data flow. The MCP layer mediates multi-step QSAR Toolbox workflows, artifact generation, and security/audit requirements.

