## Supplementary material for "ToxMCP: Guardrailed, Auditable Agentic Workflows for Computational Toxicology via the Model Context Protocol": CHANGELOG.md

# Changelog — Evidence Bundle Hardening

## 2026-02-04 (QA hardening)
- Added `README.md` at archive root (usage, integrity verification, reproduction modes).
- Added `MANIFEST.sha256` (SHA-256 for every file in the archive).
- Added `BUNDLE_INFO.json` (packaging metadata + link to source zip hash).
- Fixed minor formatting issues in `publication_artifacts/README.md`.
- Added derived `comptox_identity.json` per chemical (extracted from `comptox_details.json`).
- Added `scripts/validate_bundle.py` (structure + JSON parse + PBPK hash checks + dossier completeness).
- Added a one-paragraph licensing/redistribution note (QSAR Toolbox outputs; OSP PBPK fixtures).
