## Supplementary material for "ToxMCP: Guardrailed, Auditable Agentic Workflows for Computational Toxicology via the Model Context Protocol": EVIDENCE_BUNDLE_QA_REPORT.md

# Evidence Bundle QA Report — MCP_Benchmark_Supplementary

Source archive: `MCP_Benchmark_Supplementary.zip`  
SHA-256 (source archive): `e50a14c737e49b6d984482c02aa06ca32db78465803e70523d40a36c788874b1`

QA date (UTC): 2026-02-04T09:33:06Z

## Summary
✅ Structural integrity: PASS  
✅ JSON artifact validity: PASS (all JSON parse)  
✅ PBPK model identity: PASS (SHA-256 matches `expected_metrics.json`)  
✅ Per-chemical dossier completeness: PASS (12/12 chemicals)  
⚠️ Re-run requirements: PARTIAL (scripts present, but environment dependencies are not pinned in this archive)  
⚠️ Licensing/citation metadata: PARTIAL (no explicit LICENSE/CITATION file shipped)

## What is in the bundle (high-level)
Top level:
- `run_benchmark.py` + `expected_metrics.json` : PBPK deterministic replay harness + golden metrics
- `models/` : canonical PBPK model files (hashed)
- `publication_artifacts/` : frozen outputs used in manuscript tables/figures
- `full_MCP_Suite/` : reference scripts (some require multiple MCP servers)

Publication artifacts:
- `benchmark_summary.csv` / `.md` : aggregated 12-chemical benchmark results
- `figures/` : PNG figures used for manuscript
- `{chemical}/` : per-chemical JSON dossiers (CompTox details/HTTK/AED/CPDat, AOP mapping, O-QT Cramer class)

## Validation details
### PBPK model hash parity
All model hashes match the expected values in `expected_metrics.json`:
- Acetaminophen_Pregnancy.pkml
- Midazolam_Canonical.pkml
- Caffeine_Canonical.pkml

### Dossier completeness
Each chemical folder contains:
- `comptox_details.json`
- `comptox_aed.json`
- `comptox_httk.json`
- `comptox_cpdat.json`
- `oqt_cramer.json`
- `aop_map.json` or `aop_map_empty.json`

## Recommendations before journal submission
1. Add an archive-wide cryptographic manifest (`MANIFEST.sha256`) and a root `README.md`.
2. Add a minimal validator script so reviewers can check structure/hashes quickly.
3. Add `CITATION.cff` and a clear `LICENSE` (and clarify any third-party data/model redistribution terms).
4. Consider depositing the archive to Zenodo to obtain a DOI, then cite that DOI in the manuscript (preprint servers may restrict large supplementary uploads).

---
Generated by automated checks in this environment.
