## Supplementary material for "ToxMCP: Guardrailed, Auditable Agentic Workflows for Computational Toxicology via the Model Context Protocol": benchmark_summary.md

| Chemical      | CAS       | NAM_PoD_AED     | HTTK_Css_Median   | PBPK_Brain_Blood_Ratio   | AOP_Mapping                                                 | QSAR_Cramer_Class   |   Product_Count |
|:--------------|:----------|:----------------|:------------------|:-------------------------|:------------------------------------------------------------|:--------------------|----------------:|
| Acetaminophen | 103-90-2  | 56.99 mg/kg/day | N/A               | 0.982                    | 1 AOPs (e.g. CYP2E1 activation and formation of protein...) | N/A                 |               0 |
| Chlorpyrifos  | 2921-88-2 | 1.343 mg/kg/day | 0.0265 mg/L       | 0.982                    | 3 AOPs (e.g. Inhibitor binding to topoisomerase II lead...) | High (Class III)    |               0 |
| Triclosan     | 3380-34-5 | 2.974 mg/kg/day | N/A               | N/A                      | 2 AOPs (e.g. Uncoupling of oxidative phosphorylation le...) | N/A                 |             266 |
