## Supplementary material for "ToxMCP: Guardrailed, Auditable Agentic Workflows for Computational Toxicology via the Model Context Protocol": debug_extras.py

```
import json
import requests
import time

SERVERS = {
    "AOP": "http://localhost:803/mcp",
    "O-QT": "http://localhost:804/mcp"
}

def mcp_call(server_name, tool_name, arguments):
    url = SERVERS[server_name]
    payload = {
        "jsonrpc": "2.0",
        "method": "tools/call",
        "params": {
            "name": tool_name,
            "arguments": arguments
        },
        "id": int(time.time() * 1000)
    }
    response = requests.post(url, json=payload)
    response.raise_for_status()
    data = response.json()
    if "error" in data and data["error"]:
        print(f"Error calling {server_name}/{tool_name}: {data['error']}")
        return None
    
    result = data.get("result", {})
    if "structuredContent" in result:
        return result["structuredContent"]
    if "content" in result:
        for item in result["content"]:
            if item["type"] == "text":
                try:
                    return json.loads(item["text"])
                except:
                    return item["text"]
    return result

def debug_aop():
    print("=== Debugging AOP MCP ===")
    # Chlorpyrifos InChIKey: SBPBAQFWLVIOKP-UHFFFAOYSA-N
    # Acetaminophen InChIKey: RZVAJINKPMORJF-UHFFFAOYSA-N
    
    # Try different identifiers for Chlorpyrifos
    ids = [
        {"inchikey": "SBPBAQFWLVIOKP-UHFFFAOYSA-N"},
        {"name": "Chlorpyrifos"},
        {"cas": "2921-88-2"}
    ]
    
    for args in ids:
        print(f"Testing AOP mapping with: {args}...")
        res = mcp_call("AOP", "map_chemical_to_aops", args)
        print(json.dumps(res, indent=2))

def explore_oqt():
    print("\n=== Exploring O-QT MCP Capabilities ===")
    capabilities = {}
    
    # 1. Calculators
    print("1. Listing Calculators...")
    calcs = mcp_call("O-QT", "list_calculators", {})
    if calcs and "calculators" in calcs:
        capabilities["calculators"] = calcs["calculators"]
        print(f"Found {len(calcs['calculators'])} calculators.")
    
    # 2. Profilers
    print("\n2. Listing Profilers...")
    profs = mcp_call("O-QT", "list_profilers", {})
    if profs and "profilers" in profs:
        capabilities["profilers"] = profs["profilers"]
        print(f"Found {len(profs['profilers'])} profilers.")
        
    # 3. QSAR Models
    print("\n3. Listing QSAR Models...")
    # 'list_qsar_models' requires position, 'list_all_qsar_models' does not
    models = mcp_call("O-QT", "list_all_qsar_models", {})
    if models and "models" in models:
        capabilities["qsar_models"] = models["models"]
        print(f"Found {len(models['models'])} QSAR models.")
        
    # Save artifacts
    with open("full_MCP_Suite/bench_runs/oqt_capabilities.json", "w") as f:
        json.dump(capabilities, f, indent=2)
    print("\nSaved capabilities to 'full_MCP_Suite/bench_runs/oqt_capabilities.json'")

if __name__ == "__main__":
    # debug_aop() # AOP is working fine, just empty for InChIKey sometimes
    explore_oqt()
```
