## Supplementary material for "ToxMCP: Guardrailed, Auditable Agentic Workflows for Computational Toxicology via the Model Context Protocol": fetch_aop_details.py

```
import json
import time
import requests
import os

SERVER_URL = "http://localhost:803/mcp"

def mcp_call(tool_name, arguments):
    payload = {
        "jsonrpc": "2.0",
        "method": "tools/call",
        "params": {
            "name": tool_name,
            "arguments": arguments
        },
        "id": int(time.time() * 1000)
    }
    try:
        response = requests.post(SERVER_URL, json=payload, timeout=30)
        response.raise_for_status()
        data = response.json()
        
        result = data.get("result", {})
        if "structuredContent" in result:
            return result["structuredContent"]
        if "content" in result:
            for item in result["content"]:
                if item["type"] == "text":
                    try:
                        return json.loads(item["text"])
                    except:
                        return item["text"]
        return result
    except Exception as e:
        print(f"Error: {e}")
        return None

def fetch_aop_details():
    aops = {
        "chlorpyrifos": "559",
        "triclosan": "263",
        "acetaminophen": "260"
    }
    
    base_dir = "full_MCP_Suite/bench_runs/aops"
    os.makedirs(base_dir, exist_ok=True)
    
    for chem, aop_id in aops.items():
        print(f"Fetching details for {chem} -> AOP:{aop_id}...")
        
        # 1. Metadata
        meta = mcp_call("get_aop", {"aop_id": aop_id})
        if meta:
            with open(os.path.join(base_dir, f"{chem}_{aop_id}_meta.json"), "w") as f:
                json.dump(meta, f, indent=2)
                
        # 2. Key Events
        kes = mcp_call("list_key_events", {"aop_id": aop_id})
        if kes:
            with open(os.path.join(base_dir, f"{chem}_{aop_id}_kes.json"), "w") as f:
                json.dump(kes, f, indent=2)
                
        # 3. KERs
        kers = mcp_call("list_kers", {"aop_id": aop_id})
        if kers:
            with open(os.path.join(base_dir, f"{chem}_{aop_id}_kers.json"), "w") as f:
                json.dump(kers, f, indent=2)
                
        print(f"  Saved artifacts for {chem}.")

if __name__ == "__main__":
    fetch_aop_details()
```
