## Supplementary material for "ToxMCP: Guardrailed, Auditable Agentic Workflows for Computational Toxicology via the Model Context Protocol": generate_benchmark_summary.py

```
import json
import os
import pandas as pd

BENCH_DIR = "publication_artifacts"

def load_json(path):
    if not os.path.exists(path):
        return None
    try:
        with open(path, 'r') as f:
            return json.load(f)
    except:
        return None

def extract_summary():
    rows = []
    if not os.path.exists(BENCH_DIR):
        print(f"Directory {BENCH_DIR} not found.")
        return

    chemicals = sorted([d for d in os.listdir(BENCH_DIR) if os.path.isdir(os.path.join(BENCH_DIR, d))])
    
    for chem in chemicals:
        dir_path = os.path.join(BENCH_DIR, chem)
        row = {"Chemical": chem.capitalize()}
        
        # 1. Identity
        ident = load_json(os.path.join(dir_path, "comptox_details.json"))
        # Fallback to identity file if details missing (though CS scripts save details)
        if not ident:
             ident = load_json(os.path.join(dir_path, "comptox_identity.json"))
             
        if ident:
             # Details format
             if "casrn" in ident: 
                 row["CAS"] = ident["casrn"]
             # Search format
             elif "data" in ident and ident["data"]:
                 row["CAS"] = ident["data"][0].get("casrn", "N/A")
             else:
                 row["CAS"] = "N/A"
        else:
            row["CAS"] = "N/A"
            
        # 2. NAM PoD (AED)
        aed = load_json(os.path.join(dir_path, "comptox_aed.json"))
        pod_val = "N/A"
        if aed and "data" in aed and aed["data"]:
            vals = aed["data"][0].get("aedVal", [])
            if vals:
                pod_val = f"{vals[0]} mg/kg/day"
        row["NAM_PoD_AED"] = pod_val

        # 3. HTTK Css
        httk = load_json(os.path.join(dir_path, "comptox_httk.json"))
        css_val = "N/A"
        if httk and "data" in httk and httk["data"]:
            for rec in httk["data"]:
                if rec.get("parameter") == "Css" and rec.get("percentile") == "50%":
                    css_val = f"{rec.get('predicted')} {rec.get('units')}"
                    break
        row["HTTK_Css_Median"] = css_val
            
        # 4. PBPK Ratio
        pbpk = load_json(os.path.join(dir_path, "pbpk_metrics.json"))
        ratio = "N/A"
        if pbpk and "metrics" in pbpk:
            brain_auc = None
            plasma_auc = None
            for m in pbpk["metrics"]:
                if m["parameter"] == "Organism|Brain|Intracellular|Paracetamol|Concentration":
                    brain_auc = m["auc"]
                if m["parameter"] == "Organism|PeripheralVenousBlood|Paracetamol|Plasma (Peripheral Venous Blood)":
                    plasma_auc = m["auc"]
            if brain_auc and plasma_auc:
                ratio = round(brain_auc / plasma_auc, 3)
        row["PBPK_Brain_Blood_Ratio"] = ratio

        # 5. AOP Mapping
        aop = load_json(os.path.join(dir_path, "aop_map.json"))
        aop_text = "None"
        if aop and "results" in aop and aop["results"]:
            count = len(aop["results"])
            first_aop = aop["results"][0]["aop"].get("title", "").strip()
            if len(first_aop) > 45: first_aop = first_aop[:42] + "..."
            aop_text = f"{count} AOPs (e.g. {first_aop})"
        row["AOP_Mapping"] = aop_text

        # 6. O-QT Profiling (Cramer)
        oqt = load_json(os.path.join(dir_path, "oqt_report.json"))
        cramer = "N/A"
        
        # Try full report first
        if oqt and "log_json" in oqt:
            for prof in oqt["log_json"].get("profiler_results", []):
                if prof.get("profiler_guid") == "aa645923-a592-46fe-9069-737a2e4a7ac6":
                    res_list = prof.get("result", [])
                    if len(res_list) > 1:
                        cramer = res_list[1]
                        break
        
        # Try lightweight cramer artifact if not found
        if cramer == "N/A":
            oqt_cramer = load_json(os.path.join(dir_path, "oqt_cramer.json"))
            if oqt_cramer and "result" in oqt_cramer:
                 # result usually [category, class, etc] or similar list
                 res_list = oqt_cramer["result"]
                 if isinstance(res_list, list) and len(res_list) > 1:
                     cramer = res_list[1]
                 elif isinstance(res_list, list) and len(res_list) > 0:
                     cramer = res_list[0]

        row["QSAR_Cramer_Class"] = cramer
        
        # 7. Products
        cpdat = load_json(os.path.join(dir_path, "comptox_cpdat.json"))
        row["Product_Count"] = len(cpdat["data"]) if cpdat and "data" in cpdat and cpdat["data"] else 0
            
        rows.append(row)
        
    df = pd.DataFrame(rows)
    csv_path = os.path.join(BENCH_DIR, "benchmark_summary.csv")
    df.to_csv(csv_path, index=False)
    
    md_path = os.path.join(BENCH_DIR, "benchmark_summary.md")
    df.to_markdown(md_path, index=False)
    
    print("\n=== Nature Quality Publication Summary Table ===\n")
    print(df.to_markdown(index=False))
    print(f"\nFinal artifacts updated in {BENCH_DIR}")

if __name__ == "__main__":
    extract_summary()
```
