## Supplementary material for "ToxMCP: Guardrailed, Auditable Agentic Workflows for Computational Toxicology via the Model Context Protocol": generate_publication_figures.py

```
import json
import os
import pandas as pd
import matplotlib.pyplot as plt
import seaborn as sns
import numpy as np
import re

ARTIFACTS_DIR = "publication_artifacts"
FIGURES_DIR = os.path.join(ARTIFACTS_DIR, "figures")
os.makedirs(FIGURES_DIR, exist_ok=True)

def load_json(path):
    if not os.path.exists(path): return None
    with open(path, 'r') as f: return json.load(f)

def generate_figure_1_sensitivity():
    print("Generating Figure 1: PBPK Sensitivity Analysis...")
    sens_path = os.path.join(ARTIFACTS_DIR, "acetaminophen", "pbpk_sensitivity.json")
    data = load_json(sens_path)
    if not data or "scenarios" not in data.get("report", {}):
        print("Sensitivity data not found or invalid.")
        return

    scenarios = data["report"]["scenarios"]
    plot_data = []
    
    # Target metric: Brain concentration AUC
    target_param = "Organism|Brain|Intracellular|Paracetamol|Concentration"
    
    for sc in scenarios:
        x_val = sc.get("percent_change", 0) * 100 # Convert to %
        auc = 0
        for m in sc.get("metrics", []):
            if m["parameter"] == target_param:
                auc = m["auc"]
                break
        plot_data.append({"Change_Liver_Volume_Percent": x_val, "Brain_AUC": auc})
    
    df = pd.DataFrame(plot_data).sort_values("Change_Liver_Volume_Percent")
    
    plt.figure(figsize=(6, 5))
    sns.set_style("whitegrid")
    sns.lineplot(data=df, x="Change_Liver_Volume_Percent", y="Brain_AUC", marker="o", linewidth=2.5)
    plt.title("PBPK Sensitivity: Liver Volume vs Brain Exposure\n(Acetaminophen Pregnancy Model)", fontsize=12)
    plt.xlabel("Change in Liver Volume (%)", fontsize=10)
    plt.ylabel("Brain Paracetamol AUC (µmol*min/L)", fontsize=10)
    plt.tight_layout()
    plt.savefig(os.path.join(FIGURES_DIR, "Figure_1_PBPK_Sensitivity.png"), dpi=300)
    plt.close()

def generate_figure_2_completeness():
    print("Generating Figure 2: Tox Benchmark Completeness Heatmap...")
    csv_path = os.path.join(ARTIFACTS_DIR, "benchmark_summary.csv")
    if not os.path.exists(csv_path):
        print("Benchmark summary CSV not found.")
        return
        
    df = pd.read_csv(csv_path)
    
    # Create completeness matrix
    completeness = pd.DataFrame()
    completeness["Chemical"] = df["Chemical"]
    
    # Logic: 1 = Data present, 0 = Missing/None
    completeness["Identity"] = df["CAS"].apply(lambda x: 1 if x != "N/A" else 0)
    completeness["NAM PoD"] = df["NAM_PoD_AED"].apply(lambda x: 1 if "mg/kg" in str(x) else 0)
    completeness["HTTK"] = df["HTTK_Css_Median"].apply(lambda x: 1 if "mg/L" in str(x) else 0)
    completeness["AOP"] = df["AOP_Mapping"].apply(lambda x: 0 if "None" in str(x) else 1)
    completeness["QSAR Class"] = df["QSAR_Cramer_Class"].apply(lambda x: 1 if x != "N/A" else 0)
    completeness["Products"] = df["Product_Count"].apply(lambda x: 1 if x > 0 else 0)
    
    completeness = completeness.set_index("Chemical")
    
    plt.figure(figsize=(8, 6))
    sns.heatmap(completeness, cmap="Blues", cbar=False, annot=True, linewidths=1, linecolor='white')
    plt.title("Multi-MCP Benchmark: Data Completeness Map", fontsize=14)
    plt.xlabel("Domain Component")
    plt.ylabel("")
    plt.tight_layout()
    plt.savefig(os.path.join(FIGURES_DIR, "Figure_2_Data_Completeness.png"), dpi=300)
    plt.close()

def extract_numeric(val_str):
    if not isinstance(val_str, str): return None
    match = re.search(r"([\d\.]+)", val_str)
    if match:
        return float(match.group(1))
    return None

def generate_figure_3_scatter():
    print("Generating Figure 3: PoD vs HTTK Scatter...")
    csv_path = os.path.join(ARTIFACTS_DIR, "benchmark_summary.csv")
    df = pd.read_csv(csv_path)
    
    df["PoD_Value"] = df["NAM_PoD_AED"].apply(extract_numeric)
    df["HTTK_Css"] = df["HTTK_Css_Median"].apply(extract_numeric)
    
    # Filter valid pairs
    plot_df = df.dropna(subset=["PoD_Value", "HTTK_Css"])
    
    if plot_df.empty:
        print("No valid data points for scatter plot.")
        return

    plt.figure(figsize=(7, 6))
    sns.scatterplot(data=plot_df, x="PoD_Value", y="HTTK_Css", s=100, hue="Chemical", style="Chemical")
    
    plt.xscale("log")
    plt.yscale("log")
    plt.title("Hazard (PoD) vs Exposure (HTTK Css)", fontsize=14)
    plt.xlabel("NAM Point of Departure (mg/kg/day)", fontsize=11)
    plt.ylabel("Predicted Css (mg/L)", fontsize=11)
    plt.legend(bbox_to_anchor=(1.05, 1), loc='upper left')
    plt.grid(True, which="both", ls="--", alpha=0.5)
    plt.tight_layout()
    plt.savefig(os.path.join(FIGURES_DIR, "Figure_3_PoD_vs_HTTK.png"), dpi=300)
    plt.close()

if __name__ == "__main__":
    generate_figure_1_sensitivity()
    generate_figure_2_completeness()
    generate_figure_3_scatter()
    print(f"Figures saved to {FIGURES_DIR}")
```
