## Supplementary material for "ToxMCP: Guardrailed, Auditable Agentic Workflows for Computational Toxicology via the Model Context Protocol": inspect_artifacts.py

```
import json
import os
import glob

def inspect_json_files(root_dir):
    print(f"Inspecting JSON artifacts in {root_dir}...\n")
    
    for chem_dir in os.listdir(root_dir):
        chem_path = os.path.join(root_dir, chem_dir)
        if not os.path.isdir(chem_path):
            continue
            
        print(f"--- Chemical: {chem_dir} ---")
        json_files = sorted(glob.glob(os.path.join(chem_path, "*.json")))
        
        if not json_files:
            print("  No JSON files found.")
            continue
            
        for json_file in json_files:
            filename = os.path.basename(json_file)
            try:
                with open(json_file, 'r') as f:
                    content = f.read()
                    if not content.strip():
                        print(f"  ❌ {filename}: Empty file")
                        continue
                        
                    try:
                        data = json.loads(content)
                    except json.JSONDecodeError:
                        # Sometimes it might be just a string if my previous save logic was loose
                        print(f"  ❌ {filename}: Invalid JSON content: {content[:50]}...")
                        continue

                # content check
                status = "✅ OK"
                details = ""
                
                # Check for explicit errors
                if isinstance(data, dict) and "error" in data and data["error"]:
                    status = "❌ ERROR"
                    details = f"Error field present: {data['error']}"
                
                # Check for empty results in common structures
                elif isinstance(data, dict) and "data" in data and isinstance(data["data"], list):
                    if not data["data"]:
                        status = "⚠️ WARNING"
                        details = "Empty 'data' list"
                    else:
                        details = f"Records: {len(data['data'])}"
                
                # Check for specific files
                elif filename == "comptox_identity.json":
                    if isinstance(data, list) and not data:
                         status = "⚠️ WARNING"
                         details = "Empty search result"
                    elif isinstance(data, dict) and "data" in data:
                         # handled above
                         pass
                
                elif filename == "oqt_report.json":
                    if isinstance(data, dict):
                        # Check for PDF or log presence
                        keys = list(data.keys())
                        details = f"Keys: {keys}"
                
                elif filename == "pbpk_metrics.json":
                    if isinstance(data, dict) and "metrics" in data:
                        details = f"Metrics count: {len(data['metrics'])}"
                    else:
                        status = "⚠️ WARNING"
                        details = "No 'metrics' key found"

                print(f"  {status} {filename} ({details})")
                
            except Exception as e:
                print(f"  ❌ {filename}: Read failed ({str(e)})")

if __name__ == "__main__":
    inspect_json_files("full_MCP_Suite/bench_runs/chemicals")
```
