## Supplementary material for "ToxMCP: Guardrailed, Auditable Agentic Workflows for Computational Toxicology via the Model Context Protocol": mcp_utils.py

```
import json
import time
import requests
import os

SERVERS = {
    "PBPK": "http://localhost:8000/mcp",
    "CompTox": "http://localhost:8001/mcp",
    "AOP": "http://localhost:803/mcp",
    "O-QT": "http://localhost:804/mcp"
}

def mcp_call(server_name, tool_name, arguments):
    url = SERVERS.get(server_name)
    if not url:
        print(f"Unknown server: {server_name}")
        return None
        
    payload = {
        "jsonrpc": "2.0",
        "method": "tools/call",
        "params": {
            "name": tool_name,
            "arguments": arguments
        },
        "id": int(time.time() * 1000)
    }
    
    try:
        response = requests.post(url, json=payload, timeout=300) # 5 min timeout for long sims
        response.raise_for_status()
        data = response.json()
        
        if "error" in data and data["error"]:
            print(f"Error calling {server_name}/{tool_name}: {data['error']}")
            return None
        
        result = data.get("result", {})
        if "structuredContent" in result:
            return result["structuredContent"]
        if "content" in result:
            for item in result["content"]:
                if item["type"] == "text":
                    try:
                        return json.loads(item["text"])
                    except:
                        return item["text"]
        return result
        
    except Exception as e:
        print(f"Exception calling {server_name}/{tool_name}: {e}")
        return None

def save_artifact(chemical, filename, data):
    base_dir = f"publication_artifacts/{chemical}"
    os.makedirs(base_dir, exist_ok=True)
    path = os.path.join(base_dir, filename)
    with open(path, "w") as f:
        json.dump(data, f, indent=2)
    print(f"Saved: {path}")

def get_best_aop_mapping(chem_key, chem_name, cas, inchikey):
    """
    Tries to map to AOPs using InChIKey -> CAS -> Name priority.
    Returns the best result found.
    """
    print(f"5. AOP: Mapping to pathways for {chem_name}...")
    
    aop_map = None
    method = None
    
    # 1. Try InChIKey
    if inchikey:
        print(f"  Querying by InChIKey ({inchikey})...")
        res = mcp_call("AOP", "map_chemical_to_aops", {"inchikey": inchikey})
        if res and "results" in res and res["results"]:
            aop_map = res
            method = "InChIKey"
            
    # 2. Try CAS
    if (not aop_map or not aop_map.get("results")) and cas:
        print(f"  Querying by CAS ({cas})...")
        res = mcp_call("AOP", "map_chemical_to_aops", {"cas": cas})
        if res and "results" in res and res["results"]:
            aop_map = res
            method = "CAS"

    # 3. Try Name
    if (not aop_map or not aop_map.get("results")) and chem_name:
        print(f"  Querying by Name ({chem_name})...")
        res = mcp_call("AOP", "map_chemical_to_aops", {"name": chem_name})
        if res and "results" in res and res["results"]:
            aop_map = res
            method = "Name"
            
    if aop_map and aop_map.get("results"):
        count = len(aop_map["results"])
        print(f"  Found {count} AOP associations using {method}.")
        # Inject metadata about how it was found
        aop_map["metadata"] = {"match_method": method, "query_value": locals().get(method.lower())}
        save_artifact(chem_key, "aop_map.json", aop_map)
        return aop_map
    else:
        print("  No AOP associations found.")
        save_artifact(chem_key, "aop_map_empty.json", {"results": [], "attempts": ["inchikey", "cas", "name"]})
        return None
```
