## Supplementary material for "ToxMCP: Guardrailed, Auditable Agentic Workflows for Computational Toxicology via the Model Context Protocol": plot_aop_pathway.py

```
import json
import os
import networkx as nx
import matplotlib.pyplot as plt
import seaborn as sns

# Setup
AOP_DIR = "full_MCP_Suite/bench_runs/aops"
OUTPUT_DIR = "full_MCP_Suite/figures"
os.makedirs(OUTPUT_DIR, exist_ok=True)

# Style
sns.set_theme(style="white")
plt.rcParams.update({'font.size': 10})

def load_json(path):
    if not os.path.exists(path):
        return None
    try:
        with open(path, 'r') as f:
            return json.load(f)
    except:
        return None

def plot_aop_559():
    print("Generating Figure 12: AOP 559 Pathway Architecture (Chlorpyrifos)...")
    
    kes_path = os.path.join(AOP_DIR, "chlorpyrifos_559_kes.json")
    kers_path = os.path.join(AOP_DIR, "chlorpyrifos_559_kers.json")
    
    kes_data = load_json(kes_path)
    kers_data = load_json(kers_path)
    
    if not kes_data or not kers_data:
        print("  Skipping: Data missing")
        return

    G = nx.DiGraph()
    
    # 1. Add Nodes
    # Map ID to Title
    titles = {}
    for ke in kes_data.get("results", []):
        titles[ke["id"]] = ke["title"]
        G.add_node(ke["id"], title=ke["title"])
        
    # 2. Add Edges
    for ker in kers_data.get("results", []):
        u = ker["upstream"]["id"]
        v = ker["downstream"]["id"]
        G.add_edge(u, v)
        
    # 3. Identify MIE and AO based on topology
    # MIE: In-degree 0
    # AO: Out-degree 0
    mies = [n for n in G.nodes if G.in_degree(n) == 0]
    aos = [n for n in G.nodes if G.out_degree(n) == 0]
    
    # 4. Visualization
    plt.figure(figsize=(12, 8))
    
    # Hierarchical layout attempt (multipartite)
    # Simple layers: MIE -> intermediates -> AO
    # Calculate distance from MIEs
    if mies:
        layers = {}
        for n in G.nodes:
            try:
                # shortest path length from any MIE
                dists = [len(nx.shortest_path(G, m, n)) for m in mies if nx.has_path(G, m, n)]
                if dists:
                    layers[n] = max(dists)
                else:
                    layers[n] = 0 # Disconnected or self?
            except:
                layers[n] = 0
        
        # Assign layers
        for n, l in layers.items():
            G.nodes[n]["layer"] = l
            
        pos = nx.multipartite_layout(G, subset_key="layer", align="horizontal")
    else:
        pos = nx.spring_layout(G, seed=42)

    # Node Colors
    color_map = []
    for n in G.nodes:
        if n in mies:
            color_map.append("#2ecc71") # Green (Start)
        elif n in aos:
            color_map.append("#e74c3c") # Red (End)
        else:
            color_map.append("#3498db") # Blue (Intermediate)
            
    # Draw
    nx.draw_networkx_edges(G, pos, edge_color="gray", arrows=True, arrowsize=20, width=1.5, connectionstyle="arc3,rad=0.1")
    nx.draw_networkx_nodes(G, pos, node_color=color_map, node_size=3000, alpha=0.9, edgecolors="black")
    
    # Labels (Wrap text)
    import textwrap
    labels = {n: textwrap.fill(titles.get(n, n), width=15) for n in G.nodes}
    nx.draw_networkx_labels(G, pos, labels, font_size=8, font_weight="bold")
    
    plt.title("AOP 559: Acetylcholinesterase Inhibition to Cardiac Arrhythmia\n(Chlorpyrifos Mechanism of Action)", fontsize=16, weight='bold')
    plt.axis("off")
    
    # Legend
    import matplotlib.patches as mpatches
    mie_patch = mpatches.Patch(color='#2ecc71', label='Molecular Initiating Event (MIE)')
    ke_patch = mpatches.Patch(color='#3498db', label='Key Event (KE)')
    ao_patch = mpatches.Patch(color='#e74c3c', label='Adverse Outcome (AO)')
    plt.legend(handles=[mie_patch, ke_patch, ao_patch], loc="lower right")
    
    out = os.path.join(OUTPUT_DIR, "Figure_12_AOP_559_Pathway.png")
    plt.savefig(out, dpi=300, bbox_inches="tight")
    print(f"  Saved: {out}")
    plt.close()

if __name__ == "__main__":
    plot_aop_559()
```
