## Supplementary material for "ToxMCP: Guardrailed, Auditable Agentic Workflows for Computational Toxicology via the Model Context Protocol": plot_extended.py

```
import pandas as pd
import matplotlib.pyplot as plt
import seaborn as sns
import os
import json

# Setup
BENCH_DIR = "full_MCP_Suite/bench_runs/chemicals"
OUTPUT_DIR = "full_MCP_Suite/figures"
os.makedirs(OUTPUT_DIR, exist_ok=True)

# Style
sns.set_theme(style="whitegrid")
plt.rcParams.update({'font.size': 12})

def load_json(path):
    if not os.path.exists(path):
        return None
    try:
        with open(path, 'r') as f:
            return json.load(f)
    except:
        return None

def plot_pbpk_sensitivity():
    print("Generating Figure 6: PBPK Sensitivity...")
    path = os.path.join(BENCH_DIR, "acetaminophen", "pbpk_sensitivity.json")
    data = load_json(path)
    
    if not data or "report" not in data:
        print("  Skipping: Data missing")
        return

    # Extract scenarios
    scenarios = []
    # Baseline (1.5L implied or explicit?)
    # The JSON structure usually has "scenarios" list
    for s in data["report"]["scenarios"]:
        # Find Brain AUC metric
        brain_auc = 0
        for m in s["metrics"]:
            if "Brain" in m["parameter"] and "Concentration" in m["parameter"]:
                brain_auc = m["auc"]
                break
        
        # Determine Volume
        # scenario_id might contain the delta info
        # But better to rely on input if available. 
        # In this specific output, we have 'absolute_value' in the scenario dict!
        vol = s.get("absolute_value")
        if vol is None:
            # Infer from ID or delta?
            # "Organism_Liver_Volume_200" -> +20% -> 1.5 * 1.2 = 1.8? 
            # Wait, the artifact I saw earlier had 3.0L as baseline? No, 1.5L.
            # Let's trust "absolute_value" if present.
            pass
            
        scenarios.append({
            "Liver Volume (L)": vol,
            "Brain AUC (µmol*min/L)": brain_auc,
            "Scenario": "Baseline" if s["scenario_id"] == "baseline" else "Perturbation"
        })
        
    df = pd.DataFrame(scenarios).sort_values("Liver Volume (L)")
    
    plt.figure(figsize=(8, 6))
    
    # Line + Points
    sns.lineplot(data=df, x="Liver Volume (L)", y="Brain AUC (µmol*min/L)", marker="o", markersize=10, linewidth=2, color="#2ecc71")
    
    # Annotate points
    for i, row in df.iterrows():
        plt.text(row["Liver Volume (L)"], row["Brain AUC (µmol*min/L)"] + 1000, 
                 f"{row['Brain AUC (µmol*min/L)']:.0f}", ha='center')

    plt.title("PBPK Mechanistic Validation: Liver Volume Sensitivity\n(Acetaminophen Pregnancy Model)", fontsize=14, weight='bold')
    plt.xlabel("Liver Volume (L)", fontsize=12)
    plt.ylabel("Predicted Brain Exposure (AUC)", fontsize=12)
    
    # Add interpretation arrow
    # High volume -> Low AUC
    # Low volume -> High AUC
    # plt.annotate('Hypertrophy = Lower Risk?', xy=(3.0, 30000), xytext=(2.0, 40000),
    #             arrowprops=dict(facecolor='black', shrink=0.05))

    sns.despine()
    plt.tight_layout()
    
    out = os.path.join(OUTPUT_DIR, "Figure_6_PBPK_Sensitivity_Curve.png")
    plt.savefig(out, dpi=300)
    print(f"  Saved: {out}")
    plt.close()

def plot_triclosan_products():
    print("Generating Figure 7: Triclosan Product Exposure...")
    path = os.path.join(BENCH_DIR, "triclosan", "comptox_cpdat.json")
    data = load_json(path)
    
    if not data or "data" not in data:
        print("  Skipping: Data missing")
        return
        
    # Count categories
    cats = []
    for rec in data["data"]:
        # "gencat" seems to be "General Category" e.g. Personal care
        # "prodfam" -> Product Family e.g. shaving
        c = rec.get("gencat")
        if c: cats.append(c)
        
    df = pd.DataFrame(cats, columns=["Category"])
    counts = df["Category"].value_counts().reset_index()
    counts.columns = ["Category", "Count"]
    
    # Top 10
    top_n = counts.head(8)
    
    plt.figure(figsize=(10, 6))
    sns.barplot(data=top_n, y="Category", x="Count", palette="rocket")
    
    plt.title("Triclosan: Top Consumer Product Exposure Sources\n(EPA CPDat Evidence)", fontsize=14, weight='bold')
    plt.xlabel("Number of Registered Products", fontsize=12)
    plt.ylabel("")
    
    # Add labels
    for i, v in enumerate(top_n["Count"]):
        plt.text(v + 2, i, str(v), va='center')
        
    sns.despine()
    plt.tight_layout()
    
    out = os.path.join(OUTPUT_DIR, "Figure_7_Triclosan_Product_Landscape.png")
    plt.savefig(out, dpi=300)
    print(f"  Saved: {out}")
    plt.close()

def plot_httk_uncertainty():
    print("Generating Figure 8: HTTK Uncertainty...")
    path = os.path.join(BENCH_DIR, "chlorpyrifos", "comptox_httk.json")
    data = load_json(path)
    
    if not data or "data" not in data:
        print("  Skipping: Data missing")
        return
        
    # Extract Css predictions
    recs = []
    for r in data["data"]:
        if r.get("parameter") == "Css" and r.get("predicted") is not None:
            recs.append({
                "Model": r.get("model", "Unknown"),
                "Species": r.get("species", "Unknown"),
                "Percentile": r.get("percentile", "Unknown"),
                "Css (mg/L)": float(r["predicted"])
            })
            
    df = pd.DataFrame(recs)
    
    # Filter for Human vs Rat
    df = df[df["Species"].isin(["Human", "Rat"])]
    
    plt.figure(figsize=(10, 6))
    
    # Strip plot + Box plot combo
    sns.boxplot(data=df, x="Species", y="Css (mg/L)", hue="Model", palette="pastel", showfliers=False)
    sns.stripplot(data=df, x="Species", y="Css (mg/L)", hue="Model", dodge=True, color=".3", linewidth=1)
    
    plt.title("HTTK Predictive Uncertainty: Chlorpyrifos Kinetics\n(Model & Species Variability)", fontsize=14, weight='bold')
    plt.ylabel("Predicted Steady-State Concentration (mg/L)", fontsize=12)
    
    # Handle duplicate legend
    handles, labels = plt.gca().get_legend_handles_labels()
    # Slicing to remove duplicates if stripplot adds them
    l = len(df["Model"].unique())
    plt.legend(handles[:l], labels[:l], title="HTTK Model", loc="upper right")
    
    sns.despine()
    plt.tight_layout()
    
    out = os.path.join(OUTPUT_DIR, "Figure_8_HTTK_Kinetic_Uncertainty.png")
    plt.savefig(out, dpi=300)
    print(f"  Saved: {out}")
    plt.close()

if __name__ == "__main__":
    plot_pbpk_sensitivity()
    plot_triclosan_products()
    plot_httk_uncertainty()
```
