## Supplementary material for "ToxMCP: Guardrailed, Auditable Agentic Workflows for Computational Toxicology via the Model Context Protocol": plot_extra_figures.py

```
import pandas as pd
import matplotlib.pyplot as plt
import seaborn as sns
import os
import json
import networkx as nx

# Setup
BENCH_DIR = "full_MCP_Suite/bench_runs/chemicals"
OUTPUT_DIR = "full_MCP_Suite/figures"
os.makedirs(OUTPUT_DIR, exist_ok=True)

# Style
sns.set_theme(style="whitegrid")
plt.rcParams.update({'font.size': 12})

def load_json(path):
    if not os.path.exists(path):
        return None
    try:
        with open(path, 'r') as f:
            return json.load(f)
    except:
        return None

def plot_nam_distribution():
    print("Generating Figure 9: NAM PoD Distribution (Chlorpyrifos)...")
    path = os.path.join(BENCH_DIR, "chlorpyrifos", "comptox_aed.json")
    data = load_json(path)
    
    if not data or "data" not in data:
        print("  Skipping: Data missing")
        return
        
    # Extract AED values
    # The JSON structure has a list of records, each record has list of vals
    vals = []
    for rec in data["data"]:
        # Each record corresponds to a chemical (Chlorpyrifos)
        # But 'aedVal' is a list of values
        if "aedVal" in rec:
            vals.extend(rec["aedVal"])
            
    df = pd.DataFrame(vals, columns=["AED (mg/kg/day)"])
    
    plt.figure(figsize=(10, 6))
    
    # Violin + Strip
    sns.violinplot(data=df, x="AED (mg/kg/day)", color="lightblue", inner="quartile")
    sns.stripplot(data=df, x="AED (mg/kg/day)", color=".2", alpha=0.5, jitter=True)
    
    plt.title("High-Throughput Risk Assessment: NAM-based Point of Departure\n(Distribution of Active Equivalent Doses)", fontsize=14, weight='bold')
    plt.xlabel("Predicted Oral Equivalent Dose (mg/kg/day)", fontsize=12)
    
    # Log scale x-axis usually better for dose
    plt.xscale('log')
    
    # Add median annotation
    median = df["AED (mg/kg/day)"].median()
    plt.axvline(median, color='red', linestyle='--', label=f'Median: {median:.2f}')
    plt.legend()
    
    sns.despine()
    plt.tight_layout()
    
    out = os.path.join(OUTPUT_DIR, "Figure_9_NAM_PoD_Distribution.png")
    plt.savefig(out, dpi=300)
    print(f"  Saved: {out}")
    plt.close()

def plot_metabolic_fate():
    print("Generating Figure 10: Acetaminophen Metabolic Fate...")
    path = os.path.join(BENCH_DIR, "acetaminophen", "pbpk_metrics.json")
    data = load_json(path)
    
    if not data or "metrics" not in data:
        print("  Skipping: Data missing")
        return
        
    # Extract AUCs for "Sum" parameters (Total excreted mass usually)
    # Or Urine AUC which proxies total clearance
    
    # Metrics keys:
    # "Organism|Kidney|Urine|Paracetamol"
    # "Organism|Kidney|Urine|Paracetamol glucuronide"
    # "Organism|Kidney|Urine|Paracetamol sulfate"
    # "Organism|Kidney|Urine|NAPQI"
    
    components = {}
    for m in data["metrics"]:
        p = m["parameter"]
        if "Organism|Kidney|Urine|" in p and "|Concentration" not in p:
            name = p.split("|")[-1]
            components[name] = m["auc"] # AUC of Amount (µmol) is total mass over time if integrated? 
            # Wait, AUC of amount in urine? Amount in urine accumulates. 
            # Cmax of Amount in Urine = Total Excreted Amount at end.
            # Let's use Cmax for Urine Amount (µmol)
            components[name] = m["cmax"]

    if not components:
        print("  Skipping: No urine metrics found")
        return

    # Pie chart
    labels = list(components.keys())
    sizes = list(components.values())
    
    # Sort for better visual
    sorted_pairs = sorted(zip(sizes, labels), reverse=True)
    sizes = [x[0] for x in sorted_pairs]
    labels = [x[1] for x in sorted_pairs]
    
    plt.figure(figsize=(8, 8))
    colors = sns.color_palette('pastel')[0:len(labels)]
    
    plt.pie(sizes, labels=labels, autopct='%1.1f%%', startangle=140, colors=colors, textprops={'fontsize': 12})
    plt.title("Simulated Metabolic Fate: Urinary Excretion Profile\n(Acetaminophen Pregnancy Model)", fontsize=14, weight='bold')
    
    out = os.path.join(OUTPUT_DIR, "Figure_10_Metabolic_Fate.png")
    plt.savefig(out, dpi=300)
    print(f"  Saved: {out}")
    plt.close()

def plot_aop_network():
    print("Generating Figure 11: Chemical-AOP Network...")
    
    # Load all maps
    aop_data = []
    for chem in ["chlorpyrifos", "acetaminophen", "triclosan"]:
        path = os.path.join(BENCH_DIR, chem, "aop_map.json")
        d = load_json(path)
        if d and "results" in d:
            for r in d["results"]:
                aop_id = r["aop"]["id"]
                aop_title = r["aop"]["title"]
                aop_data.append({"Chemical": chem.capitalize(), "AOP": f"{aop_id}: {aop_title[:30]}..."})
                
    df = pd.DataFrame(aop_data)
    
    if df.empty:
        print("  Skipping: No AOP data")
        return

    # Build Graph
    G = nx.Graph()
    
    chemicals = df["Chemical"].unique()
    aops = df["AOP"].unique()
    
    for chem in chemicals:
        G.add_node(chem, type="chemical", color="skyblue")
        
    for aop in aops:
        G.add_node(aop, type="aop", color="lightgreen")
        
    for idx, row in df.iterrows():
        G.add_edge(row["Chemical"], row["AOP"])
        
    # Layout
    pos = nx.spring_layout(G, k=0.5, iterations=50)
    
    plt.figure(figsize=(12, 8))
    
    # Draw Nodes
    chem_nodes = [n for n,d in G.nodes(data=True) if d["type"]=="chemical"]
    aop_nodes = [n for n,d in G.nodes(data=True) if d["type"]=="aop"]
    
    nx.draw_networkx_nodes(G, pos, nodelist=chem_nodes, node_color="skyblue", node_size=3000, alpha=0.9)
    nx.draw_networkx_nodes(G, pos, nodelist=aop_nodes, node_color="#90EE90", node_size=2000, alpha=0.8, node_shape="s")
    
    # Draw Edges
    nx.draw_networkx_edges(G, pos, width=2, alpha=0.5, edge_color="gray")
    
    # Labels
    nx.draw_networkx_labels(G, pos, font_size=10, font_weight="bold")
    
    plt.title("Mechanistic Connectivity: Chemical-AOP Association Network", fontsize=16, weight='bold')
    plt.axis("off")
    
    # Legend
    import matplotlib.patches as mpatches
    chem_patch = mpatches.Patch(color='skyblue', label='Chemical')
    aop_patch = mpatches.Patch(color='#90EE90', label='Adverse Outcome Pathway')
    plt.legend(handles=[chem_patch, aop_patch], loc="upper left")
    
    out = os.path.join(OUTPUT_DIR, "Figure_11_AOP_Network.png")
    plt.savefig(out, dpi=300)
    print(f"  Saved: {out}")
    plt.close()

if __name__ == "__main__":
    plot_nam_distribution()
    plot_metabolic_fate()
    plot_aop_network()
```
