## Supplementary material for "ToxMCP: Guardrailed, Auditable Agentic Workflows for Computational Toxicology via the Model Context Protocol": plot_full_suite.py

```
import pandas as pd
import matplotlib.pyplot as plt
import seaborn as sns
import os
import json
import re

# Setup
INPUT_CSV = "full_MCP_Suite/bench_runs/benchmark_summary.csv"
CAPABILITIES_JSON = "full_MCP_Suite/bench_runs/oqt_capabilities.json"
OUTPUT_DIR = "full_MCP_Suite/figures"
os.makedirs(OUTPUT_DIR, exist_ok=True)

# Set style
sns.set_theme(style="whitegrid")
plt.rcParams.update({'font.size': 12})

def parse_aop_count(val):
    if pd.isna(val) or val == "None": return 0
    match = re.match(r"(\d+)", str(val))
    return int(match.group(1)) if match else 0

def parse_httk(val):
    if pd.isna(val) or val == "N/A": return 0.0
    match = re.match(r"([0-9\.]+)", str(val))
    return float(match.group(1)) if match else 0.0

def generate_dossier_figure():
    print("Generating Dossier Summary Figure...")
    df = pd.read_csv(INPUT_CSV)
    
    # Preprocessing
    df["AOP_Count_Num"] = df["AOP_Mapping"].apply(parse_aop_count)
    df["HTTK_Css_Num"] = df["HTTK_Css_Median"].apply(parse_httk)
    df["PBPK_Ratio_Num"] = pd.to_numeric(df["PBPK_Brain_Blood_Ratio"], errors='coerce').fillna(0)
    df["Product_Count"] = pd.to_numeric(df["Product_Count"], errors='coerce').fillna(0)
    
    # Create composite figure
    fig, axes = plt.subplots(2, 2, figsize=(14, 10))
    fig.suptitle("Multi-MCP Toxicology Benchmark: Dossier Summary", fontsize=16, weight='bold', y=0.95)
    
    # 1. AOP Counts
    sns.barplot(data=df, x="Chemical", y="AOP_Count_Num", ax=axes[0,0], palette="viridis", hue="Chemical", legend=False)
    axes[0,0].set_title("Adverse Outcome Pathway (AOP) Associations")
    axes[0,0].set_ylabel("Count")
    axes[0,0].set_xlabel("")
    for i, v in enumerate(df["AOP_Count_Num"]):
        axes[0,0].text(i, v + 0.1, str(v), ha='center')

    # 2. Product Counts (Log scale if Triclosan dominates)
    sns.barplot(data=df, x="Chemical", y="Product_Count", ax=axes[0,1], palette="magma", hue="Chemical", legend=False)
    axes[0,1].set_title("Consumer Product Associations (CPDat)")
    axes[0,1].set_ylabel("Count")
    axes[0,1].set_xlabel("")
    # Add labels
    for i, v in enumerate(df["Product_Count"]):
        label = str(int(v)) if v > 0 else "0"
        axes[0,1].text(i, v + 5, label, ha='center')

    # 3. PBPK Brain/Blood Ratio
    # Filter for non-zero
    pbpk_df = df[df["PBPK_Ratio_Num"] > 0]
    if not pbpk_df.empty:
        sns.barplot(data=pbpk_df, x="Chemical", y="PBPK_Ratio_Num", ax=axes[1,0], palette="coolwarm", hue="Chemical", legend=False)
        axes[1,0].set_title("PBPK Brain/Blood AUC Ratio")
        axes[1,0].set_ylabel("Ratio")
        axes[1,0].axhline(0.5, color='red', linestyle='--', label='BBB Penetration Threshold (0.5)')
        axes[1,0].legend()
        axes[1,0].set_ylim(0, 1.2)
        for i, row in enumerate(pbpk_df.itertuples()):
            axes[1,0].text(i, row.PBPK_Ratio_Num + 0.02, f"{row.PBPK_Ratio_Num}", ha='center')
    else:
        axes[1,0].text(0.5, 0.5, "No PBPK Data", ha='center')

    # 4. HTTK Css
    httk_df = df[df["HTTK_Css_Num"] > 0]
    if not httk_df.empty:
        sns.barplot(data=httk_df, x="Chemical", y="HTTK_Css_Num", ax=axes[1,1], palette="rocket", hue="Chemical", legend=False)
        axes[1,1].set_title("HTTK Predicted Steady-State Concentration")
        axes[1,1].set_ylabel("Css Median (mg/L)")
        axes[1,1].set_xlabel("")
        for i, row in enumerate(httk_df.itertuples()):
            axes[1,1].text(i, row.HTTK_Css_Num + 0.001, f"{row.HTTK_Css_Num} mg/L", ha='center')
    else:
        axes[1,1].text(0.5, 0.5, "No HTTK Data", ha='center')

    plt.tight_layout(rect=[0, 0.03, 1, 0.95])
    
    out_file = os.path.join(OUTPUT_DIR, "Figure_4_MultiMCP_Dossier_Summary.png")
    plt.savefig(out_file, dpi=300)
    print(f"Saved: {out_file}")
    plt.close()

def generate_capabilities_figure():
    print("Generating Capabilities Figure...")
    if not os.path.exists(CAPABILITIES_JSON):
        print("Capabilities JSON not found, skipping.")
        return

    with open(CAPABILITIES_JSON, 'r') as f:
        data = json.load(f)
        
    counts = {
        "Calculators": len(data.get("calculators", [])),
        "Profilers": len(data.get("profilers", [])),
        # "QSAR Models": len(data.get("qsar_models", [])) # Might be empty based on previous run, let's check
    }
    
    # QSAR models list was empty in previous run, so let's stick to what we have or check
    if data.get("qsar_models"):
        counts["QSAR Models"] = len(data["qsar_models"])

    df_cap = pd.DataFrame(list(counts.items()), columns=["Tool Type", "Count"])
    
    plt.figure(figsize=(8, 6))
    ax = sns.barplot(data=df_cap, x="Tool Type", y="Count", palette="mako", hue="Tool Type", legend=False)
    
    plt.title("O-QT MCP Server: Computational Capabilities", fontsize=14, weight='bold')
    plt.ylabel("Number of Available Tools")
    plt.xlabel("")
    
    for i, v in enumerate(df_cap["Count"]):
        ax.text(i, v + 2, str(v), ha='center', fontsize=12, weight='bold')
        
    sns.despine()
    plt.tight_layout()
    
    out_file = os.path.join(OUTPUT_DIR, "Figure_5_OQT_Capabilities.png")
    plt.savefig(out_file, dpi=300)
    print(f"Saved: {out_file}")
    plt.close()

if __name__ == "__main__":
    generate_dossier_figure()
    generate_capabilities_figure()
```
