## Supplementary material for "ToxMCP: Guardrailed, Auditable Agentic Workflows for Computational Toxicology via the Model Context Protocol": run_benchmark_batch.py

```
from full_MCP_Suite.mcp_utils import mcp_call, save_artifact, get_best_aop_mapping
import time
import sys

CHEMICALS = [
    # Case Studies
    "Acetaminophen",
    "Chlorpyrifos",
    "Triclosan",
    # Additional Chemicals
    "Bisphenol A",
    "Oxybenzone",
    "Caffeine",
    "Warfarin",
    "Atrazine",
    "Glyphosate",
    "Carbamazepine",
    "Silychristin", # User requested Silichristin
    "Endosulfan"
]

def run_batch_benchmark():
    print(f"=== Starting 10-Chemical Automated Benchmark for Publication ===")
    print(f"Target Directory: publication_artifacts/")
    
    for chem_name in CHEMICALS:
        chem_key = chem_name.lower().replace(" ", "_")
        print(f"\n--- Processing: {chem_name} ---")
        
        try:
            # 1. CompTox: Identity
            print("  1. CompTox: Resolving Identity...")
            chem_search = mcp_call("CompTox", "search_chemical", {"query": chem_name, "search_type": "equals"})
            
            dtxsid = None
            cas = None
            inchikey = None
            
            if chem_search and "data" in chem_search:
                for hit in chem_search["data"]:
                    if hit.get("preferredName").lower() == chem_name.lower():
                        dtxsid = hit.get("dtxsid")
                        cas = hit.get("casrn")
                        break
            
            # Fallback if preferred name doesn't match exactly (use first hit)
            if not dtxsid and chem_search and "data" in chem_search and len(chem_search["data"]) > 0:
                 dtxsid = chem_search["data"][0].get("dtxsid")
                 cas = chem_search["data"][0].get("casrn")

            if dtxsid:
                details = mcp_call("CompTox", "get_chemical_details", {"identifier": dtxsid, "id_type": "dtxsid"})
                if details:
                    inchikey = details.get("inchikey")
                    save_artifact(chem_key, "comptox_details.json", details)
                else:
                    # Save identity search result if details fail
                    save_artifact(chem_key, "comptox_identity.json", chem_search)

                # 2. CompTox: NAM Point of Departure (AED)
                print("  2. CompTox: Retrieving NAM PoD...")
                aed_res = mcp_call("CompTox", "get_bioactivity_aed", {"dtxsid": dtxsid})
                save_artifact(chem_key, "comptox_aed.json", aed_res)

                # 3. CompTox: HTTK
                print("  3. CompTox: Retrieving HTTK Data...")
                httk = mcp_call("CompTox", "search_httk", {"dtxsid": dtxsid})
                save_artifact(chem_key, "comptox_httk.json", httk)

                # 4. CompTox: CPDat
                print("  4. CompTox: Checking Product Data...")
                cpdat = mcp_call("CompTox", "search_cpdat", {"dtxsid": dtxsid, "vocab_name": "puc"})
                save_artifact(chem_key, "comptox_cpdat.json", cpdat)

            else:
                print(f"  [Warn] DTXSID not found for {chem_name}")

            # 5. AOP Mapping
            get_best_aop_mapping(chem_key, chem_name, cas, inchikey)

            # 6. O-QT: Cramer Rules Profiling (Lightweight)
            print("  6. O-QT: Running Cramer Rules Profiling...")
            oqt_search = mcp_call("O-QT", "search_chemicals", {"query": chem_name})
            
            oqt_chem_id = None
            if oqt_search and "results" in oqt_search and oqt_search["results"]:
                # Prefer High quality matches
                for res in oqt_search["results"]:
                    if res.get("CasSmilesRelation") == "High":
                        oqt_chem_id = res.get("ChemId")
                        break
                # Fallback to first
                if not oqt_chem_id:
                     oqt_chem_id = oqt_search["results"][0].get("ChemId")
            
            if oqt_chem_id:
                cramer_guid = "aa645923-a592-46fe-9069-737a2e4a7ac6" # Cramer rules
                profiler_res = mcp_call("O-QT", "run_profiler", {
                    "profiler_guid": cramer_guid,
                    "chem_id": oqt_chem_id
                })
                save_artifact(chem_key, "oqt_cramer.json", profiler_res)
            else:
                print(f"  [Warn] O-QT ChemId not found for {chem_name}")

        except Exception as e:
            print(f"  [Error] Failed processing {chem_name}: {e}")

    print("\n=== Batch Benchmark Complete ===")

if __name__ == "__main__":
    run_batch_benchmark()
```
