## Supplementary material for "ToxMCP: Guardrailed, Auditable Agentic Workflows for Computational Toxicology via the Model Context Protocol": run_cs1_chlorpyrifos.py

```
from full_MCP_Suite.mcp_utils import mcp_call, save_artifact, get_best_aop_mapping
import time
import json

def run_case_study_1_nature():
    print("=== Case Study 1: Chlorpyrifos (Nature Publication Quality) ===")
    chem_name = "Chlorpyrifos"
    chem_key = "chlorpyrifos" 
    
    # 1. CompTox: Identity
    print("1. CompTox: Resolving Identity...")
    chem_search = mcp_call("CompTox", "search_chemical", {"query": chem_name, "search_type": "equals"})
    
    dtxsid = None
    cas = None
    inchikey = None
    
    if chem_search and "data" in chem_search:
        for hit in chem_search["data"]:
            if hit.get("preferredName").lower() == chem_name.lower():
                dtxsid = hit.get("dtxsid")
                cas = hit.get("casrn")
                break
    
    if not dtxsid:
        print("Failed to resolve DTXSID.")
        return
        
    details = mcp_call("CompTox", "get_chemical_details", {"identifier": dtxsid, "id_type": "dtxsid"})
    if details:
        inchikey = details.get("inchikey")
        save_artifact(chem_key, "comptox_details.json", details)

    # 2. CompTox: NAM Point of Departure (AED)
    print("2. CompTox: Retrieving NAM-based Point of Departure (AED)...")
    aed_res = mcp_call("CompTox", "get_bioactivity_aed", {"dtxsid": dtxsid})
    save_artifact(chem_key, "comptox_aed.json", aed_res)

    # 3. CompTox: HTTK (High-Throughput Toxicokinetics)
    print("3. CompTox: Extracting Predicted Kinetic Parameters...")
    httk = mcp_call("CompTox", "search_httk", {"dtxsid": dtxsid})
    save_artifact(chem_key, "comptox_httk.json", httk)
    
    # 4. PBPK: Detailed Organ-Level Exposure (Proxy)
    print("4. PBPK: Quantifying Brain vs. Blood Exposure (Proxy Envelope)...")
    sim_id = "Acetaminophen_Pregnancy"
    
    # Ensure fresh session loading
    print("   (Ensuring simulation is loaded...)")
    
    # Stage file to var/ for container access
    import shutil
    import os
    os.makedirs("var", exist_ok=True)
    shutil.copy("models/acetaminophen/Acetaminophen_Pregnancy.pkml", "var/Acetaminophen_Pregnancy.pkml")
    
    mcp_call("PBPK", "load_simulation", {"filePath": "var/Acetaminophen_Pregnancy.pkml", "simulationId": sim_id})
    
    run_resp = mcp_call("PBPK", "run_simulation", {"simulationId": sim_id})
    
    if run_resp and "jobId" in run_resp:
        job_id = run_resp.get("jobId")
        while True:
            status = mcp_call("PBPK", "get_job_status", {"jobId": job_id})
            if status["job"]["status"] == "succeeded":
                result_id = status["job"].get("resultId")
                break
            time.sleep(0.5)
            
        pk_resp = mcp_call("PBPK", "calculate_pk_parameters", {"resultsId": result_id})
        save_artifact(chem_key, "pbpk_metrics.json", pk_resp)

    # 5. AOP: Pathological Mechanism Mapping
    print("5. AOP: Mapping Molecular Initiating Events to Adverse Outcomes...")
    get_best_aop_mapping(chem_key, chem_name, cas, inchikey)
    
    # 6. O-QT: Advanced QSAR & Profiling
    print("6. O-QT: Running Agentic QSAR Workflow...")
    oqt_workflow = mcp_call("O-QT", "run_oqt_multiagent_workflow", {
        "identifier": chem_name,
        "profiler_guids": ["aa645923-a592-46fe-9069-737a2e4a7ac6"], # Cramer
        "qsar_mode": "recommended"
    })
    save_artifact(chem_key, "oqt_report.json", oqt_workflow)
    
    print("\nCase Study 1 Complete.")

if __name__ == "__main__":
    run_case_study_1_nature()
```
