## Supplementary material for "ToxMCP: Guardrailed, Auditable Agentic Workflows for Computational Toxicology via the Model Context Protocol": run_cs2_acetaminophen.py

```
from full_MCP_Suite.mcp_utils import mcp_call, save_artifact, get_best_aop_mapping
import time

def run_case_study_2_nature():
    print("=== Case Study 2: Acetaminophen (Nature Publication Quality) ===")
    chem_name = "Acetaminophen"
    chem_key = "acetaminophen"
    
    # 1. CompTox: Identity
    print("1. CompTox: Resolving Identity...")
    chem_search = mcp_call("CompTox", "search_chemical", {"query": chem_name, "search_type": "equals"})
    
    dtxsid = None
    inchikey = None
    cas = None
    
    if chem_search and "data" in chem_search:
        for hit in chem_search["data"]:
            if hit.get("preferredName").lower() == chem_name.lower():
                dtxsid = hit.get("dtxsid")
                cas = hit.get("casrn")
                break
    
    if dtxsid:
        details = mcp_call("CompTox", "get_chemical_details", {"identifier": dtxsid, "id_type": "dtxsid"})
        if details:
            inchikey = details.get("inchikey")
            save_artifact(chem_key, "comptox_details.json", details)

    # 2. CompTox: NAM Point of Departure (AED)
    print("2. CompTox: Retrieving NAM-based Point of Departure (AED)...")
    aed_res = mcp_call("CompTox", "get_bioactivity_aed", {"dtxsid": dtxsid})
    save_artifact(chem_key, "comptox_aed.json", aed_res)

    # 3. PBPK: Detailed Mechanistic Simulation
    print("3. PBPK: Quantifying Dynamic Exposure in Pregnancy...")
    sim_id = "Acetaminophen_Pregnancy"
    
    # Ensure fresh session loading
    print("   (Ensuring simulation is loaded...)")
    
    # Stage file to var/ for container access
    import shutil
    import os
    os.makedirs("var", exist_ok=True)
    shutil.copy("models/acetaminophen/Acetaminophen_Pregnancy.pkml", "var/Acetaminophen_Pregnancy.pkml")
    
    mcp_call("PBPK", "load_simulation", {"filePath": "var/Acetaminophen_Pregnancy.pkml", "simulationId": sim_id})
    
    run_resp = mcp_call("PBPK", "run_simulation", {"simulationId": sim_id})
    
    if run_resp and "jobId" in run_resp:
        job_id = run_resp.get("jobId")
        while True:
            status = mcp_call("PBPK", "get_job_status", {"jobId": job_id})
            if status["job"]["status"] == "succeeded":
                result_id = status["job"].get("resultId")
                break
            time.sleep(0.5)
            
        pk_resp = mcp_call("PBPK", "calculate_pk_parameters", {"resultsId": result_id})
        save_artifact(chem_key, "pbpk_metrics.json", pk_resp)

    # 4. PBPK: Sensitivity Analysis
    print("4. PBPK: Analyzing Sensitivity to Liver Volume...")
    sens_resp = mcp_call("PBPK", "run_sensitivity_analysis", {
        "modelPath": "var/Acetaminophen_Pregnancy.pkml",
        "simulationId": sim_id,
        "parameters": [{"path": "Organism|Liver|Volume", "deltas": [-0.2, 0.2]}],
        "metrics": ["AUC_Plasma"]
    })
    save_artifact(chem_key, "pbpk_sensitivity.json", sens_resp)

    # 5. AOP: Mapping
    print("5. AOP: Mapping to Liver Injury Pathways...")
    get_best_aop_mapping(chem_key, chem_name, cas, inchikey)

    print("\nCase Study 2 Complete.")

if __name__ == "__main__":
    run_case_study_2_nature()
```
