## Supplementary material for "ToxMCP: Guardrailed, Auditable Agentic Workflows for Computational Toxicology via the Model Context Protocol": run_cs2_part2.py

```
import json
import time
import requests
import os

SERVERS = {
    "PBPK": "http://localhost:8000/mcp",
    "CompTox": "http://localhost:8001/mcp",
    "AOP": "http://localhost:803/mcp"
}

def mcp_call(server_name, tool_name, arguments):
    url = SERVERS[server_name]
    payload = {
        "jsonrpc": "2.0",
        "method": "tools/call",
        "params": {
            "name": tool_name,
            "arguments": arguments
        },
        "id": int(time.time() * 1000)
    }
    response = requests.post(url, json=payload)
    response.raise_for_status()
    data = response.json()
    if "error" in data and data["error"]:
        print(f"Error calling {server_name}/{tool_name}: {data['error']}")
        return None
    
    result = data.get("result", {})
    if "structuredContent" in result:
        return result["structuredContent"]
    if "content" in result:
        for item in result["content"]:
            if item["type"] == "text":
                try:
                    return json.loads(item["text"])
                except:
                    return item["text"]
    return result

def save_artifact(chemical, filename, data):
    base_dir = f"full_MCP_Suite/bench_runs/chemicals/{chemical}"
    os.makedirs(base_dir, exist_ok=True)
    path = os.path.join(base_dir, filename)
    with open(path, "w") as f:
        json.dump(data, f, indent=2)
    print(f"Saved: {path}")

def run_case_study_2_part2():
    print("=== Case Study 2 (Part 2): Acetaminophen Sensitivity & AOP ===")
    chem_key = "acetaminophen"
    sim_id = "Acetaminophen_Pregnancy"
    
    # We know the InChIKey from previous run: RZVAJINKPMORJF-UHFFFAOYSA-N
    inchikey = "RZVAJINKPMORJF-UHFFFAOYSA-N"

    # 3. PBPK: Liver Sensitivity (Re-running simplified version for dossier)
    print("3. PBPK: Running Liver Sensitivity (Automated)...")
    
    # Check if sim needs loading? It should stay loaded in the server unless restarted.
    # We'll try running sensitivity directly.
    
    sens_resp = mcp_call("PBPK", "run_sensitivity_analysis", {
        "modelPath": "var/Acetaminophen_Pregnancy.pkml",
        "simulationId": sim_id,
        "parameters": [{"path": "Organism|Liver|Volume", "deltas": [-0.2, 0.2]}],
        "metrics": ["AUC_Plasma"]
    })
    
    if sens_resp:
        save_artifact(chem_key, "pbpk_sensitivity.json", sens_resp)
    else:
        print("Sensitivity Analysis failed (likely simulation not loaded or server error).")

    # 4. AOP: Mapping
    print("4. AOP: Mapping to pathways...")
    if inchikey:
        aop_map = mcp_call("AOP", "map_chemical_to_aops", {"inchikey": inchikey})
        save_artifact(chem_key, "aop_map.json", aop_map)
    else:
        print("Skipping AOP (No InChIKey).")

    print("\nCase Study 2 Complete (Repaired).")

if __name__ == "__main__":
    run_case_study_2_part2()
```
