## Supplementary material for "ToxMCP: Guardrailed, Auditable Agentic Workflows for Computational Toxicology via the Model Context Protocol": run_cs3_triclosan.py

```
from full_MCP_Suite.mcp_utils import mcp_call, save_artifact, get_best_aop_mapping
import time

def run_case_study_3_nature():
    print("=== Case Study 3: Triclosan (Nature Publication Quality) ===")
    chem_name = "Triclosan"
    chem_key = "triclosan"
    
    # 1. CompTox: Identity
    print("1. CompTox: Resolving Identity...")
    chem_search = mcp_call("CompTox", "search_chemical", {"query": chem_name, "search_type": "equals"})
    
    dtxsid = None
    inchikey = None
    cas = None
    
    if chem_search and "data" in chem_search:
        for hit in chem_search["data"]:
            if hit.get("preferredName").lower() == chem_name.lower():
                dtxsid = hit.get("dtxsid")
                cas = hit.get("casrn")
                break

    if dtxsid:
        details = mcp_call("CompTox", "get_chemical_details", {"identifier": dtxsid, "id_type": "dtxsid"})
        if details:
            inchikey = details.get("inchikey")
            save_artifact(chem_key, "comptox_details.json", details)
            
        # 2. CompTox: NAM PoD
        print("2. CompTox: Retrieving NAM Point of Departure (AED)...")
        aed_res = mcp_call("CompTox", "get_bioactivity_aed", {"dtxsid": dtxsid})
        save_artifact(chem_key, "comptox_aed.json", aed_res)

        # 3. CompTox: Exposure (Consumer Products)
        print("3. CompTox: Quantifying Consumer Product Usage...")
        cpdat = mcp_call("CompTox", "search_cpdat", {"dtxsid": dtxsid, "vocab_name": "puc"})
        save_artifact(chem_key, "comptox_cpdat.json", cpdat)
        
    # 4. AOP: Endocrine/Thyroid Mechanism Mapping
    print("4. AOP: Mapping Endocrine Disruption Mechanisms...")
    get_best_aop_mapping(chem_key, chem_name, cas, inchikey)

    print("\nCase Study 3 Complete.")

if __name__ == "__main__":
    run_case_study_3_nature()
```
