## Supplementary material for "ToxMCP: Guardrailed, Auditable Agentic Workflows for Computational Toxicology via the Model Context Protocol": test_mcps.py

```
import json
import requests
import sys

SERVERS = {
    "PBPK": "http://localhost:8000/mcp",
    "CompTox": "http://localhost:8001/mcp",
    "AOP": "http://localhost:803/mcp",
    "O-QT": "http://localhost:804/mcp"
}

def check_server(name, url):
    print(f"Checking {name} ({url})...", end=" ")
    try:
        # Initialize
        init_payload = {
            "jsonrpc": "2.0",
            "method": "initialize",
            "params": {
                "capabilities": {},
                "clientInfo": {"name": "test_suite", "version": "1.0"}
            },
            "id": 1
        }
        resp = requests.post(url, json=init_payload, timeout=5)
        resp.raise_for_status()
        data = resp.json()
        
        # PBPK returns error: null in success case, which my previous script misread
        if "error" in data and data["error"] is not None:
            print(f"Error: {data['error']}")
            return False
            
        print("✓ Connected.")
        
        # List tools
        tools_payload = {
            "jsonrpc": "2.0",
            "method": "tools/list",
            "params": {},
            "id": 2
        }
        resp = requests.post(url, json=tools_payload, timeout=5)
        tools_data = resp.json()
        
        tools = tools_data.get("result", {}).get("tools", [])
        print(f"  Tools found: {len(tools)}")
        return True
        
    except Exception as e:
        print(f"FAILED: {e}")
        return False

def main():
    print("=== Multi-MCP Suite Health Check ===\n")
    results = {}
    for name, url in SERVERS.items():
        results[name] = check_server(name, url)
        print("")
        
    print("--- Summary ---")
    all_pass = True
    for name, passed in results.items():
        status = "PASS" if passed else "FAIL"
        print(f"{name}: {status}")
        if not passed:
            all_pass = False
            
    if all_pass:
        print("\nAll systems operational. Ready for benchmark.")
        sys.exit(0)
    else:
        print("\nSome systems failed. Check logs.")
        sys.exit(1)

if __name__ == "__main__":
    main()
```
