## Supplementary material for "ToxMCP: Guardrailed, Auditable Agentic Workflows for Computational Toxicology via the Model Context Protocol": MANIFEST.md

# Benchmark Suite Manifest

This repository contains the artifacts, code, and data for the Multi-MCP Toxicology Benchmark.

## Directory Structure

### `models/` (Input Truth)
Contains the canonical, hash-verified PBPK model files.
- `acetaminophen/Acetaminophen_Pregnancy.pkml`
- `midazolam/Midazolam_Canonical.pkml`
- `caffeine/Caffeine_Canonical.pkml`

### `publication_artifacts/` (Output Truth)
Contains the frozen outputs from the benchmark run.
- `benchmark_summary.csv` / `.md`: The final aggregated results table.
- `figures/`: Generated publication figures (Sensitivity, Heatmap, Scatter).
- `acetaminophen/`, `chlorpyrifos/`, etc.: Per-chemical "Dossier Bundles" containing raw JSON responses from all MCPs.
- `pbpk_outputs/`: Raw outputs from the PBPK server execution.

### `full_MCP_Suite/` (Code)
- `run_benchmark_batch.py`: Script to execute the 12-chemical automated benchmark.
- `run_cs{1-3}_*.py`: Deep case study scripts.
- `generate_benchmark_summary.py`: Aggregation logic.
- `generate_publication_figures.py`: Visualization logic.

### Root Scripts
- `run_benchmark.py`: The PBPK-specific deterministic replay harness.
- `expected_metrics.json`: The "Golden Manifest" defining the PBPK ground truth (hashes + tolerances).

## Reproducibility
To reproduce the PBPK metrics:
1. Ensure the PBPK MCP server is running.
2. Run `python3 run_benchmark.py` (Validates hashes, loads models, replays simulations).

To reproduce the Tox Dashboard:
1. Ensure CompTox, AOP, O-QT, and ADMET MCPs are running.
2. Run `python3 -m full_MCP_Suite.run_benchmark_batch`.
3. Run `python3 full_MCP_Suite/generate_benchmark_summary.py`.
