## Supplementary material for "ToxMCP: Guardrailed, Auditable Agentic Workflows for Computational Toxicology via the Model Context Protocol": plot_benchmark_results.py

```
import json
import matplotlib.pyplot as plt
import pandas as pd
import seaborn as sns
import numpy as np
import os

# Configuration
RESULTS_FILE = "benchmark_results.json"
OUTPUT_DIR = "figures"
HEATMAP_FILENAME = "Figure_3_PBPK_ReplayHeatmap.pdf"
BOXPLOT_FILENAME = "Figure_3_PBPK_ReplayBoxplots.pdf"

# Ensure output directory exists
os.makedirs(OUTPUT_DIR, exist_ok=True)

def generate_dummy_data():
    """Generates dummy benchmark results for demonstration."""
    print(f"Warning: {RESULTS_FILE} not found. Generating dummy data for demonstration.")
    data = []
    cases = ["Acetaminophen", "Midazolam", "Caffeine", "Buprenorphine"]
    metrics = ["AUC_Plasma", "CMax_Plasma", "TMax_Plasma"]
    
    for case in cases:
        for run in range(1, 11): # 10 runs
            for metric in metrics:
                # Simulate small random deviations
                deviation = np.random.normal(0, 0.05) # Mean 0, SD 0.05%
                data.append({
                    "case_id": case,
                    "run_id": run,
                    "metric": metric,
                    "deviation_percent": abs(deviation)
                })
    return pd.DataFrame(data)

def load_data():
    if not os.path.exists(RESULTS_FILE):
        return generate_dummy_data()
    
    with open(RESULTS_FILE, 'r') as f:
        raw_data = json.load(f)
    
    # Handle new metadata wrapper
    if isinstance(raw_data, dict) and "rows" in raw_data:
        return pd.DataFrame(raw_data["rows"])

    # Check if data is already in flat format (list of dicts with 'deviation_percent')
    if isinstance(raw_data, list) and raw_data and "deviation_percent" in raw_data[0]:
        return pd.DataFrame(raw_data)

    # Fallback legacy/nested structure logic (removed for clarity as we control the producer)
    print("Unknown JSON format. Using dummy data.")
    return generate_dummy_data()

def plot_heatmap(df):
    """Generates Figure 3A: Replay Error Heatmap (Max Deviation)."""
    # Aggregate: Max deviation per (Case, Metric)
    pivot_df = df.groupby(["case_id", "metric"])["deviation_percent"].max().unstack()
    
    plt.figure(figsize=(10, 8)) # Increased size for labels
    sns.set_theme(style="white")
    
    # Create heatmap
    ax = sns.heatmap(
        pivot_df, 
        annot=True, 
        fmt=".2e", # Scientific notation
        cmap="Reds", 
        linewidths=.5, 
        cbar_kws={'label': 'Max Absolute Deviation (%)'},
        vmin=0, vmax=1.0 # Anchor scale for "tolerance" visualization
    )
    
    plt.title("PBPK Engine Replayability: Max Deviation across Replays", pad=20)
    plt.ylabel("Model")
    plt.xlabel("PK Metric")
    plt.tight_layout()
    
    output_path_pdf = os.path.join(OUTPUT_DIR, HEATMAP_FILENAME)
    output_path_png = os.path.join(OUTPUT_DIR, HEATMAP_FILENAME.replace(".pdf", ".png"))
    
    plt.savefig(output_path_pdf)
    plt.savefig(output_path_png, dpi=300)
    print(f"Saved Heatmap to {output_path_pdf} and {output_path_png}")
    plt.close()

def plot_boxplots(df):
    """Generates Figure 3B: Distribution of Deviations."""
    plt.figure(figsize=(10, 6))
    sns.set_theme(style="ticks")
    
    # Boxplot
    ax = sns.boxplot(
        data=df,
        x="metric",
        y="deviation_percent",
        hue="case_id",
        palette="vlag"
    )
    
    # Add tolerance line
    plt.axhline(y=1.0, color='r', linestyle='--', label='Tolerance (1%)')
    
    # Use symlog to show small deviations while handling zero
    plt.yscale('symlog', linthresh=1e-6)
    
    plt.title("Distribution of Deterministic Replay Deviations", pad=20)
    plt.ylabel("Absolute Deviation (%) - Log Scale", fontsize=10)
    plt.xlabel("PK Metric")
    plt.legend(title="Model", loc='upper right')
    sns.despine(offset=10, trim=True)
    plt.tight_layout()
    
    output_path_pdf = os.path.join(OUTPUT_DIR, BOXPLOT_FILENAME)
    output_path_png = os.path.join(OUTPUT_DIR, BOXPLOT_FILENAME.replace(".pdf", ".png"))
    
    plt.savefig(output_path_pdf)
    plt.savefig(output_path_png, dpi=300)
    print(f"Saved Boxplots to {output_path_pdf} and {output_path_png}")
    plt.close()

def main():
    print("Loading benchmark results...")
    df = load_data()
    
    print("Generating figures...")
    plot_heatmap(df)
    plot_boxplots(df)
    
    print("Done.")

if __name__ == "__main__":
    main()
```
