## Supplementary material for "ToxMCP: Guardrailed, Auditable Agentic Workflows for Computational Toxicology via the Model Context Protocol": benchmark_summary.md

| Chemical      | CAS        | NAM_PoD_AED      | HTTK_Css_Median   | PBPK_Brain_Blood_Ratio   | AOP_Mapping                                                 | QSAR_Cramer_Class       |   Product_Count |
|:--------------|:-----------|:-----------------|:------------------|:-------------------------|:------------------------------------------------------------|:------------------------|----------------:|
| Acetaminophen | 103-90-2   | 56.99 mg/kg/day  | 0.003538 mg/L     | 0.982                    | 1 AOPs (e.g. CYP2E1 activation and formation of protein...) | High (Class III)        |             149 |
| Atrazine      | 1912-24-9  | 0.1258 mg/kg/day | 0.03152 mg/L      | N/A                      | 4 AOPs (e.g. GnRH pulse disruption leading to mammary a...) | High (Class III)        |             114 |
| Bisphenol_a   | 80-05-7    | 0.4274 mg/kg/day | 0.5297 mg/L       | N/A                      | 4 AOPs (e.g. Binding to estrogen receptor (ER)-α in imm...) | High (Class III)        |             326 |
| Caffeine      | 58-08-2    | 25.85 mg/kg/day  | 0.0239 mg/L       | N/A                      | None                                                        | Intermediate (Class II) |             933 |
| Carbamazepine | 298-46-4   | 3.071 mg/kg/day  | 0.04942 mg/L      | N/A                      | None                                                        | High (Class III)        |               1 |
| Chlorpyrifos  | 2921-88-2  | 1.343 mg/kg/day  | 0.0265 mg/L       | 0.982                    | 3 AOPs (e.g. Inhibitor binding to topoisomerase II lead...) | High (Class III)        |             161 |
| Endosulfan    | 115-29-7   | 0.2216 mg/kg/day | 1.918 mg/L        | N/A                      | 2 AOPs (e.g. Binding to the picrotoxin site of ionotrop...) | High (Class III)        |               3 |
| Glyphosate    | 1071-83-6  | N/A              | N/A               | N/A                      | None                                                        | High (Class III)        |              37 |
| Oxybenzone    | 131-57-7   | 9.433 mg/kg/day  | 0.1719 mg/L       | N/A                      | None                                                        | High (Class III)        |            3804 |
| Silychristin  | 33889-69-9 | N/A              | N/A               | N/A                      | None                                                        | High (Class III)        |               0 |
| Triclosan     | 3380-34-5  | 2.974 mg/kg/day  | 0.1731 mg/L       | N/A                      | 2 AOPs (e.g. Uncoupling of oxidative phosphorylation le...) | High (Class III)        |             266 |
| Warfarin      | 81-81-2    | 0.5872 mg/kg/day | 7.87 mg/L         | N/A                      | 1 AOPs (e.g. Anticoagulant rodenticide inhibition of vi...) | High (Class III)        |              15 |
