## Supplementary material for "ToxMCP: Guardrailed, Auditable Agentic Workflows for Computational Toxicology via the Model Context Protocol": README.md

# Multi-MCP Toxicology Benchmark Artifacts

This directory contains the reproducible outputs for the Multi-MCP Toxicology Benchmark study.

## Structure

- **benchmark_summary.csv / .md**: Aggregated summary table of the 12-chemical automated benchmark.
- **figures/**: Generated publication-quality figures.
    - `Figure_1_PBPK_Sensitivity.png`
    - `Figure_2_Data_Completeness.png`
    - `Figure_3_PoD_vs_HTTK.png`
    - `Figure_5_PBPK_Determinism_Proof.png`
- **{chemical_name}/**: Dossier folder for each chemical containing raw JSON artifacts from the MCPs.

## Chemicals Benchmarked
1. Acetaminophen
2. Atrazine
3. Bisphenol A
4. Caffeine
5. Carbamazepine
6. Chlorpyrifos
7. Endosulfan
8. Glyphosate
9. Oxybenzone
10. Silychristin
11. Triclosan
12. Warfarin

## Chemical Dossier Contents

Each chemical folder contains provenance-tracked JSON responses from the MCP agents:

- `comptox_identity.json`: Chemical identification and keys (DTXSID, CAS, InChIKey).
- `comptox_aed.json`: NAM-based Point of Departure (Bioactivity).
- `comptox_httk.json`: High-Throughput Toxicokinetics predictions (Css, Half-life).
- `comptox_cpdat.json`: Consumer Product Database records (Exposure potential).
- `aop_map.json`: Mapped Adverse Outcome Pathways (AOP-Wiki).
- `pbpk_metrics.json`: (Case Studies only) Deterministic PBPK simulation metrics.
- `pbpk_sensitivity.json`: (Case Studies only) Sensitivity analysis results.
- `oqt_report.json`: (Case Studies only) Full QSAR Toolbox agentic workflow logs.
- `oqt_cramer.json`: (All chemicals) Cramer Rules safety profiling result.

## Reproduction

These artifacts were generated by:
1. `full_MCP_Suite.run_cs{1-3}_{name}` (Deep Case Studies)
2. `full_MCP_Suite.run_benchmark_batch` (12-Chemical Automation with O-QT Cramer Profiling)
3. `full_MCP_Suite.generate_benchmark_summary` (Aggregation)
4. `full_MCP_Suite.generate_publication_figures` (Visualization)

## System Configuration
Ports and endpoints are deployment-specific. The scripts assume localhost defaults unless configured otherwise.
