## Supplementary material for "ToxMCP: Guardrailed, Auditable Agentic Workflows for Computational Toxicology via the Model Context Protocol": README.md

# ToxMCP Evidence Bundle — Benchmark Supplement (Supplementary Archive S1)

This archive contains the *frozen artifacts* (JSON tool outputs, summary tables, figures) and *replay utilities* used in the
ToxMCP manuscript case studies and the 12-chemical multi-MCP benchmark.

> Intended use: readers/reviewers can (i) inspect raw evidence objects, (ii) regenerate the manuscript figures/tables from cached
> artifacts, and (iii) (optionally) re-run PBPK determinism checks against a running PBPK MCP server.

## Top-level contents

- `MANIFEST.md` — human-readable overview of the directory structure and reproduction entrypoints.
- `MANIFEST.sha256` — cryptographic hashes for **all files** in this archive (integrity / tamper-evidence).
- `expected_metrics.json` — "gold" PBPK regression targets (model file hashes + expected PK metrics + tolerance).
- `run_benchmark.py` — PBPK deterministic replay harness (requires PBPK MCP server).
- `plot_benchmark_results.py` — plotting helper for PBPK replay results.
- `publication_artifacts/` — frozen outputs that back the manuscript tables/figures.
- `models/` — canonical PBPK model files used by the PBPK replay harness.
- `full_MCP_Suite/` — reference scripts used to generate/inspect artifacts (some require multiple MCP servers).

## Reproduction modes

### A) Offline reproduction (no MCP servers)
You can regenerate the publication figures from the cached artifacts in `publication_artifacts/` using:
- `python3 MCP_Benchmark_Supplementary/full_MCP_Suite/generate_publication_figures.py`

### B) PBPK determinism replay (requires PBPK MCP server)
1. Start your PBPK MCP server (default endpoint assumed by `run_benchmark.py`):
   - `http://localhost:8000/mcp`
2. Run:
   - `python3 MCP_Benchmark_Supplementary/run_benchmark.py`
3. Optional: plot results
   - `python3 MCP_Benchmark_Supplementary/plot_benchmark_results.py`

### C) Full multi-MCP benchmark re-run (requires multiple MCP servers)
Re-running the 12-chemical benchmark requires CompTox, AOP, ADMETlab, and O-QT MCP servers in addition to PBPK.
See `full_MCP_Suite/full_MCP.md` for the expected tool surface and scripts.

## Integrity verification
To verify file integrity:
- Compare `MANIFEST.sha256` against your local copy after download/unzip.
- The PBPK model hashes are also redundantly verified via `expected_metrics.json` by `run_benchmark.py`.


## Licensing and third-party terms

Licensing note: This archive redistributes only (i) derived result artifacts (e.g., exported JSON summaries / report artifacts) produced using third-party scientific tools and services, and (ii) PBPK model fixtures in PKML format used for deterministic replay. It does **not** include the OECD QSAR Toolbox installer/executables or any donated databases/modules covered by the QSAR Toolbox End User Licence Agreement (EULA); reproducing QSAR workflows therefore requires users to separately download and use QSAR Toolbox under its EULA. PBPK simulations are executed with the Open Systems Pharmacology Suite (PK-Sim®/MoBi®; GPLv2); this archive does not ship OSP binaries and assumes users install OSP separately. If upstream data sources impose additional use terms, those terms remain applicable to the underlying content.
## Notes on identity artifacts
The 12-chemical benchmark folders include `comptox_identity.json` as a **derived identity summary** extracted from the frozen
`comptox_details.json` responses. This makes canonical identifiers explicit without requiring an additional live CompTox lookup.

---
Generated/packaged for preprint submission as Supplementary Archive S1.
