## Supplementary material for "ToxMCP: Guardrailed, Auditable Agentic Workflows for Computational Toxicology via the Model Context Protocol": run_benchmark.py

```
import json
import hashlib
import time
import os
import requests
import sys

# Configuration
MCP_URL = "http://localhost:8000/mcp"
EXPECTED_METRICS_FILE = "expected_metrics.json"
RESULTS_FILE = "benchmark_results.json"
NUM_REPLAYS = 5
POLL_INTERVAL = 1.0

def calculate_sha256(filepath):
    """Calculates the SHA256 hash of a file."""
    if not os.path.exists(filepath):
        return None
    sha256_hash = hashlib.sha256()
    with open(filepath, "rb") as f:
        for byte_block in iter(lambda: f.read(4096), b""):
            sha256_hash.update(byte_block)
    return sha256_hash.hexdigest()

def mcp_call(tool_name, arguments):
    """Helper to call MCP tools via HTTP."""
    payload = {
        "jsonrpc": "2.0",
        "method": "tools/call",
        "params": {
            "name": tool_name,
            "arguments": arguments
        },
        "id": int(time.time() * 1000)
    }
    response = requests.post(MCP_URL, json=payload)
    # print(f"DEBUG: {tool_name} response: {response.text}") # Uncomment for verbose debug
    response.raise_for_status()
    data = response.json()
    
    if data.get("error"):
        return {"error": data["error"]}
    
    # Parse the content text if it's a JSON string inside the result
    if "result" in data and data["result"] and "content" in data["result"]:
        for item in data["result"]["content"]:
            if item["type"] == "text":
                try:
                    return json.loads(item["text"])
                except json.JSONDecodeError:
                    return item["text"]
    return data.get("result")

def run_benchmark():
    print(f"Loading configuration from {EXPECTED_METRICS_FILE}...")
    with open(EXPECTED_METRICS_FILE, "r") as f:
        config = json.load(f)

    results = []

    for case in config["cases"]:
        case_id = case["id"]
        model_name = case["name"]
        model_filename = case["modelPath"]
        expected_hash = case["sha256"]
        expected_metrics = case["expectedMetrics"]
        
        # Determine Simulation ID (filename without extension)
        simulation_id = os.path.splitext(os.path.basename(model_filename))[0]
        
        # Staging Logic for Docker/Container:
        # We try staging directly to 'var/' as we know that worked previously.
        
        import shutil
        staging_dir = "var"
        os.makedirs(staging_dir, exist_ok=True)
        staging_path = os.path.join(staging_dir, os.path.basename(model_filename))
        
        # Copy from source (models/...) to staging (var/...)
        shutil.copy(model_filename, staging_path)
        
        # MCP sees 'var/...' relative to its root
        mcp_file_path = staging_path
        local_file_path = model_filename

        print(f"\n--- Processing Case: {model_name} ({case_id}) ---")
        
        # 1. Verify File (Local check skipped if running against containerized var/)
        if os.path.exists(local_file_path):
            current_hash = calculate_sha256(local_file_path)
            if expected_hash != "PENDING_CONVERSION" and current_hash != expected_hash:
                 print(f"⚠️  Warning: Hash mismatch for {local_file_path}.")
                 print(f"    Expected: {expected_hash}")
                 print(f"    Found:    {current_hash}")
        else:
            print(f"ℹ️  Local file '{local_file_path}' not found, assuming it exists in server '{mcp_file_path}'.")
        
        # 2. Load Simulation
        print(f"Loading simulation '{simulation_id}' from '{mcp_file_path}'...")
        load_resp = mcp_call("load_simulation", {"filePath": mcp_file_path})
        
        if isinstance(load_resp, dict) and "error" in load_resp:
            # Check if it's just "already registered"
            err_msg = load_resp["error"].get("message", "")
            if "already registered" in err_msg:
                print("    (Simulation already loaded, proceeding)")
            else:
                print(f"❌ Failed to load simulation: {err_msg}")
                continue

        # 3. Replay Loop
        for i in range(1, NUM_REPLAYS + 1):
            print(f"  Run {i}/{NUM_REPLAYS}...", end="", flush=True)
            
            # Start Run
            run_resp = mcp_call("run_simulation", {"simulationId": simulation_id})
            
            if run_resp is None:
                print(" Failed: Empty response from tool.")
                continue

            if "error" in run_resp:
                 print(f" Failed to start: {run_resp['error'].get('message', 'Unknown error')}")
                 continue
            
            job_id = run_resp.get("jobId")
            if not job_id:
                print(" Failed: No Job ID returned.")
                continue
                
            # Poll Status
            while True:
                status_resp = mcp_call("get_job_status", {"jobId": job_id})
                job_status = status_resp.get("job", {}).get("status")
                
                if job_status == "succeeded":
                    result_id = status_resp["job"].get("resultId")
                    break
                elif job_status in ["failed", "cancelled"]:
                    job_error = status_resp.get("job", {}).get("error")
                    print(f" Job {job_status}. Error: {job_error}")
                    result_id = None
                    break
                
                time.sleep(POLL_INTERVAL)
            
            if not result_id:
                continue

            # Calculate Parameters
            pk_resp = mcp_call("calculate_pk_parameters", {"resultsId": result_id})
            if "error" in pk_resp:
                print(f" Failed to calc PK: {pk_resp['error']['message']}")
                continue
            
            # Extract Metrics
            observed_values = {}
            target_path = case.get("parameterPath")
            
            for metric_entry in pk_resp.get("metrics", []):
                # If a specific path is configured, use strict matching
                if target_path:
                    if metric_entry["parameter"] == target_path:
                        observed_values["AUC_Plasma"] = metric_entry["auc"]
                        observed_values["CMax_Plasma"] = metric_entry["cmax"]
                        observed_values["TMax_Plasma"] = metric_entry["tmax"]
                        break # Found our target
                
                # Fallback to loose matching if no path specified (legacy behavior)
                elif "Plasma" in metric_entry["parameter"] and "Concentration" in metric_entry["parameter"]:
                     # Try to avoid metabolites if possible, but this is brittle
                     if "glucuronide" not in metric_entry["parameter"] and "sulfate" not in metric_entry["parameter"]:
                        observed_values["AUC_Plasma"] = metric_entry["auc"]
                        observed_values["CMax_Plasma"] = metric_entry["cmax"]
                        observed_values["TMax_Plasma"] = metric_entry["tmax"]
            
            # Compute Deviations
            for key, expected_val in expected_metrics.items():
                if expected_val == 0.0: continue # Skip if no expectation set
                
                observed = observed_values.get(key)
                if observed is not None:
                    dev_percent = abs(observed - expected_val) / expected_val * 100
                    
                    results.append({
                        "case_id": case_id,
                        "run_id": i,
                        "metric": key,
                        "observed": observed,
                        "expected": expected_val,
                        "deviation_percent": dev_percent
                    })
            
            print(" Done.")

    # Wrap results with metadata
    output_data = {
        "metadata": {
            "timestamp_utc": time.strftime("%Y-%m-%dT%H:%M:%SZ", time.gmtime()),
            "num_replays": NUM_REPLAYS,
            "tolerance_percent": config.get("tolerancePercent", 1.0),
            "engine_type": "PBPK MCP"
        },
        "rows": results
    }

    print(f"\nSaving results to {RESULTS_FILE}...")
    with open(RESULTS_FILE, "w") as f:
        json.dump(output_data, f, indent=2)
    print("Benchmark completed.")

if __name__ == "__main__":
    run_benchmark()
```
