## Supplementary material for "ToxMCP: Guardrailed, Auditable Agentic Workflows for Computational Toxicology via the Model Context Protocol": validate_bundle.py

```
#!/usr/bin/env python3
import argparse
import csv
import hashlib
import io
import json
import os
import sys
from pathlib import Path

REQUIRED_PER_CHEM = [
    "comptox_details.json",
    "comptox_identity.json",
    "comptox_aed.json",
    "comptox_httk.json",
    "comptox_cpdat.json",
    # AOP can be either populated or explicitly empty
    # QSAR (Cramer) always present in this bundle
    "oqt_cramer.json",
]

AOP_OK = {"aop_map.json", "aop_map_empty.json"}


def sha256_file(path: Path) -> str:
    h = hashlib.sha256()
    with path.open("rb") as f:
        for chunk in iter(lambda: f.read(1024 * 1024), b""):
            h.update(chunk)
    return h.hexdigest()


def load_json(path: Path):
    with path.open("r", encoding="utf-8") as f:
        return json.load(f)


def main() -> int:
    ap = argparse.ArgumentParser(description="Validate the MCP_Benchmark_Supplementary evidence bundle.")
    ap.add_argument("--bundle", type=str, default=".", help="Path to MCP_Benchmark_Supplementary directory (extracted).")
    args = ap.parse_args()

    root = Path(args.bundle).resolve()
    if root.name != "MCP_Benchmark_Supplementary":
        # allow pointing to parent directory
        if (root / "MCP_Benchmark_Supplementary").exists():
            root = (root / "MCP_Benchmark_Supplementary").resolve()

    ok = True

    # 1) Expected files exist
    for p in ["expected_metrics.json", "MANIFEST.md", "MANIFEST.sha256", "publication_artifacts/benchmark_summary.csv"]:
        if not (root / p).exists():
            print(f"[FAIL] Missing required file: {p}")
            ok = False

    # 2) JSON parse sanity
    for jf in root.rglob("*.json"):
        try:
            load_json(jf)
        except Exception as e:
            print(f"[FAIL] JSON parse error: {jf.relative_to(root)} :: {e}")
            ok = False

    # 3) Model hashes match expected_metrics
    try:
        exp = load_json(root / "expected_metrics.json")
        tol = exp.get("tolerancePercent", None)
        cases = exp.get("cases", [])
        if tol is None or not isinstance(cases, list):
            raise ValueError("expected_metrics.json does not match expected schema (tolerancePercent, cases[])")
        for c in cases:
            mp = c["modelPath"]
            expected_sha = c["sha256"]
            fp = root / mp
            if not fp.exists():
                print(f"[FAIL] Model file missing: {mp}")
                ok = False
                continue
            actual_sha = sha256_file(fp)
            if actual_sha != expected_sha:
                print(f"[FAIL] Model hash mismatch: {mp}")
                print(f"       expected: {expected_sha}")
                print(f"       actual:   {actual_sha}")
                ok = False
        if ok:
            print("[PASS] PBPK model hashes match expected_metrics.json")
    except Exception as e:
        print(f"[FAIL] Could not validate expected_metrics.json :: {e}")
        ok = False

    # 4) Benchmark summary CSV sanity
    summary_path = root / "publication_artifacts" / "benchmark_summary.csv"
    try:
        with summary_path.open("r", encoding="utf-8") as f:
            rows = list(csv.DictReader(f))
        if len(rows) < 10:
            print(f"[WARN] benchmark_summary.csv has only {len(rows)} rows (expected ~12).")
        else:
            print(f"[PASS] benchmark_summary.csv loaded ({len(rows)} rows).")
    except Exception as e:
        print(f"[FAIL] Could not read benchmark_summary.csv :: {e}")
        ok = False

    # 5) Per-chemical dossier completeness
    pub = root / "publication_artifacts"
    skip = {"figures", "pbpk_outputs", "pbpk_server_var"}
    chemicals = [d for d in pub.iterdir() if d.is_dir() and d.name not in skip]
    if not chemicals:
        print("[FAIL] No chemical folders found in publication_artifacts/")
        ok = False
    else:
        for chem in chemicals:
            missing = [f for f in REQUIRED_PER_CHEM if not (chem / f).exists()]
            aop_present = any((chem / f).exists() for f in AOP_OK)
            if not aop_present:
                missing.append("aop_map.json or aop_map_empty.json")
            if missing:
                print(f"[FAIL] {chem.name}: missing {missing}")
                ok = False
        if ok:
            print(f"[PASS] All {len(chemicals)} chemical dossiers contain required artifacts.")

    if ok:
        print("\nVALIDATION: PASS ✅")
        return 0
    print("\nVALIDATION: FAIL ❌")
    return 2


if __name__ == "__main__":
    raise SystemExit(main())
```
