## Supplementary figures and images for "ToxMCP: Guardrailed, Auditable Agentic Workflows for Computational Toxicology via the Model Context Protocol"

### Figure_1_PBPK_Sensitivity.png

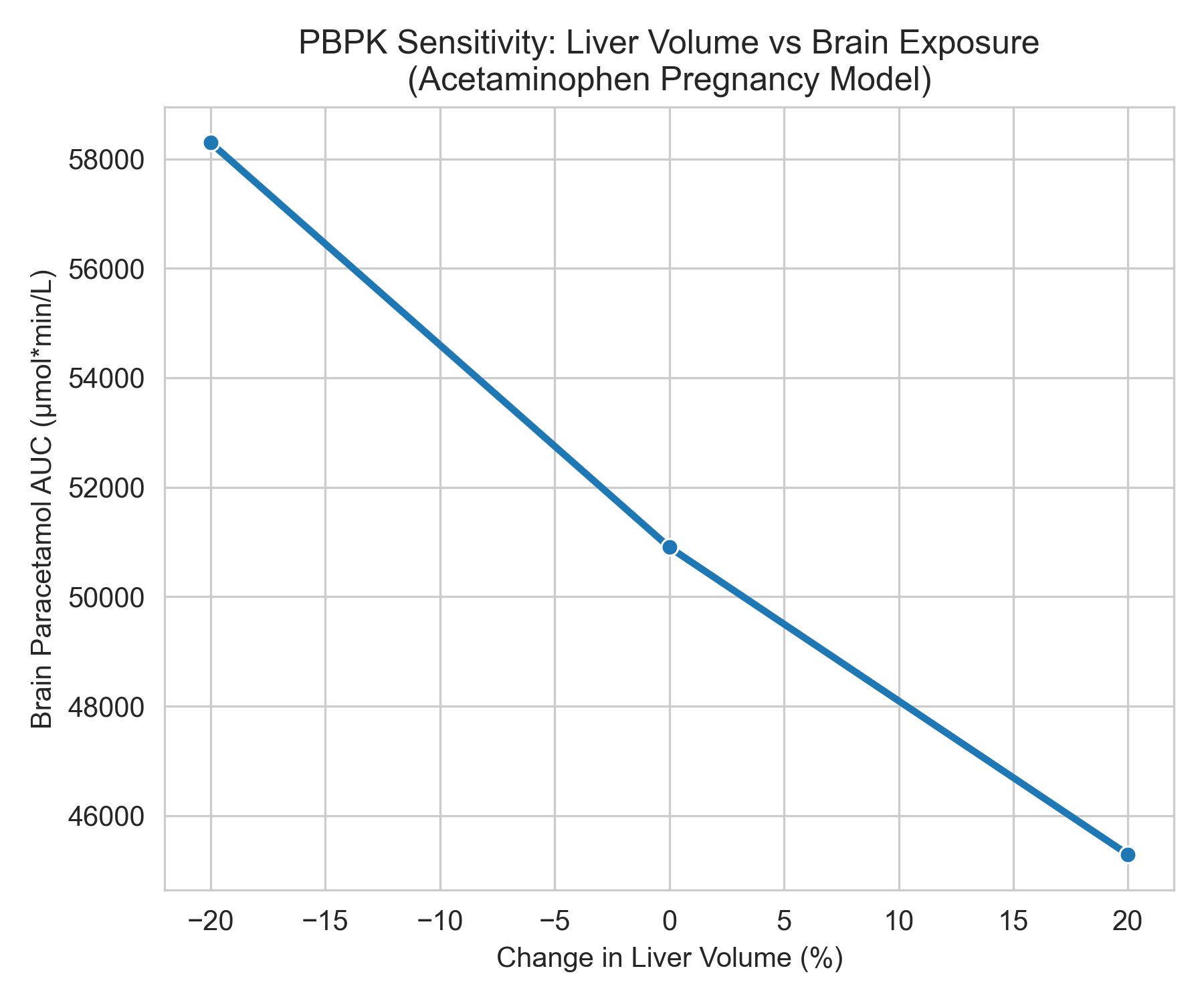

### Figure_2_Data_Completeness.png

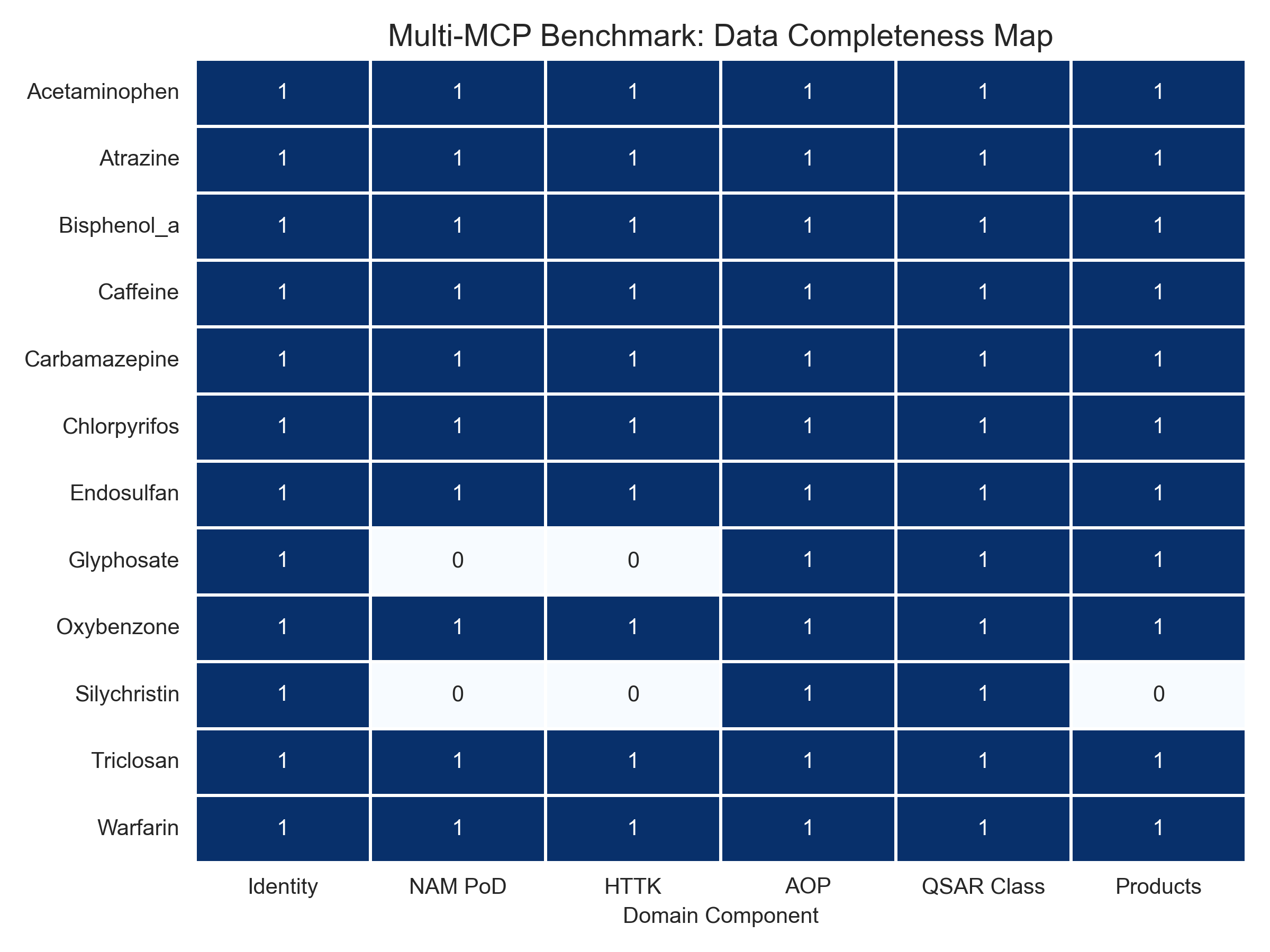

### Figure_3_PoD_vs_HTTK.png

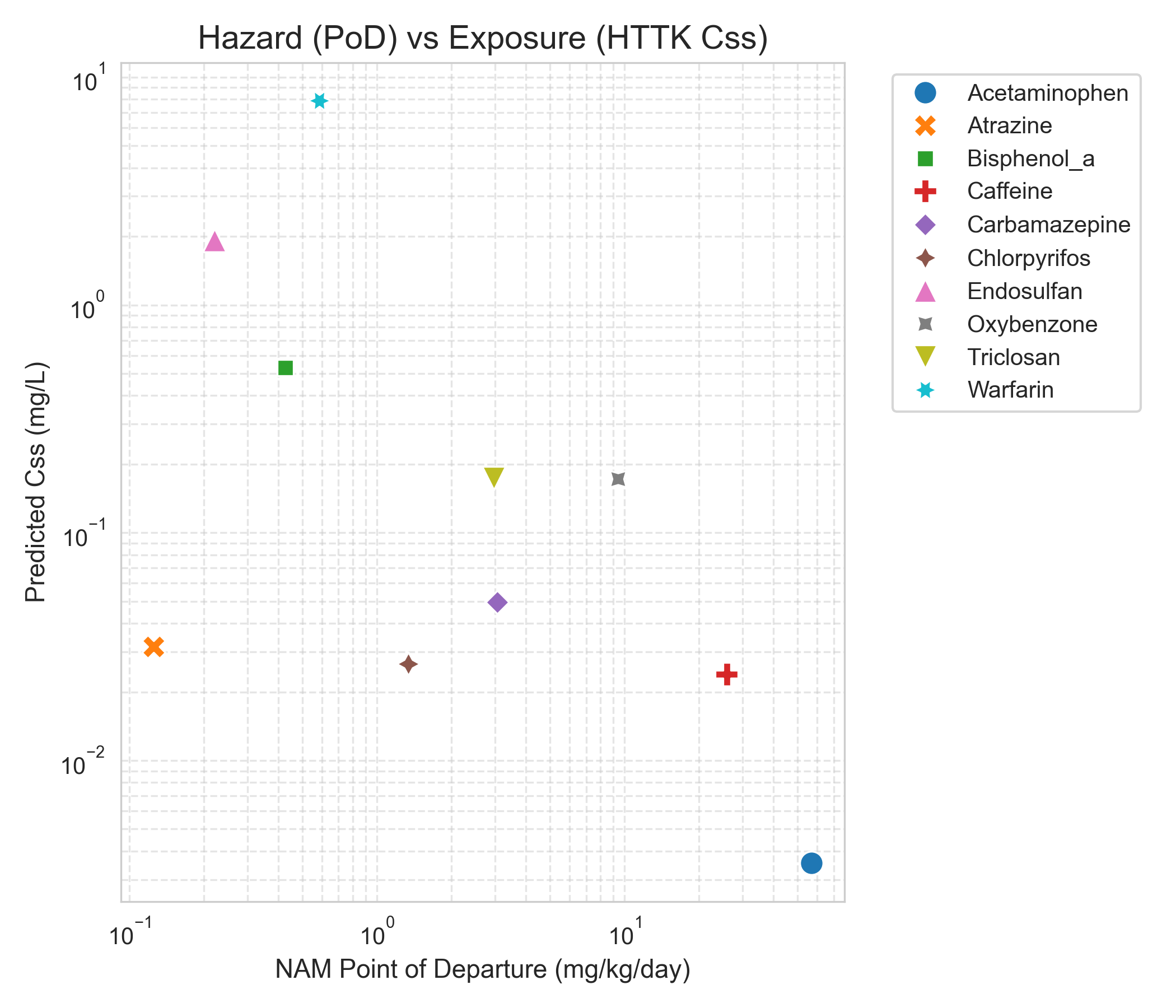

### Figure_4_MultiMCP_Dossier_Summary.png

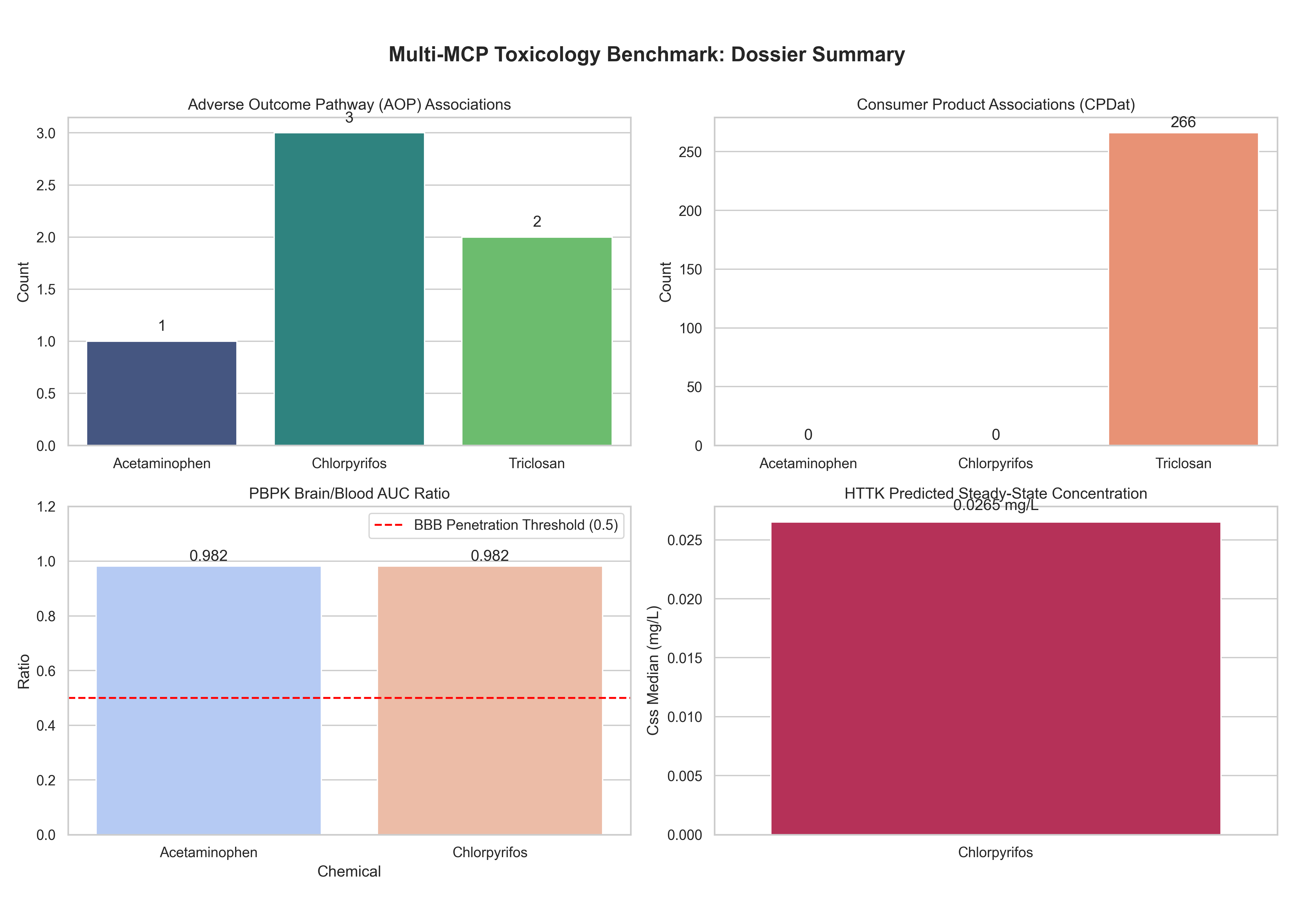

### Figure_5_OQT_Capabilities.png

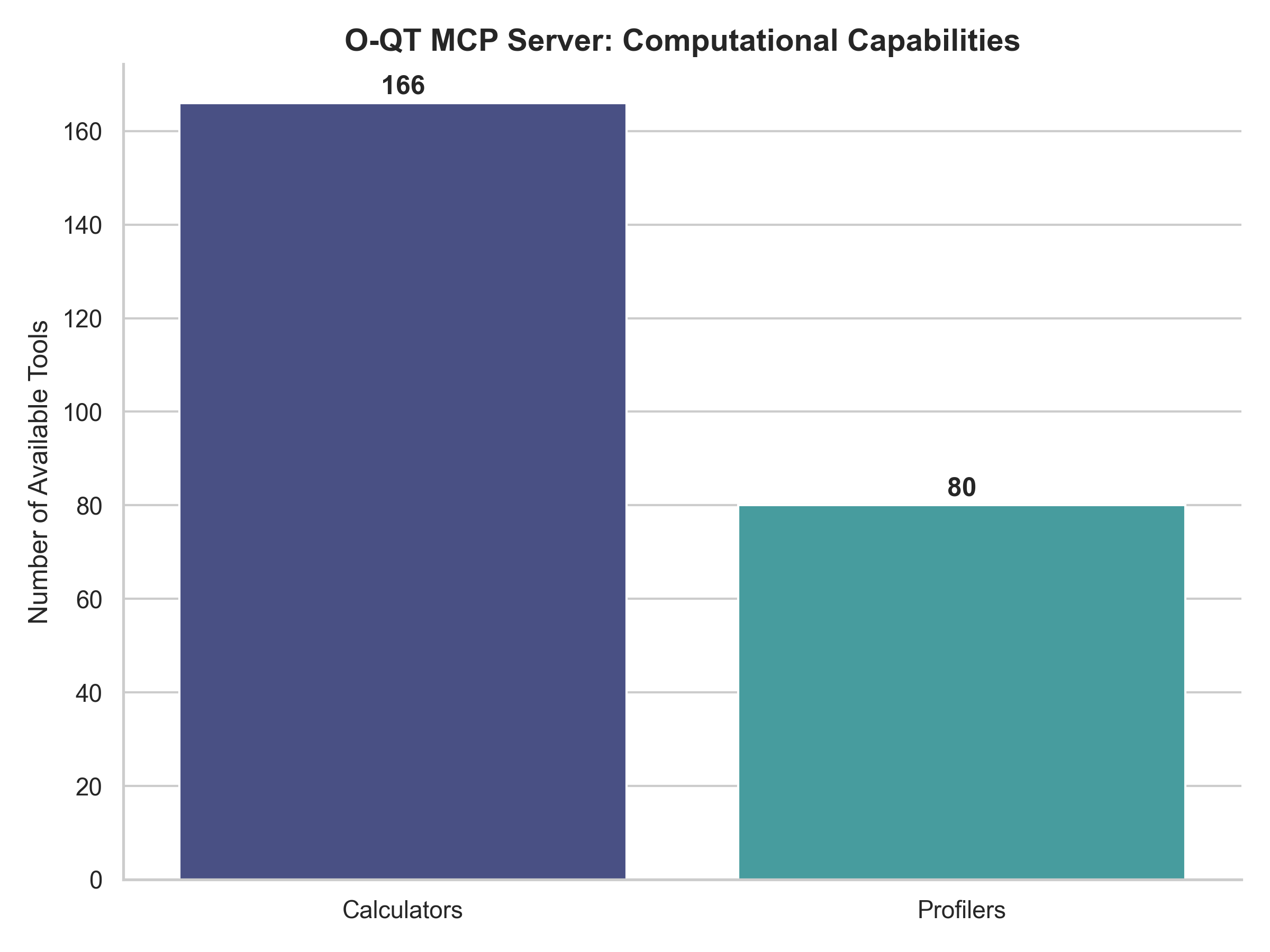

### Figure_5_PBPK_Determinism_Proof.png

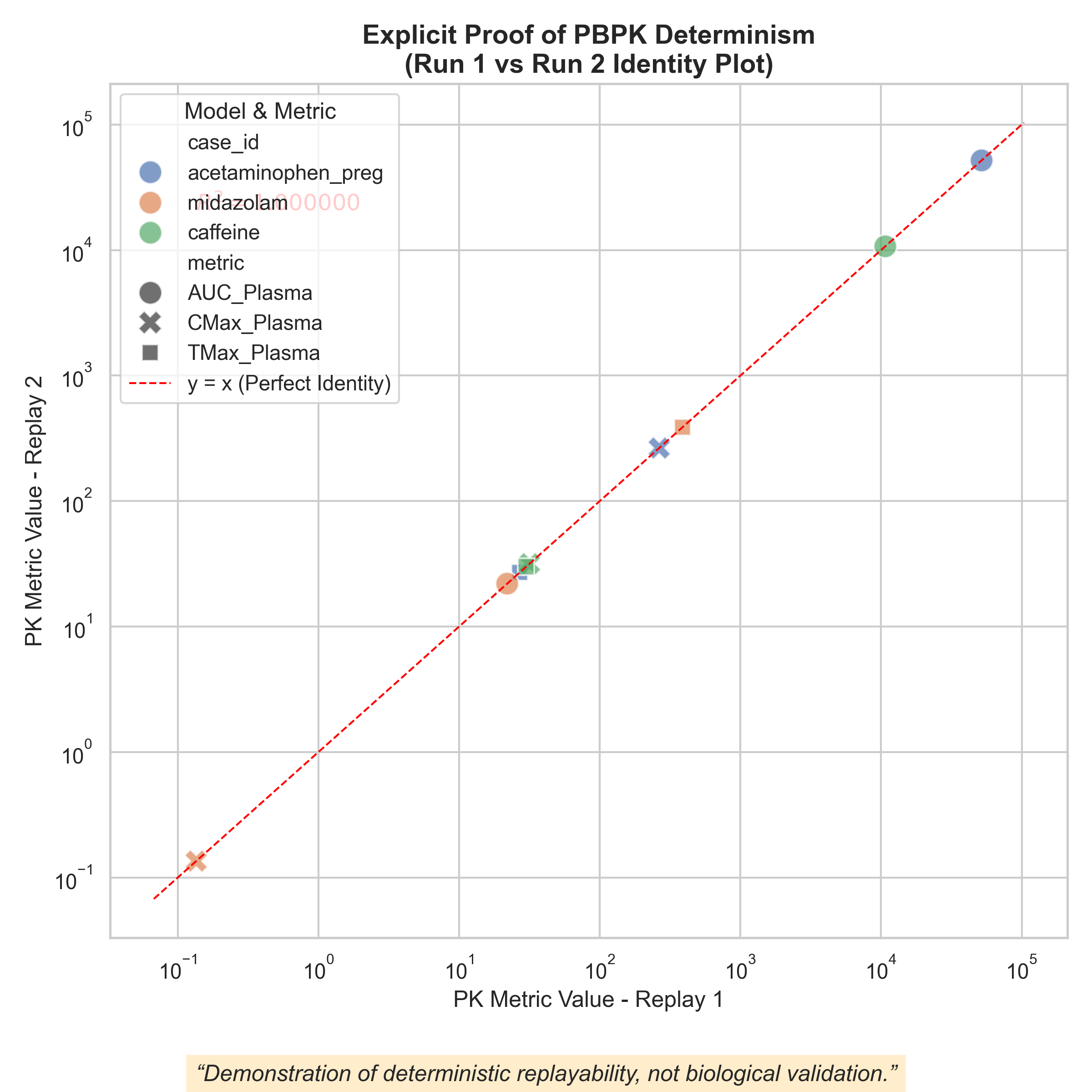

### Figure_6_PBPK_Sensitivity_Curve.png

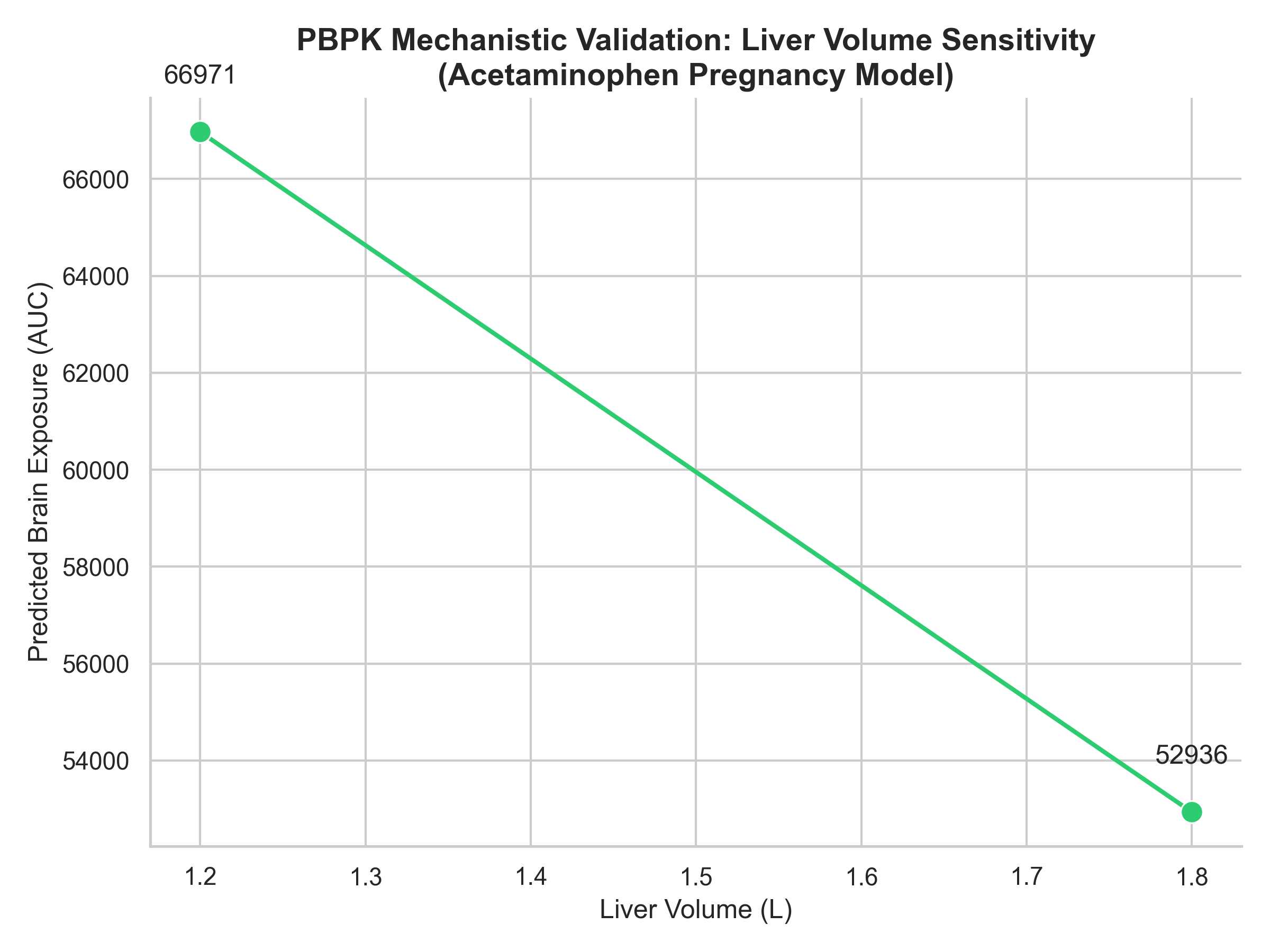

### Figure_7_Triclosan_Product_Landscape.png

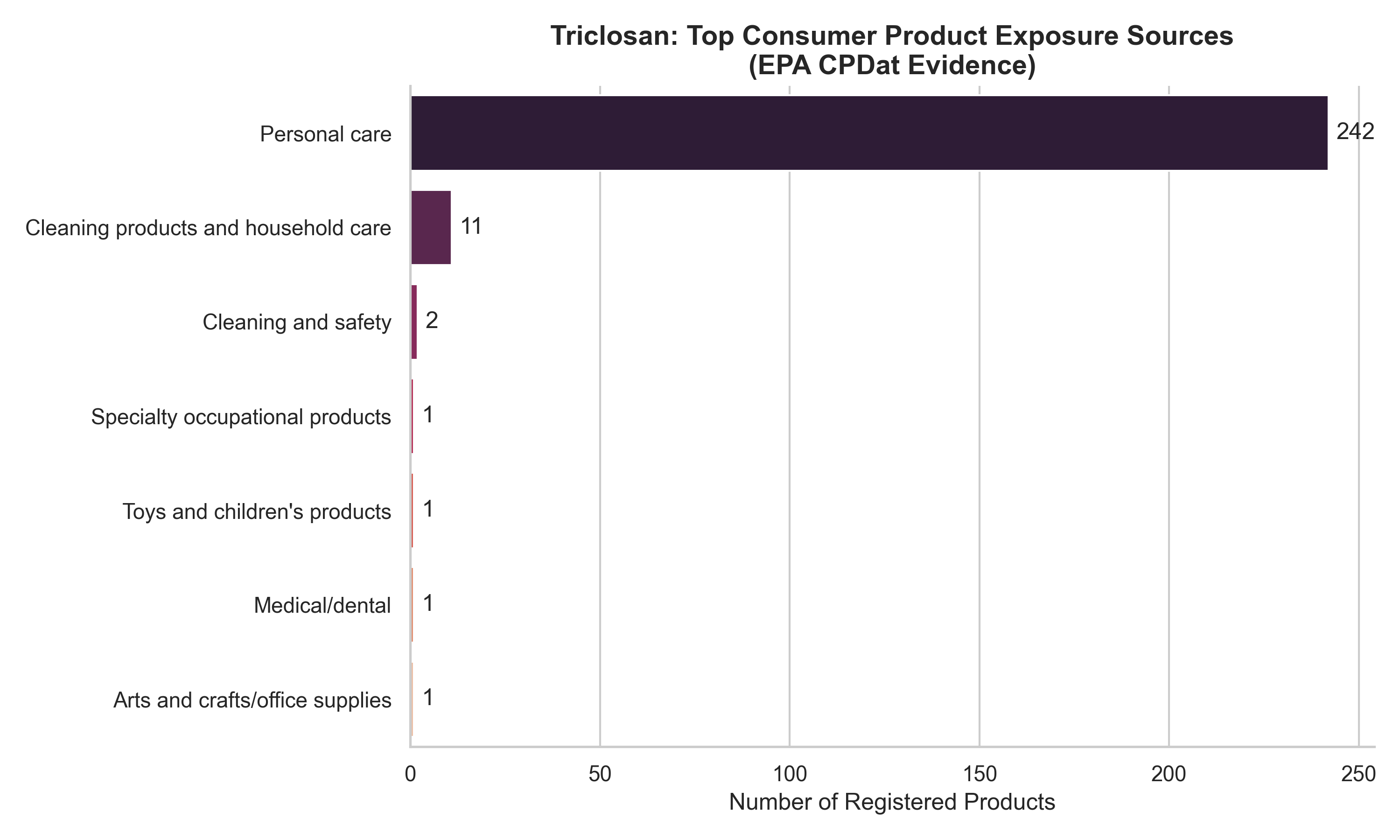

### Figure_8_HTTK_Kinetic_Uncertainty.png

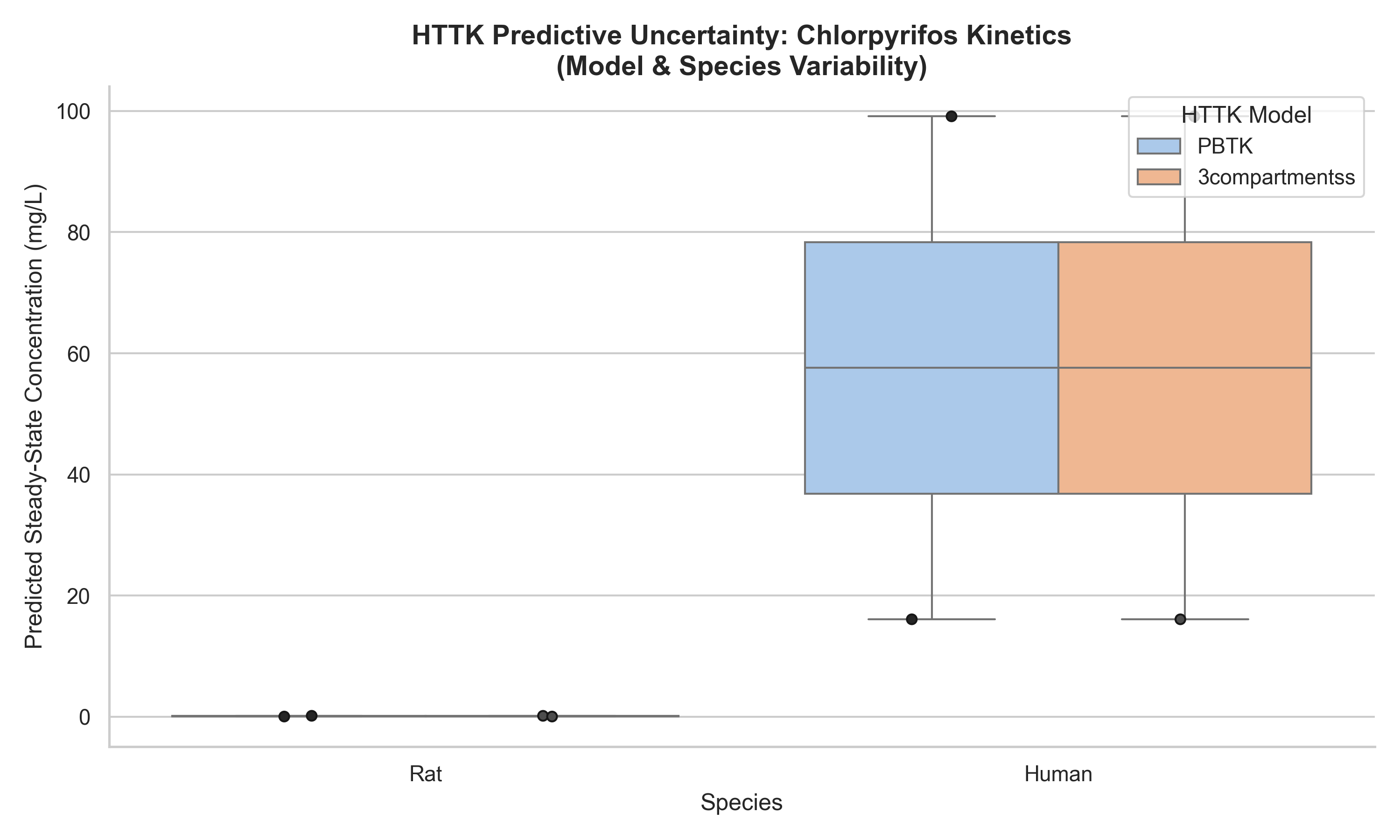

### Figure_9_NAM_PoD_Distribution.png

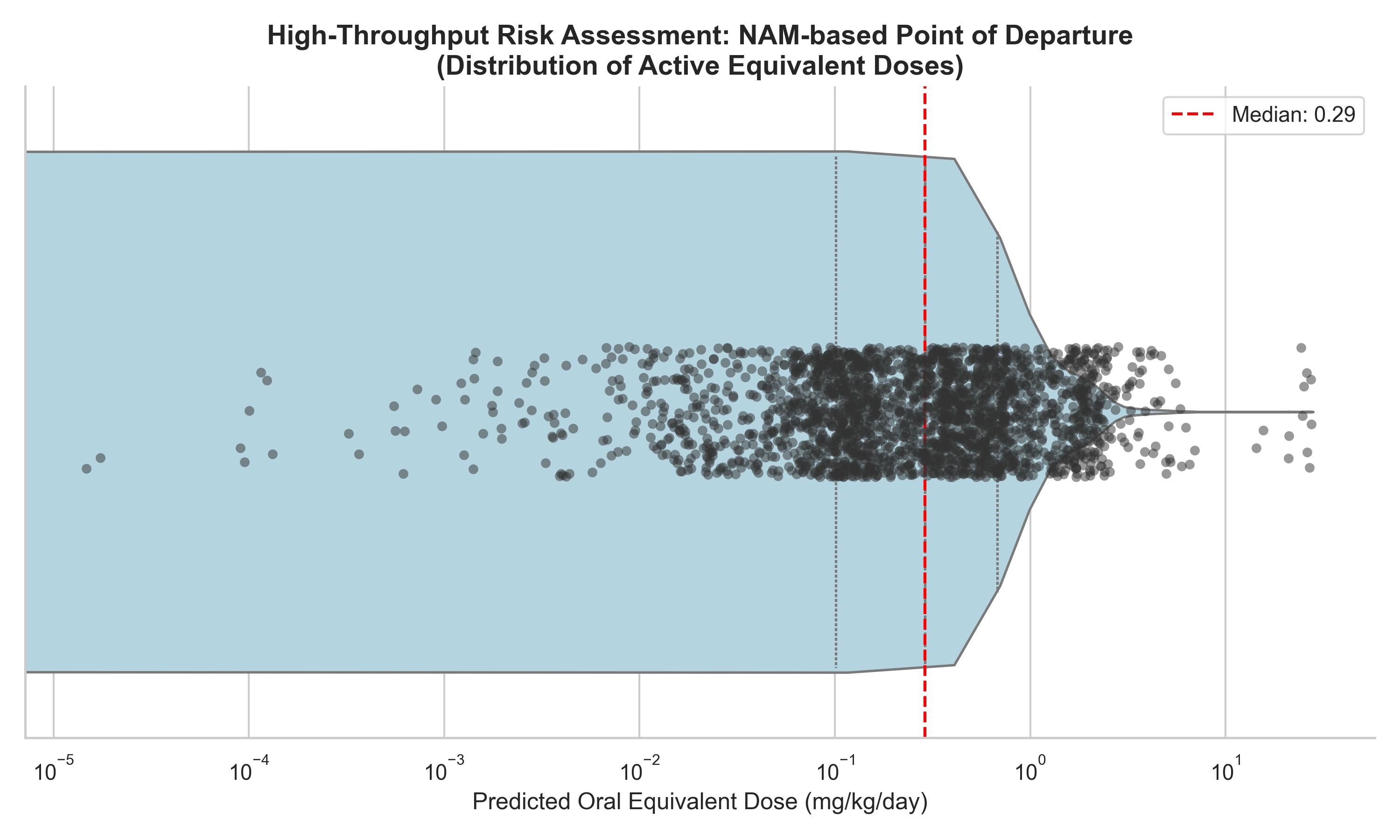

### Figure_10_Metabolic_Fate.png

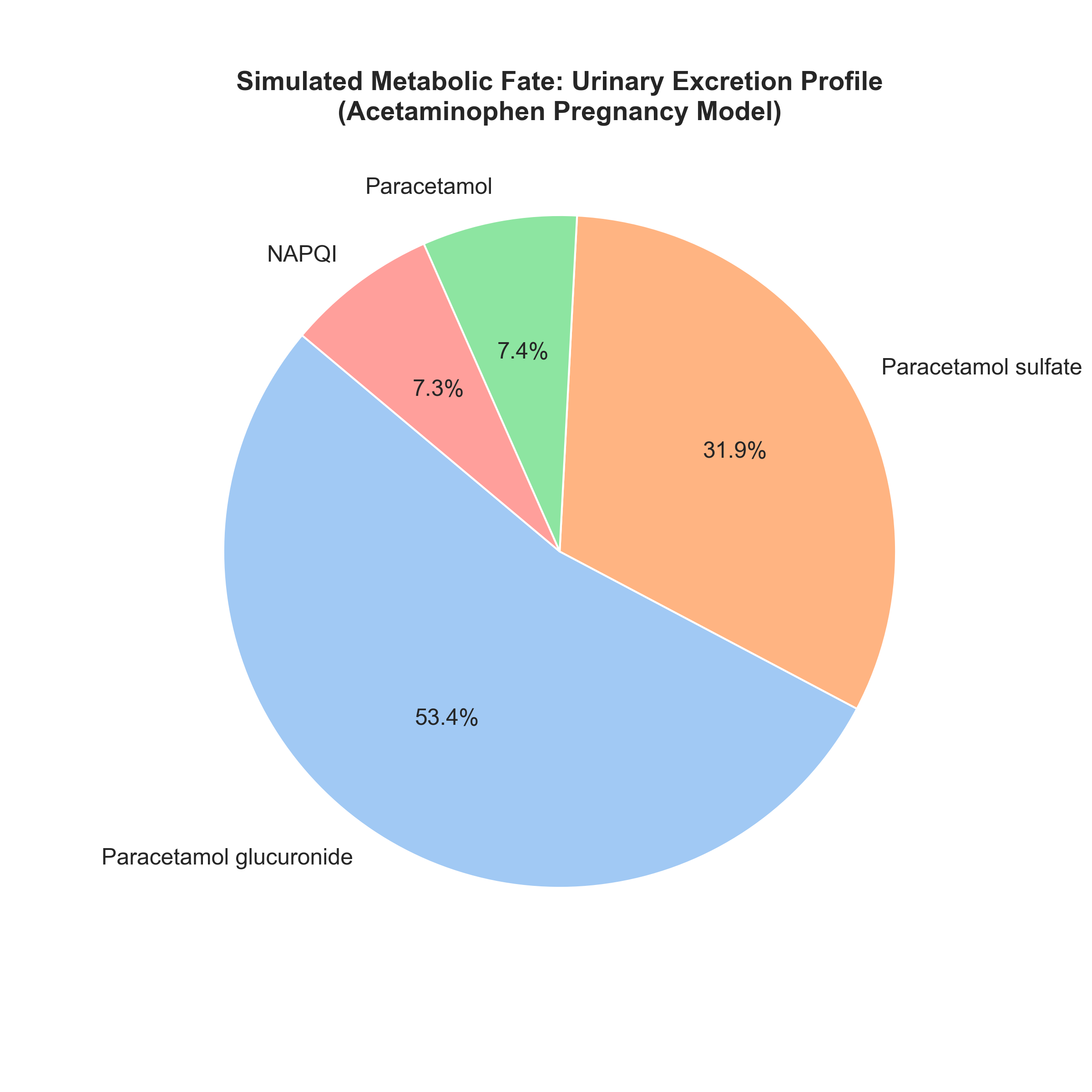

### Figure_11_AOP_Network.png

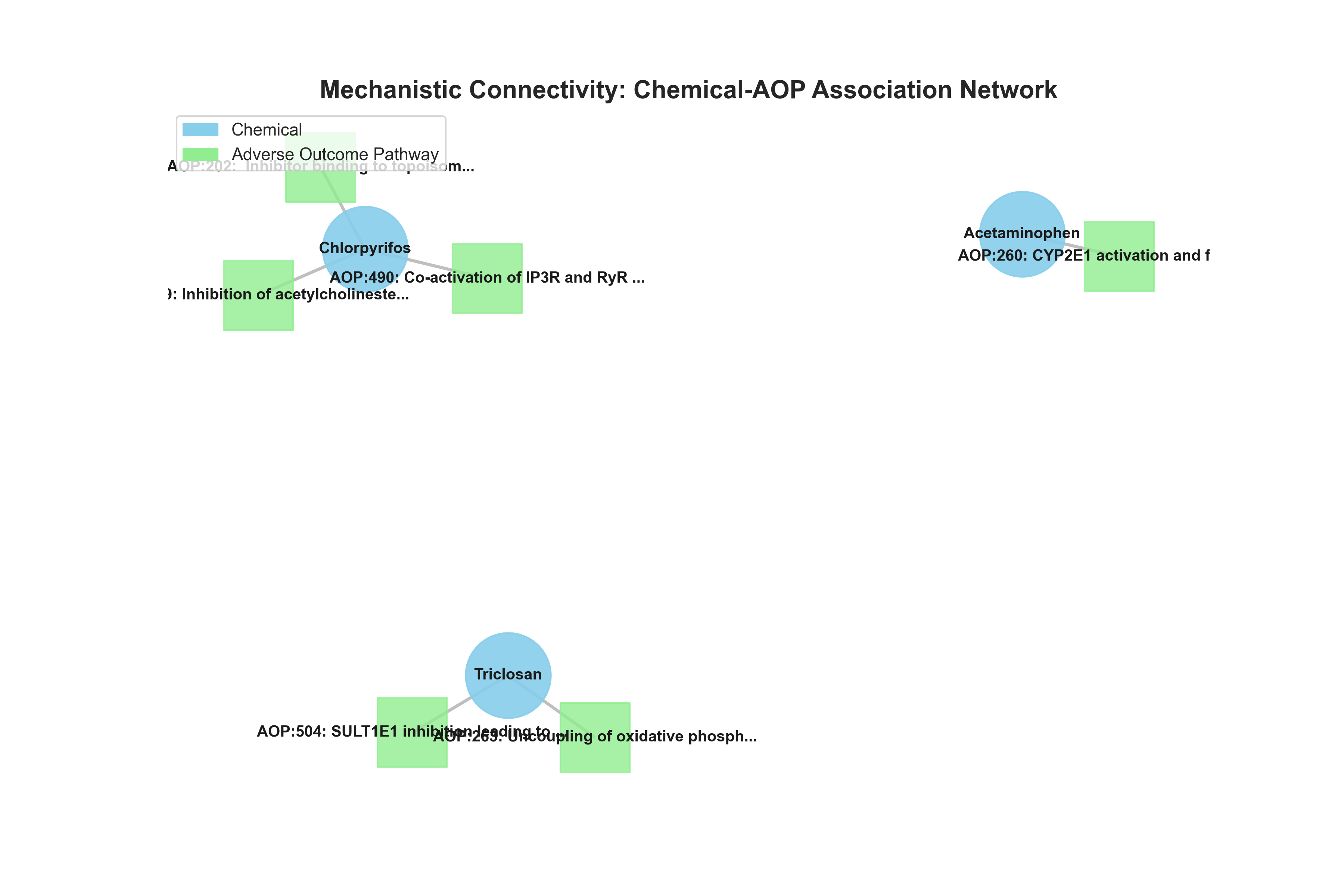

### Figure_12_AOP_559_Pathway.png

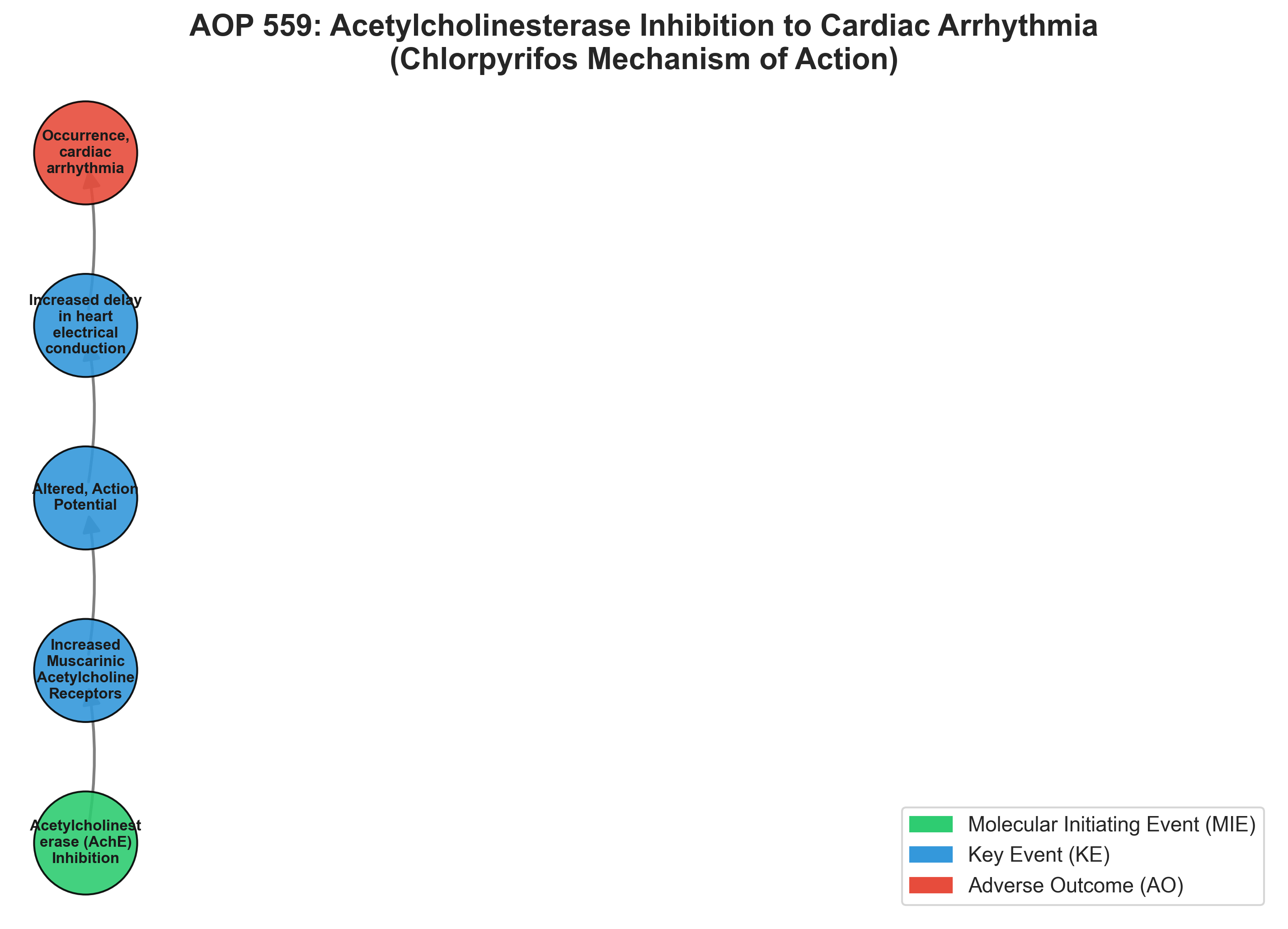
